# Genome organization by SATB1 binding to base-unpairing regions (BURs) provides a scaffold for SATB1-regulated gene expression

**DOI:** 10.1101/2021.12.19.473323

**Authors:** Yoshinori Kohwi, Xianrong Wong, Mari Grange, Thomas Sexton, Hunter W. Richards, Yohko Kitagawa, Shimon Sakaguchi, Ya-Chen Liang, Cheng-Ming Chuong, Vladimir A. Botchkarev, Ichiro Taniuchi, Karen L. Reddy, Terumi Kohwi-Shigematsu

## Abstract

Mammalian genomes are organized by multi-level folding, yet how this organization contributes to cell type-specific transcription remain unclear. A nuclear protein SATB1 forms a SATB1-rich subnuclear structure that resists high-salt extraction. SATB1 binds in the minor groove of double-stranded base-unpairing regions (BURs), genomic elements with high unwinding propensities. We uncovered that SATB1 establishes a two-tiered chromatin organization, one through indirect binding and another by direct binding of BURs. Published ChIP-seq datasets show SATB1 binding to highly accessible chromatin at enhancers and CTCF sites, but not to BURs. By employing urea ChIP-seq, which retains only directly bound protein:DNA complexes, we found that BURs, but not CTCF sites, are direct SATB1 binding targets genome-wide. BURs bound to the SATB1 nuclear substructure interact with accessible chromatin crossing multiple topologically associated domains (TADs). SATB1 is required for these megabase-scale interactions linked to cell type-specific gene expression. BURs are highly enriched within lamina associated domains (LADs), but some (∼10%) are found in gene-rich accessible chromatin outside LADs as well. Only a subset of BURs is bound to SATB1 depending on cell type. Notably, despite the mutually exclusive SATB1-binding profiles uncovered by the two ChIP-seq methods, we found most peaks in both profiles are valid and require SATB1. Based on these and previous data, we propose that the SATB1 protein network forms a chromatin scaffold, providing an interface that connects accessible chromatin to a subnuclear architectural structure, thereby facilitating the three-dimensional organization linked to cell type-specific gene expression.

## Introduction

Three-dimensional (3D) chromatin architecture, formed by chromatin folding at multiple hierarchical levels, is thought to play a fundamental role in genome activity such as gene expression (*1-7*). Whether and how genome folding contributes to gene regulation, however, remains largely unknown. High resolution chromatin interaction maps, generated by chromosome conformation capture methods such as Hi-C (all-to-all contacts), have revealed chromatin to be partitioned mainly into two types of megabase (Mb)-sized compartments, A and B (*8*). The euchromatic A compartment is gene-rich, transcriptionally active and enriched in histone marks associated with active chromatin, conferring high chromatin accessibility. On the other hand, the B compartment is gene-poor, transcriptionally repressive, and enriched in repressive histone marks. The largely transcriptionally repressive lamina-associated domains (LADs) are among the B compartment chromatin displaying the typical features of heterochromatin (*9, 10*). LADs have been identified by the lamin B1-mediated adenine methyltransferase identification (DamID) method (*11*) as genomic regions that are spatially proximal to the nuclear lamina, an intermediate filament meshwork adjacent to the inner nuclear membrane. LADs represent discrete chromatin domains of 40kb to 30 Mb in size (*9*). Approximately 1000-1500 LADs have been identified in mouse and human cells and, in a given cell type, typically cover ∼30% of the genome (*2*). By sequestering chromatin to the repressive nuclear compartment, the nuclear lamina is thought to play an important role in genome organization.

At sub-megabase scales, using Hi-C, chromatin domains called “topological-associated domains” (TADs) were previously detected that are relatively insulated from adjacent TADs by preferentially making intra-domain contacts via looping (*12-14*). CTCF and cohesin are the two major architectural proteins required for folding chromatin into loops. The loop extrusion model postulates that cohesin functions in loop extrusion until it encounters the CCCTC-binding factor (CTCF) bound to its target sites in a convergent orientation (*15, 16*). CTCF- and cohesin-binding sites are enriched at the boundaries of TADs (*12, 17*) and LADs (*9*). Acute depletion of CTCF (*18*), cohesin (*19, 20*) or induced removal of cohesin loading factor NIPBL (*21*) results in a major loss of TADs. Unexpectedly, the majority of protein-coding genes remained largely unchanged in their expression levels in the absence of TADs. These results challenge the view that TADs provide a unit of gene expression control by insulating enhancer-promoter interactions from neighboring TADs. In the last few years, using super-resolution live-cell imaging, genome interactions were studied at the single cell levels. These studies revealed that chromatin looping formed by CTCF and cohesin are highly dynamic and variable among individual single cells, suggesting that TADs are an emergent property of the cell population average of heterogeneous chromatin folding (*22-26*). Therefore, TADs, enriched in contacts, may not represent a stable structural unit of chromatin flanked by two boundary loci stably bound by CTCF and cohesin [reviewed in (*25, 27, 28*)]. Super-resolution microscopy detected not only single cohesin-mediated loops but also radially stacked loops forming “rosette” like structures in individual single cells. These loops were also transient and a single cell spends a minority of its time in the configuration, and depletion of cohesin led to increased variability in gene expression at single cell levels (*29*). As such, although research on CTCF and cohesin has greatly advanced our understanding on chromatin loop formation, there is still much to learn about how and what functional 3D chromatin structure underlies gene activity, especially cell-type specific gene expression. It is increasingly clear that it is necessary to explore additional nuclear factors that potentially drive chromatin folding. In this paper, we studied function of a tissue-specific architectural protein SATB1 in genome organization.

SATB1 was originally identified by virtue of its specific binding to genomic sequences called Base Unpairing Regions (BURs). Previous *in vitro* studies have shown that SATB1 directly binds BURs without requiring additional molecules, such as RNA or proteins (*30-32*). BURs have a strong unwinding property under negative superhelical stress, and BURs are empirically identified using unpaired DNA-specific chemical probes (*30, 33*) because they lack primary sequence consensus. However, BURs have a unique sequence context: a typical BUR (∼200-300bp) contains a cluster of short sequence segments (25-50 bp/segment) containing exclusively As, Ts, and Cs on one strand (ATC context) (*30, 32*). BURs are at least 65% AT-rich, but importantly, not all AT-rich sequences are BURs. Importantly, SATB1 binds to BURs in the minor groove of double-stranded DNA, recognizing the altered phosphate backbone structure; it does not bind to single-stranded DNA (*32*). Disruption of the ATC context by mutagenesis, without affecting the AT content, results in loss of unwinding property of a BUR and SATB1 binding. While BURs are obviously present in the genome across cell types, SATB1 protein is cell-type restricted, most abundantly expressed in thymocytes and among multiple adult progenitor cells (*34-36*), as well as terminal differentiated postnatal neurons in the cortex and amygdala (*37, 38*). Located in nuclear interior, SATB1 has an interconnected cage-like distribution in thymocytes (*39*) and a fine spider web-like distribution in neurons (*37*) forming a unique SATB1 subnuclear structures that are resistant to high-salt extraction. Some BURs have been identified and characterized as *in vivo* SATB1 binding sites and, when bound to SATB1, are also resistant to salt extraction (*39, 40*). SATB1 has been shown to be essential for loop formation connecting a subset of these BURs and recruiting chromatin remodeling complexes, epigenetic factors and transcription factors to specific gene loci (*41*) Of particular interest, SATB1 regulates expression of hundreds of protein-coding genes in a cell-type specific manner to enable cells to acquire new phenotypes, such as during T cell activation (*42*), development of many adult progenitor cells (*34–37, 43-49*), and cancer metastasis (*41, 50-52*). For example, by directing lineage-specific transcriptional programs, SATB1 plays an essential role in development of CD4^+^ T cells, CD8^+^ T cells, NKT cells, and Foxp3^+^ regulatory T cells (Treg) in the thymus(*46, 47*) as well as in promoting the pathogenic effector program of tissue Th17 cells in autoimmune disease in mice(*48*). These activities have been shown to mediate co-induction of interleukin genes in T helper cells (*42*) and regulate ∼1000 genes in cancer to promote metastasis(*50*). Thus, SATB1 has long been thought to play a role in regulating genome architecture and in gene expression.

Although BURs have been identified to be specific and direct binding target sequences of SATB1, mysteriously the genome-wide SATB1 binding profiles obtained by the widely used, standard chromatin immunoprecipitation-deep sequencing (ChIP-seq) does not detect SATB1 interaction with BURs. Instead, SATB1 has been mapped mainly to enhancers and to a region that would become an active super-enhancer upon differentiation (*45-47*). In line with this, recent studies on chromatin looping with Hi-C also shows that SATB1 mediates interactions involving a subset of enhancers and promoters for cell identity gene expression and that SATB1 binding sites coincide with many CTCF-bound loci (*53-55*). These results align with SATB1’s role in assembling transcription complexes at regulatory regions. However, they are also puzzling, considering that BURs have been identified as the direct binding targets of SATB1 by *in vitro* binding assays. We therefore hypothesized that SATB1-BUR interactions are somehow undetectable by the standard ChIP-seq approach and CUT&Tag methods, and if identified, would provide additional mechanisms for chromatin looping events *in vivo* that link to gene expression. ChIP-seq has been a gold standard for profiling chromatin-protein interactions. However, standard ChIP-seq approaches have several drawbacks. First, chromatin structure can introduce biases into ChIP-seq analysis. Fragmentation of crosslinked chromatin in whole cells/nuclei can result in biased capture of transcriptionally active, highly accessible “open” chromatin and generation of false-positive “phantom” peaks (*56-60*). Second, it fails to distinguish between direct and indirect (piggybacking) chromatin binding of a protein of interest. Here, we optimized urea-ChIP-seq, a modified ChIP-seq that first isolate stringently purified whole intact genomic DNA that retains only its directly bound proteins from crosslinked cells prior to fragmentation and immunoprecipitation. This captures the entire genome, regardless of its original chromatin accessibility status. This approach has allowed us to identify genomic sites directly bound by SATB1 and revealed BURs to be the primary *direct* SATB1 target sites, previously hidden by standard ChIP-seq. The SATB1-bound BURs are mutually exclusive with enhancer-enriched SATB1-bound sites mapped by standard ChIP-seq. Instead, a majority of BURs are mapped within LADs, and depending on cell type, SATB1 binds *in vivo* to selective subsets of these BURs. Using the urea 4C-seq (one-to-all contacts), select SATB1-bound BURs were found to interact extensively over the >5.7Mb gene-rich region within DNase 1 hypersensitive (DNase 1 HS) regions, but not with adjacent gene-poor regions. SATB1 is essential for both megabase-level chromatin interactions and proper expression of multiple genes in this gene-rich region. Collectively, our results suggest that SATB1-mediated genome organization exhibits two distinct features, a “skeletal framework” where SATB1 binds BURs directly, and a network of gene-rich open chromatin that is tethered to BURs indirectly through SATB1-mediated chromatin looping, which is correlated to gene expression.

## Results

### Development of urea ChIP-seq method to detect genome-wide SATB1-binding profiles

Unlike many nuclear proteins (e.g. transcription and chromatin modifying factors), SATB1 resides in nuclear substructures exhibiting resistance to extraction with high salt or lithium 3,5-diidosalicylate (LIS) (*40, 44*). Consistent with this, after extraction of nuclei with buffer containing 2M NaCl that removed most of the proteins from DNA, individually cloned SATB1-bound BURs remained anchored to the residual SATB1 protein network (SATB1-rich subnuclear architecture) found in the interior of nuclei (*39*). By contrast, in high-salt extracted *Satb1*^−/−^ (SATB1-KO) thymocyte nuclei, these BURs lost their anchored sites and were exclusively found in the distended DNA halos that spread around the residual nuclei, which is illustrated (*Figure 1A*) (*39*). Thus, we hypothesized that SATB1-BUR interactions are likely embedded in an insoluble subnuclear structure that could not be detected to date by the widely used standard ChIP-seq methods. It is possible that ChIP-seq experiments have inherent biases arising from chromatin structure such as heterochromatin and transcriptionally active regions (*61*).

**Figure 1.**
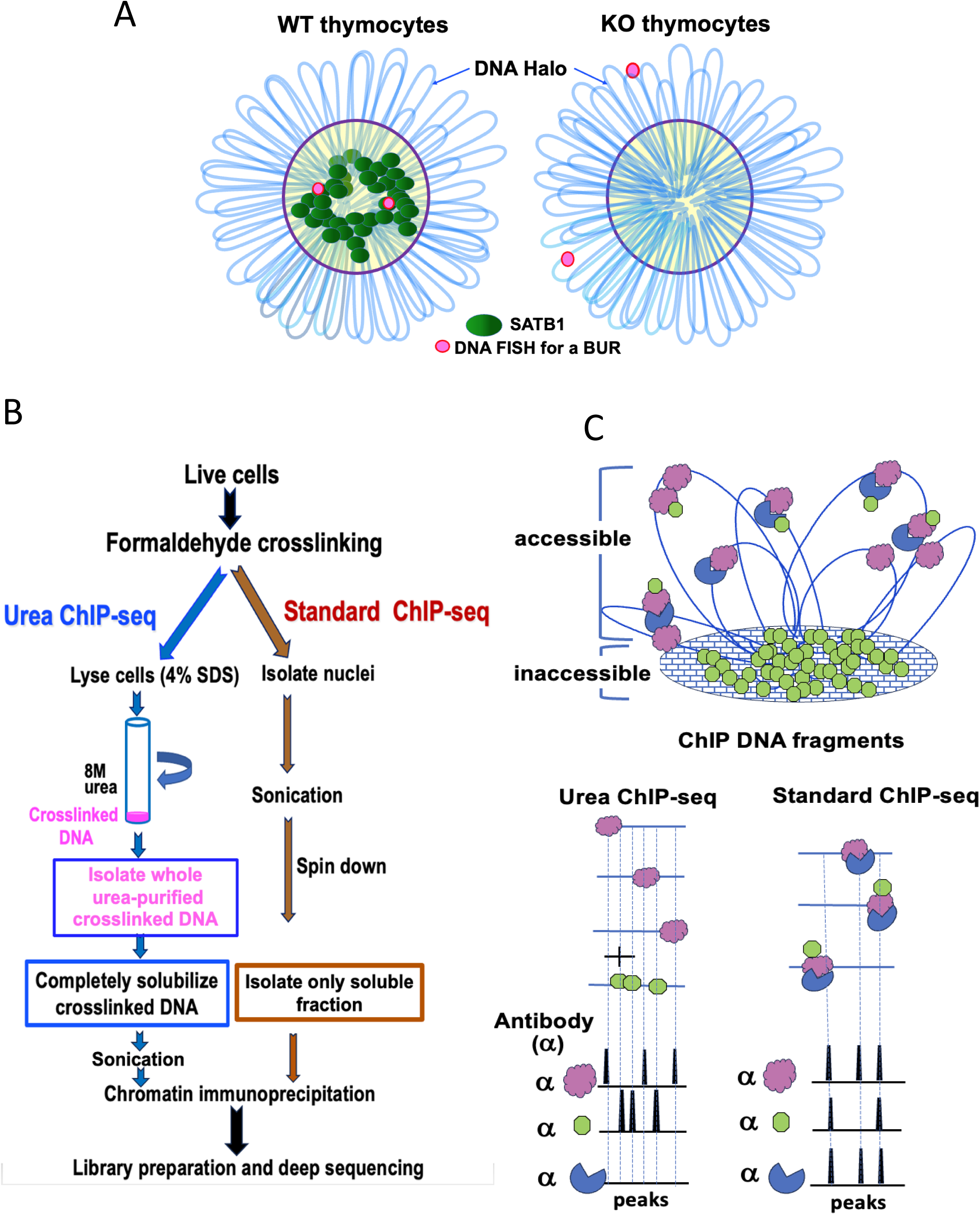
Scheme of urea ChIP-seq and difference from standard ChIP-seq. A) An illustration for a BUR anchored to the high-salt extraction resistant SATB1 nuclear architecture (*39*). DNA-FISH signals (red dots) for a cloned SATB1-bound DNA in both alleles are shown in *Satb1*^+/+^(WT) and *Satb1*^−/−^(KO) thymocytes on slides treated with 2M NaCl solution to produce DNA halos around the nuclei (blue loops). SATB1 and the SATB1-bound DNA remain in the residual nucleus in WT thymocytes after salt treatment. However, in KO thymocytes the DNA regions normally bound by SATB1 are located in the DNA halos after the treatment. B) A comparative overview of the urea ChIP-seq and standard ChIP-seq protocols. In urea ChIP-seq, entire crosslinked chromatin purified by 8M urea ultracentrifugation is solubilized (see Materials and Methods) before chromatin immunoprecipitation. By contrast, in standard ChIP-seq, crosslinked chromatin in whole nuclei is sonicated, centrifuged and the soluble fraction is isolated for chromatin immunoprecipitation. These critical steps that are different between urea and standard ChIP-seq are highlighted in boxed texts. C) Differences in the profiles of the peaks generated from urea ChIP-seq versus standard ChIP-seq. A diagram is shown for ChIP-seq peaks using antibodies (α) against a DNA direct-binding protein (pink), a DNA indirect-binding protein (blue), and DNA direct-binding protein that resides in the inaccessible chromatin region (green) Note: This diagram illustrate a hypothetical scenario where some of the proteins (green) in the inaccessible chromatin region may also reside in the accessible chromatin binding indirectly to chromatin.

To detect BURs as the primary targets of SATB1 genome wide, we devised a urea ChIP-seq method to purify genomic DNA with only its directly bound proteins. Through this apporach, we found that a vast majority of *in vivo* SATB1-binding sites are BURs in the genome. In this method, cells are first crosslinked with formaldehyde. Then, cells are lysed in a 4% SDS-containing buffer and subjected to ultracentrifugation with 8M urea. In this manner, crosslinked chromatin is stringently purified by removing all free-floating proteins and proteins bound to DNA indirectly. Therefore, urea ChIP-seq is performed with the genomic DNA retaining only directly bound proteins. It is important to note that without prior formaldehyde crosslinking, urea centrifugation removes 99.9% of chromatin-associated protein from chromatin and sediments unsheared pure genomic DNA at the bottom of the tube (*62*). As a denaturant for DNA, RNA, and proteins, high concentrations of urea (e.g. 6-8M) disrupt the secondary structures of nucleic acids and the three-dimensional structure of proteins (*63-65*). Crucially, purified genomic DNA obtained from 8M urea ultracentrifugation remains double-stranded and can therefore be readily digested with restriction enzymes (this study and (*66*). Additionally, the proteins that remain bound to urea-purified DNA from crosslinked cells can still be detected with the antibodies compatible with standard ChIP protocols. By acting as both a hydrogen bond donor and acceptor, high concentrations of urea destabilize double-stranded DNA by weakening the hydrogen bonds between base pairs, thereby decreasing the melting temperature of DNA (*67, 68*). However, even 6-8 M urea alone, without heat treatment, is insufficient to convert double-stranded DNA into single-stranded DNA (*69*). Denaturing polyacrylamide electrophoresis for separating DNA oligonucleotides by size typically uses 6-8 M urea. For this, DNA samples need to be heat denatured first to make them single-stranded (*70*). Our 8 M urea centrifugation protocol lacks heat treatment and includes prior formaldehyde fixation, thus allowing genomic DNA to remain double-stranded and bound by interacting proteins

Critically, the urea ChIP-seq method (*66*) allows detection of genomic regions solely based on their direct protein binding regardless of their original chromatin status, such as ‘accessible’ or ‘inaccessible’ chromatin. To obtain reproducible direct binding profiles of SATB1 at the genome-wide scale, it was necessary to make additional optimizations. The modified urea ChIP-seq protocol allows quantitative solubilization of the unsheared crosslinked genomic DNA fiber after urea ultracentrifugation, while still maintaining intact chromatin-protein interactions during fragmentation. The critical steps that are different between urea and standard ChIP-seq are highlighted (*Figure 1B*). A putative scenario as to how DNA-binding sites of chromatin-associated factors are captured by the two different ChIP-seq methods is illustrated (*Figure 1C*). Urea ChIP-seq allows detection of chromatin-bound proteins regardless of the chromatin’s accessibility because the whole genome is purified. In contrast, standard ChIP-seq discards most of the insoluble fraction and retains primarily the soluble fraction for the subsequent chromatin immunoprecipitation step (*61*). This results in samples that are enriched in accessible “open” chromatin. Because a fraction of SATB1 proteins can be easily extracted from cells with a high salt-containing buffer whereas other fraction resists extraction, both soluble and insoluble fractions are expected to contain SATB1(*40*). This could reflect SATB1 proteins being heterogeneous with regards to their mobility. By fluorescence recovery after photobleaching (FRAP) assay in thymocytes, we found one fraction of SATB1 essentially immobile and the other with high mobility (our unpublished results). Due to the lack of specific markers to distinguish these SATB1 proteins, we cannot confirm the exact relationship between fast-mobile SATB1 and immobile SATB1 to the soluble and insoluble fractions, respectively. Comparing the two ChIP-seq protocols, standard ChIP-seq, therefore, captures both directly and indirectly bound proteins for chromatin fragments in the soluble fraction, and their binding profiles cannot be distinguished. In contrast, urea ChIP-seq strongly enriches for directly bound proteins, including those in chromatin regions that are difficult to solubilize. Thus, urea ChIP-seq is uniquely suited for detecting direct DNA binding profiles of SATB1 that resides in the high-salt extraction resistant subnuclear structure.

### Urea ChIP-seq detects BURs as direct binding targets of SATB1 *in vivo*

To determine whether SATB1 binds to BURs *in vivo*, it is essential to validate *in vivo* SATB1-bound sites as BURs. We previously generated a genome-wide BUR reference map by deep sequencing DNA fragments generated from purified genomic DNA bound by recombinant SATB1 protein *in vitro* (*66)*. Since SATB1 binds specifically to BURs *in vitro*, we used SATB1 as a biological probe to identify all potential BURs in the genome. Using this approach, we reproducibly identified approximately 240,000 BURs (q<0.01), establishing a BUR reference map for both mouse and human genomes. Comparing ChIP-seq peaks for SATB1 with the BUR reference map allows us to assess whether SATB1 binding sites *in vivo* correspond to BURs.

We performed urea ChIP-seq for SATB1 in thymocytes and compared the results with standard ChIP-seq for this protein and the BUR reference map. Strikingly, urea ChIP-seq produced SATB1 binding profiles that are entirely different from those produced by the standard ChIP-seq method (*Figure 2A*). The genome-wide analysis in mouse thymocytes using deepTools revealed that only 0.92% (an average of two replicates per ChIP-seq method) of peaks in urea SATB1 ChIP-seq coincided with those in standard SATB1 ChIP-seq (*supplementary Table 1 and Table 2*). In a large genomic region covering a 4.48Mb region in chromosome 17 (*Figure 2A, top*), a gene-rich region was found in the valley between BUR-enriched domains (*Figure 2A, track 3*), and standard ChIP-seq detects clusters of SATB1 binding sites in this gene-rich region (*Figure 2A, tracks 1 and 2*). In contrast, urea ChIP-seq detected SATB-binding to the domains surrounding the gene rich region, corresponding to regions heavily enriched in BUR sites (*Figure 2A, tracks 3-5*). In a zoomed-in view, covering a 430 kb region, SATB1 sites identified from urea ChIP-seq precisely coincided with BUR peaks and are excluded in the standard SATB1 ChIP-seq peaks (*Figure 2B, tracks 1-3*). SATB1 bound sites uncovered from urea ChIP-seq avoided CTCF sites (*Figure 2B, tracks 4 and 5 versus track 6)*, H3K27ac-marked sites and DNase 1 HS (*Figure 2B tracks 7 and 8)*, whereas SATB1 sites from standard ChIP-seq primarily co-mapped to these sites (*Figure 2B, tracks 1,2,6-9*). To unequivocally demonstrate that urea ChIP-seq peaks for SATB1 coincide with BURs, we examined these peaks shown in Figure 2B at a higher resolution (*Figure 2—figure supplement 1*). All four SATB1 peaks in Figure 2B precisely co-map with BURs. For one of these peaks, a typical BUR sequence is shown that contains a cluster of DNA sequence stretches in which either Gs or Cs are lacking from one strand. This DNA sequence context is referred to as the ATC sequence context. We confirmed the relationship of SATB1-binding sites by urea ChIP-seq and BURs at the genome-wide level using deepTools. Among SATB1 urea ChIP-seq peaks, an average of 77% of two replicates coincided with BURs identified in our reference maps (*Figure 2C*). The statistical significance of enrichment of BURs compared to a random overlap in either urea ChIP-seq or standard ChIP-seq peaks is p<0.001 or p=1, respectively, using 1000 bootstrapped genomic intervals sampled using the *bootRanges* approach (*71*) (*Figure 2C*). Consistent with this, genome-wide distribution profile analyses show that while SATB1 binding profiles by standard ChIP-seq specifically exclude BURs, showing a sharp indentation at the peak of BUR distribution, SATB1-binding profiles by urea ChIP-seq coincide with BUR distribution (*Figure 2D*). Heatmap analyses were also conducted to examine SATB1-binding sites among all BURs and BURs occurrence in all SATB1-bound sites in thymocytes *(Figure 2E).* Consistent with results shown in *Figures 2C* and *2D*, the heatmap study shows that the majority of SATB1-binding sites coincide with BURs. Also, SATB1 binds a small subpopulation of BURs in thymocytes (see below for cell-type dependent SATB1 occupancy of BURs). These results indicate that SATB1 predominantly and directly targets BURs genome-wide *in vivo* and that the SATB1-binding profiles obtained by standard and urea ChIP-seq methods are mutually exclusive. We conclude that our urea ChIP-seq, which solely captures DNA with its directly bound proteins, unmasks SATB1 direct binding profiles missed by standard ChIP-seq.

**Figure 2.**
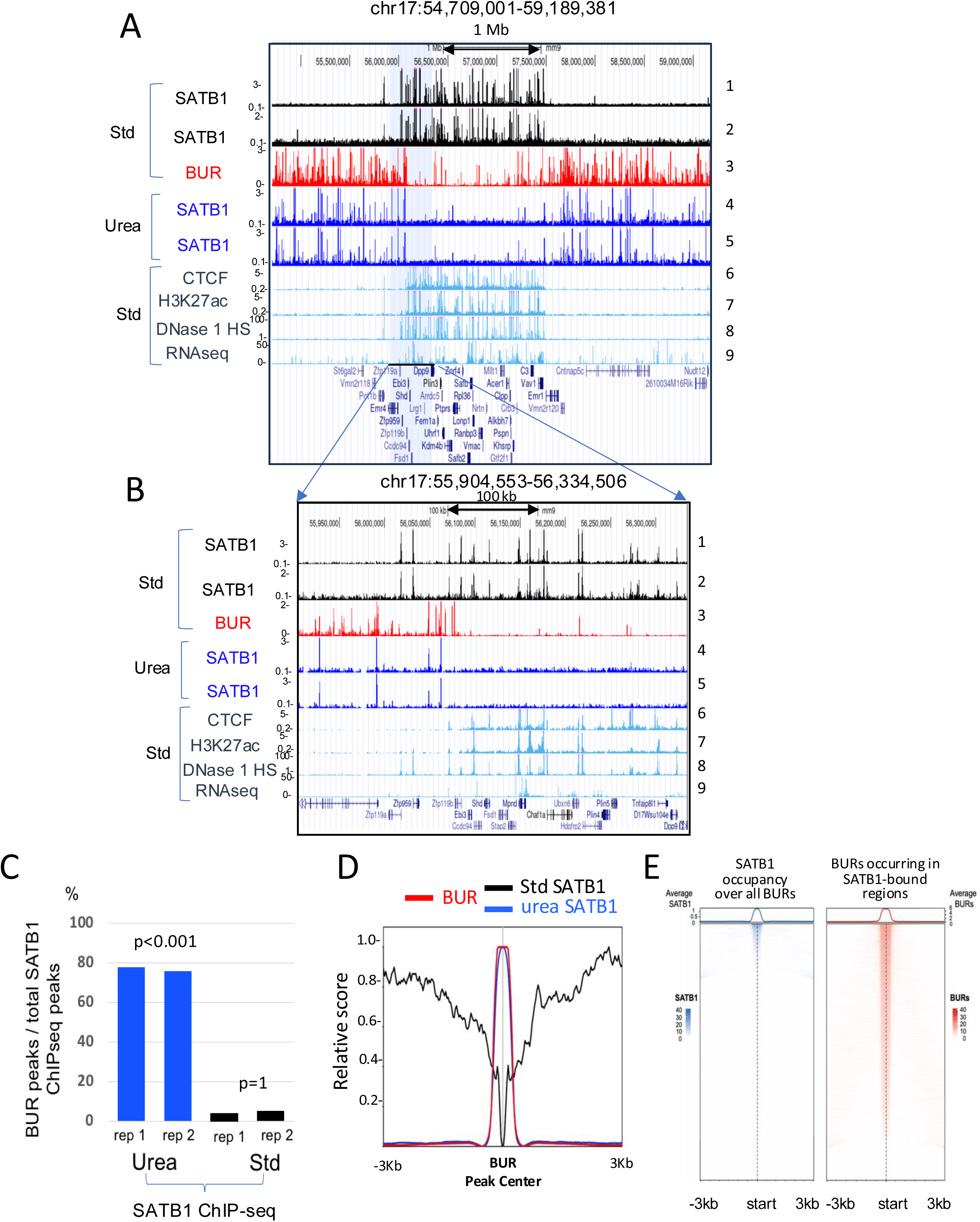
Identification of BURs as direct targets of SATB1 *in vivo* by urea ChIP-seq. A) Contrasting distributions of SATB1-binding profiles determined by urea ChIP-seq (Urea) compared to standard ChIP-seq (Std) in mouse thymocytes. B) A zoomed-in view of the gray-highlighted region shown in (panel A). The track 3 (panels A and B) represents all potential BUR sites serving as a reference for BUR sites. For both panels A and B, two independent urea and standard ChIP-seq experiments show high reproducibility (tracks 1 and 2 and tracks 4 and 5). Tracks 6 and 7 [ENCODE, Bring Ren’s Laboratory at the Ludwig Institute for Cancer Research (LICR)] and tracks 8 and 9 (ENCODE, the University of Washington (UW) group]. C) The percentages of urea (Urea) and standard (Std) SATB1 ChIP-seq peaks that intersect with the BUR reference map (see *supplementary Table 1*). The significance (p values) of the overlap in each of the four cases (urea and standard ChIP-seq samples with two replicates each) with BURs is estimated as the fraction of the overlap counts greater than or equal to the observed count over 1000 random bootstrapped genomic features. p<0.001: none of 1000 random fragment showed greater overlap with BURs than observed in ChIP-seq samples, p=1 means all 1000 random fragments showed greater overlap with BURs than observed in ChIP-seq, indicating avoidance of BURs (see Methods). D) Normalized genome-wide average intensity distribution profiles (normalized relative score) of SATB1 binding sites determined by standard ChIP-seq, urea ChIP-seq as well as BURs, over a ±3kb window centered at BUR peaks (top). Relative distance, projection, and Jaccard tests all indicate strong correlation (p< 0.05) between urea ChIP-seq peaks for SATB1 and BURs, but no overlap (p< 0.05) between standard ChIP-seq peaks for SATB1 and BUR peaks. E) (Left) Heatmaps showing SATB1 ChIP-seq signal intensity centered on all BUR elements. Signal is plotted within ±3 kb of each BUR center using 50 bp bins using EnrichedHeatmap package (v1.30.0). The line plots above the heatmaps display the average SATB1 signal across all regions. Only a subset of BURs exhibits strong SATB1 binding, consistent with selective occupancy. (Right) Heatmaps showing BUR signal intensity centered on SATB1-bound regions (from ChIP-seq peaks). The vast majority of SATB1 binding events are positioned within BURs, as evidenced by sharp, centered BUR signal at nearly all SATB1 peaks. Line plots reflect the average BUR signal profile across all SATB1 sites in each domain class. BUR intensities are derived from *in vitro* SATB1 BUR-binding assays.

### CTCF-binding profiles remain unchanged between urea and standard ChIP-seq

To investigate the potential cause for the discrepancy in the SATB1-binding profiles between the two ChIP-seq approaches, we applied the urea ChIP-seq method to other factors and asked whether binding profiles differed between urea and standard ChIP-seq. We investigated the DNA-binding profile for CTCF by urea ChIP-seq and found its profile to be very similar to that obtained by standard ChIP-seq from ENCODE, which largely overlapped with DNase 1 HS (*72*) (*Figure 3A, tracks 2, 4, 5 and 6*). Similarly, we also found that urea ChIP-seq generated essentially identical DNA-binding profiles for polycomb repressive complex 2 (PRC2) core subunits, SUZ12, JARID2 and EZH2 as generated by others using standard ChIP-seq (*73*) (*Figure 3—figure supplement 1*). In line with recent studies (*53-55*), 37% of SATB1 peaks from standard ChIP-seq (ENCODE) (average of two replicates, *supplementary Table 1*) intersected with CTCF peaks genome-wide. In contrast, only 0.8% of SATB1 peaks from urea ChIP-seq coincided with CTCF peaks (average of two replicas, *supplementary Table 1*), confirming the UCSC browser track views (*Figure 2A* and *3A*). These results verify that urea ChIP-seq can accurately identify the DNA-binding sites of chromatin associated proteins. Thus, urea ChIP-seq, compared to standard ChIP-seq, produces contrasting DNA-binding profiles for SATB1, but this is not necessarily the case for other nuclear proteins. Thus, we conclude that for SATB1, urea and standard ChIP-seq produces contrasting SATB1 binding profiles. Importantly, only urea ChIP-seq can faithfully detects direct and specific SATB1 interactions with BURs, suggesting a different mode of chromatin interactions. As mentioned in the introduction, it is important to note that these urea ChIP-seq results support a large body of prior binding assay and cytological data for SATB1, indicating that SATB1 forms salt-extraction resistant subnuclear protein network, and that the genomic sites directly bound to SATB1 reside in the “inaccessible” chromatin.

**Figure 3.**
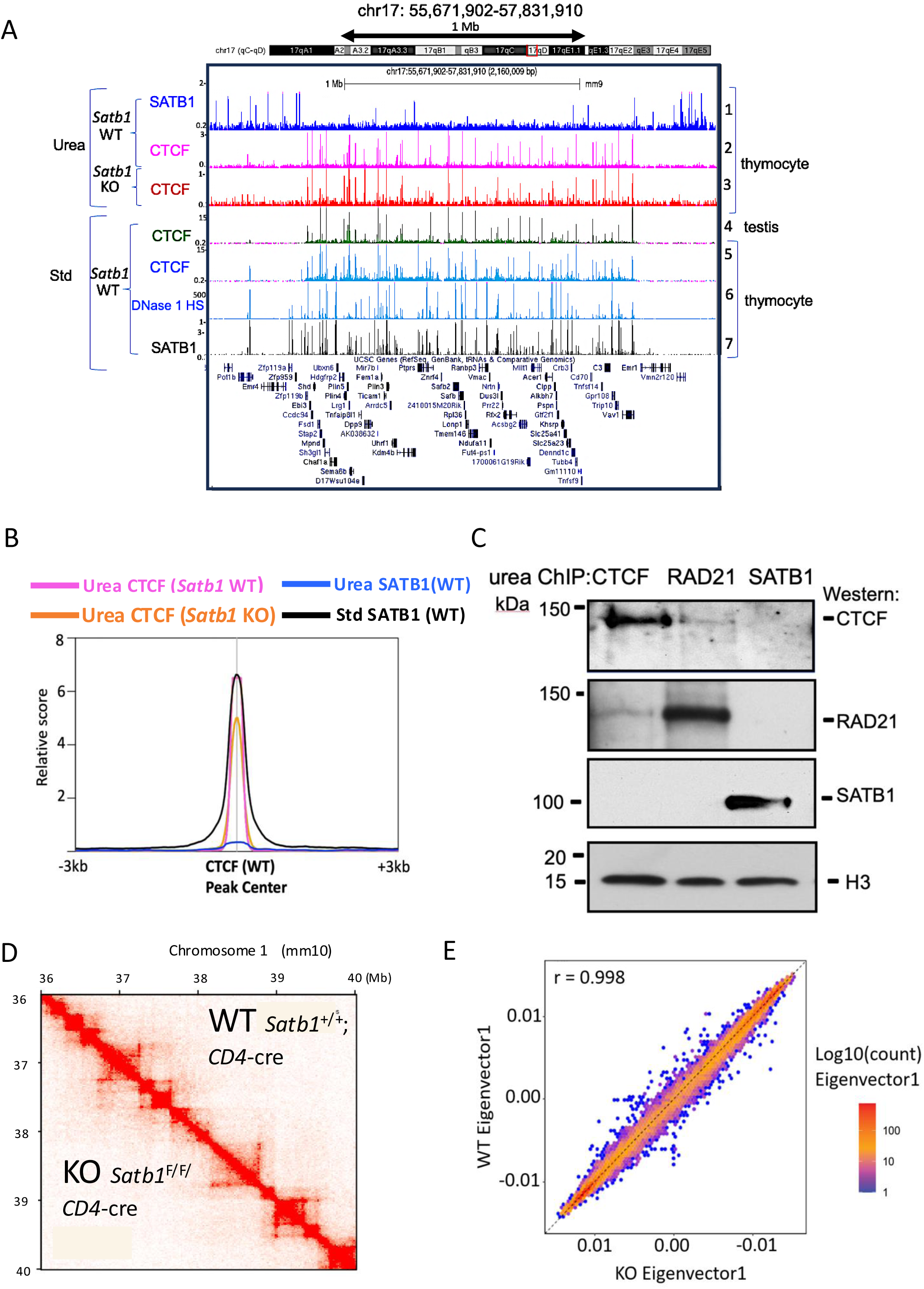
Identical CTCF-binding profiles are produced by urea and standard ChIP-seq, independent of SATB1. A) Urea ChIP-seq (Urea) profiles for CTCF in *Satb1*^+/+^ (WT) thymocytes and *Satb1*^−/−^ (KO) thymocytes (track 2 and 3), and for SATB1 in WT thymocytes (track 1). These binding patterns were compared against CTCF-binding profiles generated by standard ChIP-seq (Std) from ENCODE (tracks 4 and 5) for testis and thymocytes (ENCODE, LICR), respectively, along with the DNase1 HS profile from ENCODE, UW (track 6) as well as the SATB1-binding profile from standard ChIP-seq (track 7). B) Genome-wide average intensity distribution (relative score) of peaks from CTCF and SATB1 urea ChIP-seq for *Satb1*^+/+^ thymocytes (WT), CTCF urea ChIP-seq for *Satb1*^−/−^ thymocytes (KO), and SATB1 standard ChIP-seq (WT) over a +/− 3kb window centered at CTCF peaks (WT). Relative distance, projection, and Jaccard tests all indicate strong correlation (p< 0.05) between CTCF urea ChIP-seq peaks for WT and KO thymocytes, between these peaks and standard ChIP-seq peaks for SATB1, but no overlap (p< 0.05) between SATB1 urea ChIP-seq peaks. C) Urea-ChIP-Western results for CTCF, RAD21, and SATB1, showing that SATB1 does not associate with CTCF nor RAD21 on chromatin. Also, CTCF and RAD21 co-binding to chromatin appears to be infrequent, suggested by the weak ChIP-Western signals. D) Hi-C interaction heatmaps for *Satb1*^+/+^:*Cd4*-cre (WT) (top right triangle) and *Satb1* ^F/F^:*Cd4*-cre (KO) thymocytes (bottom left triangle) show essentially identical patterns. E) Genome-wide comparison of distributions of cis-Eigenvector 1 values shows no difference between WT and KO thymocytes (high correlation at r=0.998), indicating that genomic compartmentalization is unaffected in *Satb1* KO thymocytes. Experiments were done in two replicates each for WT and KO samples. The data shown in D and E were generated by combining data from the replicates for each condition.

### CTCF and SATB1 have distinct and independent binding profiles

Our results above indicate that CTCF and SATB1 *direct* binding sites do not overlap. However, because both CTCF and SATB1 are known to affect chromatin organization, there might be a functional link between the two proteins. Recently, CTCF-binding sites mapped by CUT&Tag were found to be SATB1 independent in specific gene loci (*55*). We validated this result by generating the CTCF-binding profiles by urea ChIP-seq in *Satb1*^−/−^ (SATB1-KO) and *Satb1*^+/+^ (SATB1-WT) thymocytes. These CTCF-binding profiles showed no major changes depending on SATB1 (*Figure 3A, compare tracks 2 and 3*), as shown in a randomly chosen region in chromosome 17 in wild-type (WT) and SATB1-KO thymocytes. This was confirmed by genome-wide analyses which showed that CTCF urea ChIP-seq peak distributions from SATB1-WT and SATB1-KO thymocytes coincided (*Figure 3B*). This indicates that CTCF binding to genomic DNA is SATB1-independent genome wide.

To corroborate the independence of CTCF/cohesin and SATB1, we next examined whether SATB1 interacts with CTCF and/or cohesin or both, components known to be important for loop formation. Distinct from protein-protein co-immunoprecipitation (co-IP) using whole cell or nuclear extracts, we examined the direct co-binding status on chromatin *in vivo* of SATB1 and CTCF or cohesin by urea ChIP-Western. Urea-purified crosslinked-chromatin fragments immunoprecipitated with anti-SATB1 antibody (SATB1-bound urea-ChIP fragments) contained SATB1 as expected, detected by Western-blot of the de-crosslinked urea-ChIP sample. However, neither CTCF nor RAD21 (a cohesin complex component) was detected in the SATB1 urea-ChIP samples. Reciprocal experiments using either CTCF or RAD21-bound urea ChIP samples confirmed lack of SATB1 co-binding to these chromatin fragments (*Figure 3C*, *Figure 3—figure supplement 2A*). We confirmed that a similar amount of urea-purified crosslinking chromatin was used for each ChIP-Western sample with an anti-H3 antibody (which is, as expected, retained on urea-purified crosslinked chromatin). Therefore, these results from urea ChIP-Western indicate that at least for the direct DNA co-binding status of SATB1 with either CTCF or RAD21, SATB1 does not co-bind chromatin with either of these proteins (*Figure 3C*). These results are consistent with mutually exclusive direct binding sites of CTCF and SATB1 (*Figure 3A* and *3B*).

If cohesin and CTCF bind directly to chromatin for loop formation according to the loop extrusion model, then we expect that the urea ChIP-Western results will show that cohesin-bound DNA fragments are co-bound by CTCF at the base of loops. As expected, we do find co-binding, however, fragments co-bound by RAD21 and CTCF represent only a small fraction of either total RAD21-bound or CTCF-bound chromatin fragments (*Figure 3C*). These results show that CTCF and RAD21 direct co-binding to chromatin occurs much less frequently than their individual binding to chromatin. This result agrees with a recent report which used super-resolution imaging of individual cells to show that the stable CTCF-mediated loops are rare, and >95% of the time, the loops are either only partially extruded, or no loops are formed (*25*).

Given that CTCF deletion causes most TADs to disappear, we next examined whether SATB1 depletion has any effects on TAD formation. We compared bulk Hi-C results between thymocytes isolated from wild-type (*Satb1*^+/+^: *Cd4*-*cre*) and SATB1-deficient (*Satb1*^F/F^:*Cd4*-*cre*) mice in duplicates (*Figure 3—figure supplement 2B*). Our Hi-C results revealed largely indistinguishable TAD profiles between the two thymocytes (*Figure 3D*) examined at 10kb resolution. Neither TAD number nor TAD sizes were changed by SATB1 depletion from thymocytes (*Figure 3—figure supplement 2C).* Intra-TAD interactivity (the sum of Hi-C chromatin contacts inside the TAD) also remained mostly unchanged (*Figure 3—figure supplement 2D*). These bulk Hi-C results upon SATB1 deletion are in good agreement with recently published studies (*53, 54*). Similar to the results from the CTCF depletion study (*18*), SATB1 depletion also did not alter segregation of active (A) and inactive chromatin compartmentalization (B) (*Figure 3E*). Furthermore, our scaling plot displaying how the Hi-C contacts decrease as a function of genomic distance shows that SATB1 depletion does not affect chromosome compaction (*Figure 3—figure supplement 2F*). These Hi-C results taken together with SATB1-independent CTCF-binding to DNA and non-overlapping direct genomic DNA-binding of SATB1 and CTCF/cohesin, strongly suggest that SATB1 and CTCF are two independent proteins that have distinct roles in chromatin organization, consistent with SATB1 having no roles in TAD organization.

### Non-specific peaks appear in *Satb1*^−/−^ thymocytes by standard SATB1 ChIP-seq

An important question is whether SATB1-binding profiles obtained by standard ChIP-seq are strictly SATB1 dependent. If so, these binding sites likely reflect real but indirect SATB1 association with chromatin. To address this question, we performed standard ChIP-seq on *Satb1*^−/−^ (KO) thymocytes using two different anti-SATB1 antibodies (1583 and ab109112, both of which are validated for immunoprecipitation). The results are highlighted in two randomly chosen ∼3Mb regions in chromosome 17 and in X chromosome, each containing a gene-rich region surrounded by gene-poor regions. Both antibodies generated similar peaks in KO thymocytes (KO peaks) by standard ChIP-seq. These KO peaks (false-positive) were predominantly detected in gene-rich, open chromatin regions in KO thymocytes, and many of them coincided with peaks originally detected in WT thymocytes (WT peaks) (*Figure 4—figure supplement 1 A and 1B, top and bottom, tracks 9 and 10*). This is in agreement with an earlier report that ChIP-seq generate false-positive peaks in highly accessible, transcriptionally active chromatin region (*56-60*). Intersection analyses with deepTools show that 4.1% of SATB1 standard ChIP-seq peaks obtained from our experiments are false-positive peaks (based on average of two standard KO ChIP-seq samples) (*supplementary* Table 3 and 4). Besides those KO peaks that overlapped with WT peaks, there were several thousands of KO-specific peaks generated by standard ChIP-seq in our study. In contrast, SATB1 urea ChIP-seq had no false-positive peaks (0.0%). Also, only ∼100 KO-specific peaks (not overlapping with WT peaks) were detected (average of two samples) compared to ∼40,000 total WT peaks from SATB1 urea ChIP-seq (*supplementary Table 3 and 4*). Therefore, validation using knockout cells is critical to distinguish real peaks that depend on SATB1 from SATB1-independent phantom peaks derived from the standard ChIP-seq method. The percentages of phantom peaks in standard ChIP-seq likely depend on the specificity of antibodies used and the condition of sample preparation.

**Figure 4.**
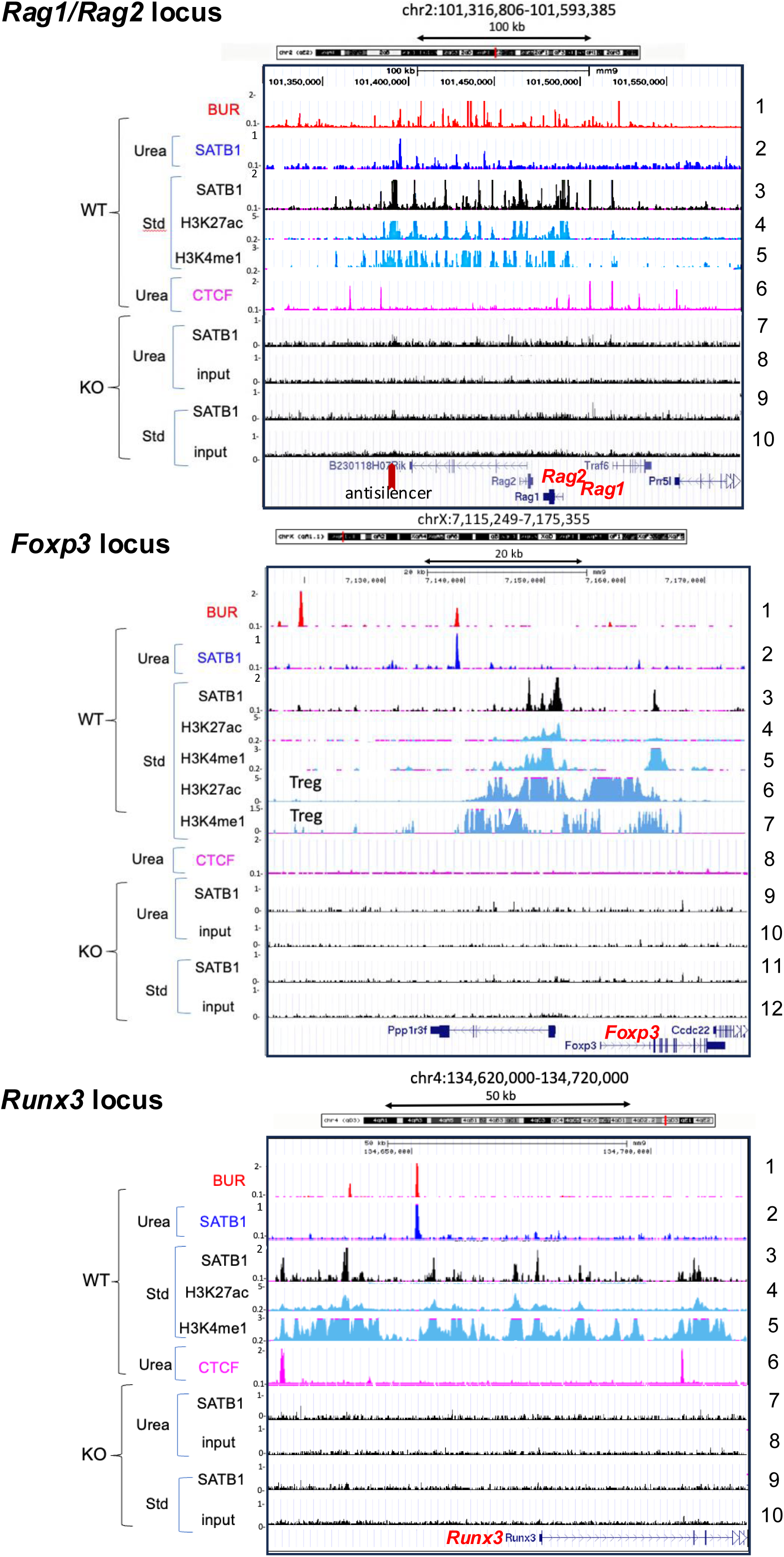
SATB1-bound sites identified by standard ChIP-seq at regulatory regions are SATB1 dependent. SATB1-binding profiles in immature thymocytes determined by urea ChIP-seq (Urea) and standard ChIP-seq (Std) covering three loci at *Rag1/Rag2* (top) *Foxp3* (middle), and *Runx3* (bottom) are shown. For each of the three loci, the standard SATB1 ChIP-seq generates peaks in *Satb1*^+/+^ (WT) thymocytes (track 3), but not in *Satb1*^−/−^ (KO) thymocytes (tracks 9 for *Rag 1/2* and *Runx3* and tracks 11 for *Foxp3*). Input chromatin for standard ChIP-seq in these regions did not produce any major peaks (track 10 in *Rag1/Rag2* and *Runx3* and track 12 in *Foxp3*). The SATB1 peaks from standard ChIP-seq in *Rag1/Rag2* and *Runx3* loci largely overlap with enhancers (marked by H3K4me1 and H3K27ac). At the *Foxp3* locus in immature thymocytes SATB1 peaks (track 3) appear in a region marked by low levels of H3K27ac and H3Kme1 (tracks 4 and 5) where super-enhancer is established upon differentiation into Treg cells (tracks 6 and 7)(*47*). SATB1-bound peaks produced by urea ChIP-seq (track 2) are consistently located in distinct positions from those peaks obtained by standard ChIP-seq in these gene loci. Urea ChIP-seq did not generate peaks in KO thymocytes with anti-SATB1 antibody. Input chromatin also did not produce any peaks. H3K27ac and H3Kme1 data (tracks 4 and 5) are from Encode, generated by Bing Ren’s laboratory at LICR, and data for Treg shown in (tracks 6 and 7) are from Kitagawa(*47*).

After validation with SATB1-KO cells, we conclude that most SATB1 sites detected from standard ChIP-seq are bound by SATB1, but most likely through indirect binding, as these peaks are not detected by urea ChIP-seq, which removes any indirect binding proteins by urea ultracentrifugation. SATB1 proteins are found in high salt-resistant fraction as well as salt-extracted fraction (*40*). Thus, it is possible that soluble SATB1 may associate with “open” chromatin. In order to further understand these differences and potential proximity of SATB1 binding sites detected by the two ChIP-seq methods, we focused on three individual gene loci, i. e. *Rag1* and *Rag2* genes (*45*), *Foxp3* (*47*), and *Runx3* (*46*), for which specific SATB1-binding sites by standard ChIP-seq have been reported (*Figure 4*). In addition, these genes are known to be regulated by SATB1 in thymocytes and are thus excellent candidate regions to investigate how SATB1 is regulating chromatin and gene expression. By either direct urea-ChIP or standard ChIP, we found SATB1-bound sites within all three loci (*Rag1/2*, *Foxp3*, and *Runx3*) to be SATB1-dependent. These sites were detected only in WT (*Figure 4, track 3*) but not in SATB1 KO-thymocytes (*Figure 4, tracks 7 and 9 for Rag/Rag2* and *Runx3*, *and tracks 9 and 11 for Foxp3*). In these regions, input sample (without antibodies) had no peaks (*Figure 4*, *tracks 8 and 10 for Rag1/Rag2 and Runx 3, and tracks 10 and 12 for Foxp3*). For the *Rag1* and *Rag2* loci, SATB1 has been shown by standard ChIP-seq to bind the anti-silencer element (ASE) and this binding is thought to counteract the activity of an intergenic silencer, thus promoting expression of *Rag 1* and *Rag 2* (*45*). During development of Foxp3^+^-regulatory T cells (Treg) in the thymus, SATB1 expression in Treg precursor cells is essential for the future establishment of a Treg cell–specific super-enhancers at the *Foxp3* locus in Treg cells (*47*). Thus, thymocyte precursor-specific SATB1 deficiency, using *Satb1^fl/fl^* CD4-cre mice but not *Satb1^fl/fl^ Foxp3*-cre mice, impairs Treg super-enhancer activation and expression of *Foxp3*(*47*). Impaired Treg cell development results in autoimmune disease (*46, 47, 74, 75*). Finally, SATB1 is also essential for generation of CD4^+^ and CD8^+^ T cells and NKT cells (*46*). In response to T cell receptor signaling by MHC class I and II, SATB1 regulates enhancer function in gene loci encoding lineage-specifying factors (e.g. *Runx3*) and directs lineage-specific transcriptional programs in the thymus (*46*). Strikingly, in these three regions, SATB1-bound BURs, as detected by urea ChIP-seq, are found infrequently, and appear as isolated single peaks within 30-50kb regions. Such SATB1-bound BUR sites are located close to but not overlapping with the cluster of enhancers marked by H3K27ac and H3Kme1 modifications. These SATB1-bound BUR sites reside within 3kb of the nearest putative SATB1-bound enhancers (detected by standard ChIP-seq) for the *Rag1/Rag2* and *Runx3* loci as well as from the super-enhancer that becomes established at the *Foxp3* loci in matured Treg cells. Results from these three loci show that, while the SATB1-bound peaks generated by urea ChIP and standard ChIP-seq did not overlap, they were nonetheless proximal to each other, and both were SATB1 dependent. The two distinct ChIP-seq protocols are likely detecting two different modes of DNA binding for SATB1 i.e., direct binding to BURs and indirect binding to open chromatin regions.

### Long-distance chromatin interactions involving a BUR depend on SATB1

We next explored whether there is any three-dimensional relationship between SATB1-bound BURs detected by urea-ChIP-seq and SATB1-bound sites in the open chromatin region detected by standard ChIP-seq. We hypothesized that chromatin architecture formed by direct binding of SATB1 to BURs may recruit open chromatin regions by indirect association with their chromatin-bound protein complexes by SATB1. If so, such interactions would necessarily be dependent on SATB1, and they might be correlated to gene expression. To address these questions, we studied a gene-rich region in chromosome 2 that contains the *Rag1* and *Rag2* genes whose expression is SATB1 dependent (*45*). We first examined genome-wide interactions from a SATB1-bound BUR that appeared as a single, strong peak (*Figure 4, top track2, and 5, top)* as the viewpoint or bait (BUR-1). The BUR-1 primer sequence is located near the border between the 3.9 Mb gene-desert region and the 5.8Mb gene rich region on chromosome 2 where the major BUR bound to SATB1 is located. BUR-1 resides in the latter accessible chromatin region and 1.4 kb away from the ASE (*Figure 4, top, track2*), which is essential for proper *Rag1* and *Rag2* expression in double-positive thymocytes and for differentiation of single-positive thymocytes (*76*).

To study interactions from the SATB1-bound BUR, we performed 4C-seq using urea purified chromatin (urea 4C-seq, one-to-all) to better access and capture BUR-mediated interactions, which are expected to emanate from the SATB1 subnuclear architectural structure. In our urea 4C-seq, we used *Hind*III for initial digestion. The 6-base cutter, *Hind*III, was used instead of a 4-base cutter like *Dpn*II (frequently used for 4C-seq), because multiple *Dpn*II sites are often found within individual SATB1-bound BURs, thus producing small fragments, likely disrupting SATB1-binding to BURs. The “BUR-1” bait region at a SATB1-bound BUR is designed at the end of the *Hind*III fragment, which is adjacent to the *Hind*III fragment containing ASE. We examined interaction events from the fragment containing BUR-1 and its interaction sites were mapped on the genome for visualization.

While urea purification removes indirectly associated protein and thus may reduce some chromatin interactions, we speculated that the overall 3D chromatin structure of fixed chromatin will still retain its proximity events, especially in the environment of chromatin which is strongly anchored to the subnuclear structure (as we predicted for SATB1 scaffold anchoring at BURs) to enable detection in urea 4C-seq. In the ChIP-seq protocol (in general), we expect that in the absence of a ligation step as is routinely done in 4C protocols (as described below), subsequent steps for ChIP-seq (sonication, stringent washes of immunoprecipitated fragments, etc) results in dissociation of interacting fragments. We carried out urea 4C-seq experiments to test if we could indeed detect such strong SATB1-mediated 3D interactions. Briefly, for urea 4C-seq (outlined in *Figure 5—figure supplement 1*), crosslinked chromatin purified by urea ultracentrifugation is first digested with a restriction enzyme (*Hind*III). Digestion is followed by ligation of chromatin fragments whose physically proximity status in 3D space was fixed at the time of formaldehyde crosslinking. After these steps, DNA was purified. Beyond this point urea-4C-seq differs from the commonly used 4C-seq called circular chromosome conformation capture (referred to here as circular 4C-seq)(*77-82*). In circular 4C-seq, ligated DNA fragments are circularized after second restriction enzyme digestion before PCR amplification. In contrast, in urea 4C-seq a single strand DNA capture strategy was used to isolate a specific pool of single strand DNA, each containing a ligation-captured DNA fragment surrounded by the bait fragment and the DNA linker (see details in *Figure 5—figure supplement 1* legend and Materials and Methods). The captured DNA sequences in the single strand DNA were PCR amplified using a primer from the bait sequence and another primer from the DNA linker, thus avoiding amplification of any non-specific DNA fragments. This was confirmed by the lack of PCR products using either primer alone (data not shown). This critical step results in highly specific signals with very low background signals. The high signal-to-noise ratio in urea 4C-seq contrasts with uniform and strong genome-wide background signals typically seen with circular 4C-seq, which are thought to reflect random chromatin interaction events. Some non-interacting sequences could also be trapped at the circulization step, and even if such incidence is rare, would result in background signals upon PCR amplification. Because the background signals for urea 4C-seq are extremely low in contrast to circular 4C-seq, the visual appearances of the interacting profiles from the two methods are greatly different. Nevertheless, urea 4C-seq detects large-scale interactions involving BURs that are strictly dependent on SATB1 as described below.

**Figure 5.**
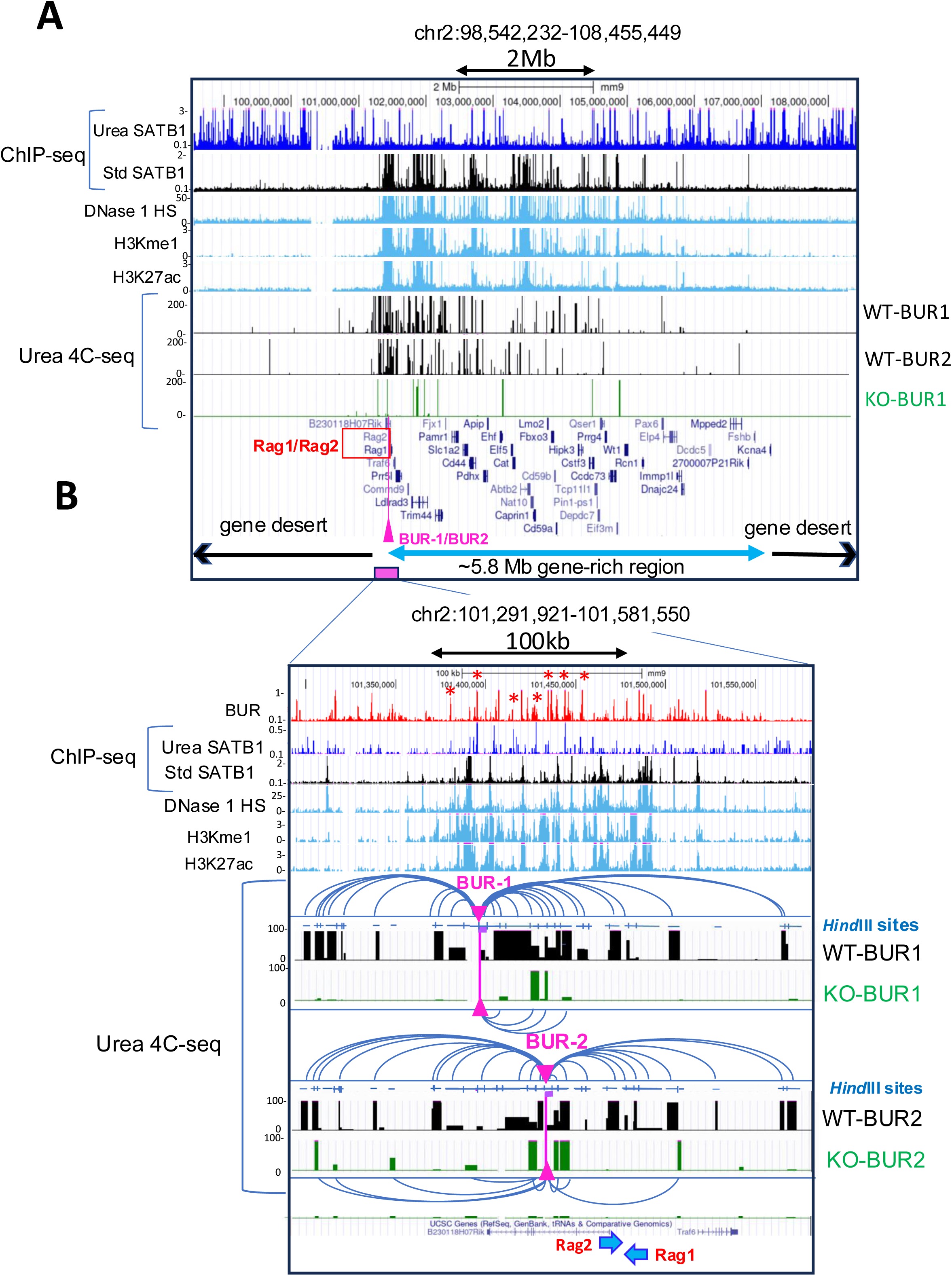
SATB1 mediates dense chromatin interactions over *Rag1/2*-containing gene-rich region. A) Chromatin interactions detected by urea 4C-seq using a SATB1-bound BUR (BUR-1) and another site (BUR-2) as baits. These baits are mapped in an accessible chromatin and located at near the border of a gene-poor region and a gene-rich region containing SATB1-regulated *Rag1* and *Rag2* genes (pink vertical bar with arrowhead). BUR-2 was found to be one of the interacting sites of BUR-1. Therefore, it was used as a reciprocal control for BUR-1 to check the reproducibility of BUR-1 interactions. We used *Satb1^+/+^* (WT) and *Satb1^−/−^* (KO) thymocytes from 2 weeks old mice for urea 4C-seq. BUR-1 and BUR-2 interacts with many sites within the ∼5.8Mb gene-rich region, containing *Rag1* and *Rag2* (tracks WT-BUR1 and WT-BUR2). These interactions were greatly reduced upon SATB1 deletion (track KO-BUR1). B) A zoom-in view, focusing on the *Rag1* and *Rag2* region, shows that BUR-1 interacts extensively over this region in WT thymocytes (shown by parabolas). Such interactions were virtually diminished in the absence of SATB1 (KO-BUR1 and KO-BUR2 tracks). The pink vertical bars with arrowheads indicate the positions of the BUR-1 and BUR-2 bait primers. Each of these primer sequences is located at the end of the *Hind*III fragment containing the primer. These *Hind*III fragments are marked by thick purple bars adjacent to pink arrowheads pointing primer positions. The map of *Hind*III fragments is shown for interacting regions (track *Hind*III sites). BUR-1 and BUR-2, along with other BURs bound by SATB1 (as confirmed by SATB1 urea ChIP-seq peaks and the BUR reference) that interact with BUR-1 or BUR-2 are marked by small red stars. Note: Reads to the *Hind*III fragment immediately adjacent to each of the bait primers (BUR-1 and BURs) were removed as they comprised more than 2% of the total reads, typically derived from re-ligation of digested chromatin fragments to their original neighboring sequences and/or from a small undigested chromatin fraction.

In wild-type thymocytes, we detected BUR-1 making extensive long-distance interactions covering most of the ∼5.8Mb adjacent gene-rich region (*Figure 5A*). Interestingly, BUR-1 interactions are found to be scarce over the gene-desert region upstream of BUR-1, despite the presence of many SATB1-bound BURs in this region (*Figure 5A, top track for urea SATB1 ChIP-seq, Figure 5—figure supplement 2*). Though 4C-seq can detect both up- and downstream interactions from a chosen single bait site, we observed heavily direction-biased interactions of BUR-1 with the gene-rich region, perhaps indicative of its involvement in gene regulation. BUR-1 interactions were found particularly concentrated over the ∼3 megabase (Mb) gene-rich region, containing SATB1-dependent *Rag1* and *Rag2* genes. These dense and strong interaction signals must not be due to their spatial proximity from the bait because similar dense interactions were not detected with the SATB1-KO cells. In fact, most of the BUR-1 interactions were absent in SATB1-KO cells (*Figure 5A, track KO-BUR1*), indicating that SATB1 is required for the numerous long-distance interactions over the gene-rich region. We also validated these interactions using a second bait (BUR-2) as an experimental control as BUR-1 interacts with the *Hind*III fragment containing BUR-2, located in the second intron of *B230118H07Rik* gene (see *Figure 5B* for specific location). This fragment contains two strong BUR peaks. Although weakly, SATB1 binds to one of these BURs. Hence, urea 4C-seq using BUR-2 serves as a reciprocal experiment of BUR-1 to validate reproducibility of the BUR-1 interaction profile. By comparing with the Hi-C heatmap in this 5.8Mb region we found that BUR-1 interactions traverse multiple TADs and are not constrained by any specific TADs (*Figure 5—figure supplement 2*), further supporting the CTCF independent nature of these SATB1-mediated interactions. Within this gene rich region, numerous interactions from BUR-1 were detected within DNase 1 HS containing enhancers marked by H3K27ac and H3Kme1 modifications. Therefore, dense chromatin looping events involving these SATB1-bound BURs indicate a highly interactive chromatin architecture for the gene-rich region, bringing together many gene regulatory sequences into proximity. We further examined a wider region covering >44Mb that includes distal neighboring gene-rich regions *(Figure 5 and Figure 5—figure supplement 3A and 3B).* Although interaction frequencies were lower than that within the first ∼5.8Mb gene-rich region, we detected BUR-1 interactions with further distal gene-rich region coinciding with enhancer-rich DNase 1 HS regions, where clusters of SATB1 binding sites were detected by standard ChIP-seq, skipping over gene-poor regions and a gene-rich region containing transcriptionally silent olfactory receptor genes.

A zoomed-in view (chr2:101,308,846-101,599,827; 291Kb) of BUR-1 interactions confirmed a marked difference in BUR-1 interactions (∼200Kb) over the *Rag1* and *Rag 2* loci between wild-type and *Satb1*^−/−^thymocytes, confirming BUR-1 interactions require SATB1 (*Figure 5B*). In addition to many BUR-1 interacting chromatin fragments close to or containing enhancers (marked by H3K27ac and H3Kme1), some of these fragments contain additional BUR sites, including BUR-2 adjacent to enhancer(s) *(Figure 5B,* gray shaded with red stars). BUR-2 interacted heavily over the *Rag1* and *Rag2* loci in wild-type thymocytes and well reproduced the interactions observed with BUR-1, consistent with the larger region (*Figure 5A*), suggesting that both BUR-1 and BUR-2 are part of the common interaction network that requires SATB1. Concerning the correlation of chromatin interactions and gene expression, among 43 genes identified with official gene symbols mapped in the 5.8Mb gene rich region, 8 genes display various levels of SATB1-dependent expression (p<0.05) (*54*). These genes were either up or downregulated upon SATB1 deletion in thymocytes based on RNA-seq data (*Figure 5-figure supplement 4*), and *Rag1* and *Rag2* were most strongly downregulated by SATB1 depletion in this region. Collectively, these data suggest a strong correlation between SATB1-dependent gene expression and extensive looping events over long distance linking SATB1-bound BURs and regulatory regions in the open chromatin regions. Therefore, SATB1 function on chromatin is not limited to merely connecting a specific target gene locus with nearby enhancers and promoters within TADs. Instead, SATB1 appears to form a large interaction network covering the entire gene-rich regions.

### SATB1 binds BURs in a cell-type dependent manner

We next examined whether SATB1 binds different subsets of BURs depending on cell type, by studying SATB1-binding profiles in three different types of mouse tissues– thymus, frontal cortex and skin – that contain SATB1-expressing cells. SATB1 expression is primarily restricted to adult progenitor cells, such as thymocytes and in basal layer of epidermis. SATB1 is also expressed in differentiated neurons in specific brain subregions (e.g. frontal cortex and amygdala). We compared UCSC browser images of two selected regions, a ∼2.4 Mb region in chromosome 6 containing the *Cd4* locus expressed in thymocytes (*Figure 6A*) and a 406kb region in chromosome 5 containing *Gabra2* and *Gabrg1* loci expressed in the mammalian brain (*Figure 6B*). BURs were most frequently bound by SATB1 in brain compared to thymocytes, and the least frequently bound in skin. Whereas SATB1-bound BURs show cell-type dependency, CTCF binding sites that largely coincide with DNase1 HS were largely similar between these three types of cells (*Figure 6A*).

**Figure 6.**
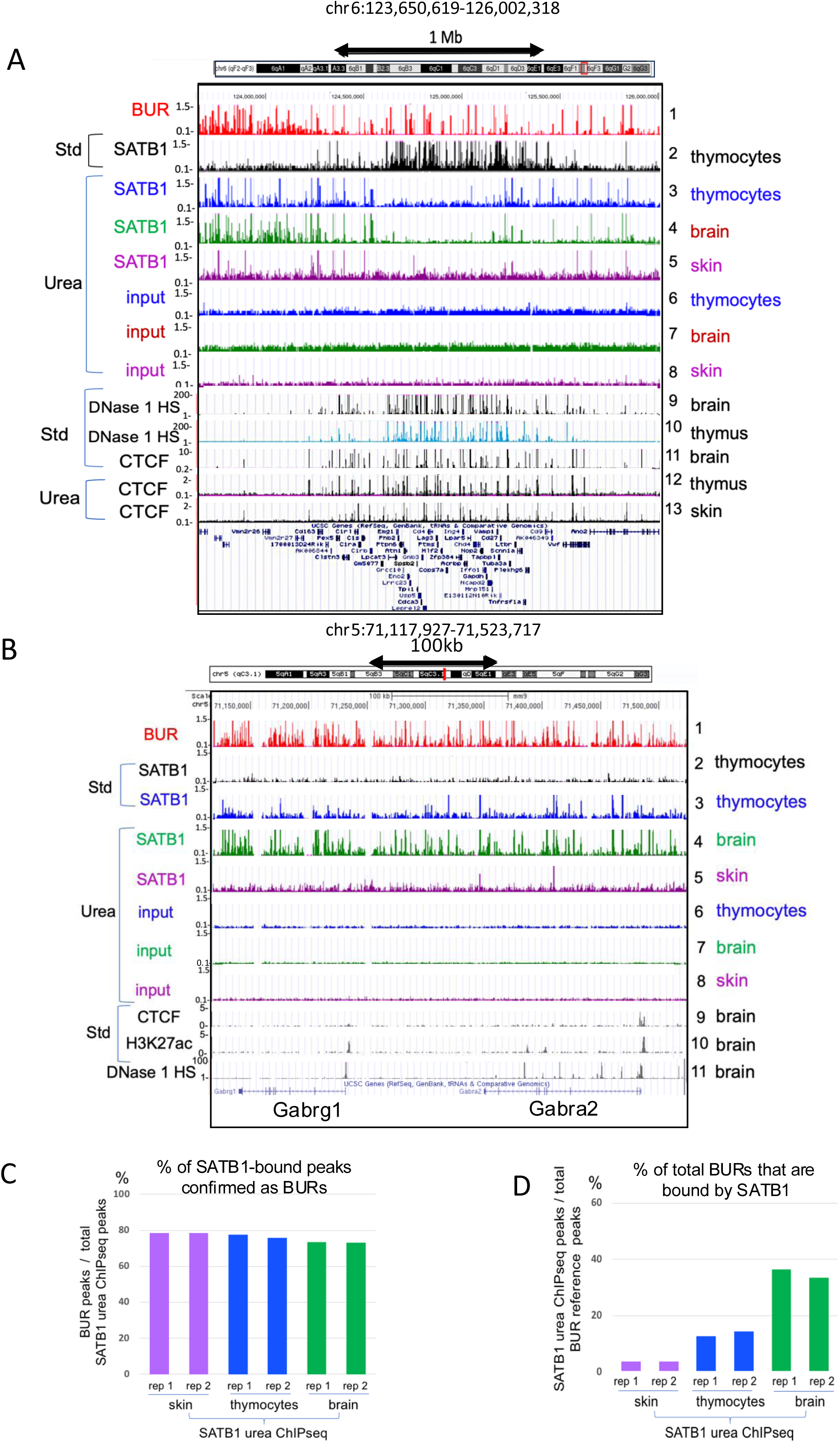
SATB1 binds to BURs in a cell-type dependent manner. A) SATB1-binding profiles in a ∼2.4 Mb region in chromosome 6 containing *Cd4* in a gene-rich region from standard ChIP-seq (Std) (track 2) and urea ChIP-seq (Urea) (tracks 3-8). Urea ChIP-seq-derived SATB1 profiles in thymocytes (track 3), brain (track 4) and skin (track 5) show difference in peak numbers depending on cell type (see *supplementary Table 1* and *2*). This contrasts with mostly identical CTCF patterns in these cell types (tracks 11-13). B) SATB1-binding profiles in a 406kb region in chromosome 5 containing *Gabra2* and *Gabrg1*, highly expressed in brain, also shows much increased BUR binding in brain, compared to thymocytes and skin similar to A). In this region, there is no SATB1-binding site detected by standard ChIP-seq (track 2), and CTCF binding sites are mostly absent (track 9). C) Percentages of SATB1 urea-ChIP-seq peaks in skin, thymocytes and brain intersecting with the BUR reference map, indicating that the majority of the peaks are BURs (*supplementary Table 1*). D) The percentage of BURs among all BURs identified in the BUR reference map that intersected with urea SATB1 ChIP-seq peaks identified in skin, thymocytes and brain. DNase1 HS data are from ENCODE (UW), the CTCF-binding profile and H3K27ac data from standard ChIP-seq are from ENCODE (LICR).

We analyzed whether BURs are targeted by SATB1 depending on cell type. Consistent with the cell-type specific function of SATB1, we observed cell-type dependent differences in BUR occupancy by SATB1. For all three types of cells, high percentages of urea ChIP-seq peaks (close to 80%, *supplementary Table 1*) correspond to BURs, as revealed by peak intersection studies using deepTools (*83*) and comparing to the BUR reference map (*Figure 6C*, *supplementary Table 1*). Despite similar total aligned sequence reads in urea ChIP-seq, the number of SATB1-bound BUR peaks greatly differed between cell types (*supplementary Table 2*). Among 238,380 mapped BURs (q<0.01), the percentage of SATB1-bound BURs in skin, thymocytes, and brain were 3.7%, 13.5% and 35.0% (based on average of two replicas/cell type), respectively (*Figure 6D and supplementary Table 1*). These results indicate that frequency of BURs targeted by SATB1 *in vivo* can vary more than 10-fold depending on cell type. Interestingly, while the number of BURs bound by SATB1 greatly differs between cell types, the majority (83.9%) of BURs targeted by SATB1 in skin were bound to SATB1 in thymocytes, and nearly all (91.6%) SATB1-bound BURs in thymocytes were bound to SATB1 in brain. Therefore, each cell type has an overlapping set of SATB1-bound BURs in common, with increasing/decreasing numbers of sites depending on cell type, rather than distinct subsets. Despite the highest BUR occupancy by SATB1 was observed in the brain, this is not due to the brain expressing the highest levels of SATB1 protein. In fact, the thymus expresses by far the highest levels of SATB1, which comprises ∼0.3% of the total protein (*84*).

### BURs are enriched in LADs

LADs are large mostly heterochromatic domains in close contact with the nuclear lamina and are known to be enriched in AT-rich sequences (*85*). Although AT-rich sequences are not necessarily BURs, BURs have a minimum of 65% AT content (many of them have a higher AT content) with the ATC sequence context with an exceptionally strong unwinding propensity (*41*). We therefore asked whether and to what extent any of the ∼240,000 BURs identified in the genome could be found within LADs. We first visually compared the distribution of all potential BURs with LADs mapped by DamID (*86-88*) and noted some correlation of BURs and LADs. As an example, the 9 Mb region in chromosome 5 containing the neuronal gene cluster locus that encodes subunits of GABA_A_ receptor complex (*Gabrg1,* and *Gabra1, Gaba6, Gabrb2)* has a BUR-enriched region overlapping with LADs obtained from thymocytes (*Figure 7A*). We checked if this LAD-BUR overlap is not restricted to this specific region by examining a larger view of a randomly chosen region covering 47.4 Mb in chromosome 3 (*Figure 7—figure supplement 1*). In this region CTCF-binding sites, DNase1 HS, and standard SATB1 ChIP-seq peaks all converge in the gene-rich inter-LAD regions, whereas distribution of LADs and BURs largely overlap, and these regions are enriched in SATB1 urea ChIP-seq peaks.

**Figure 7.**
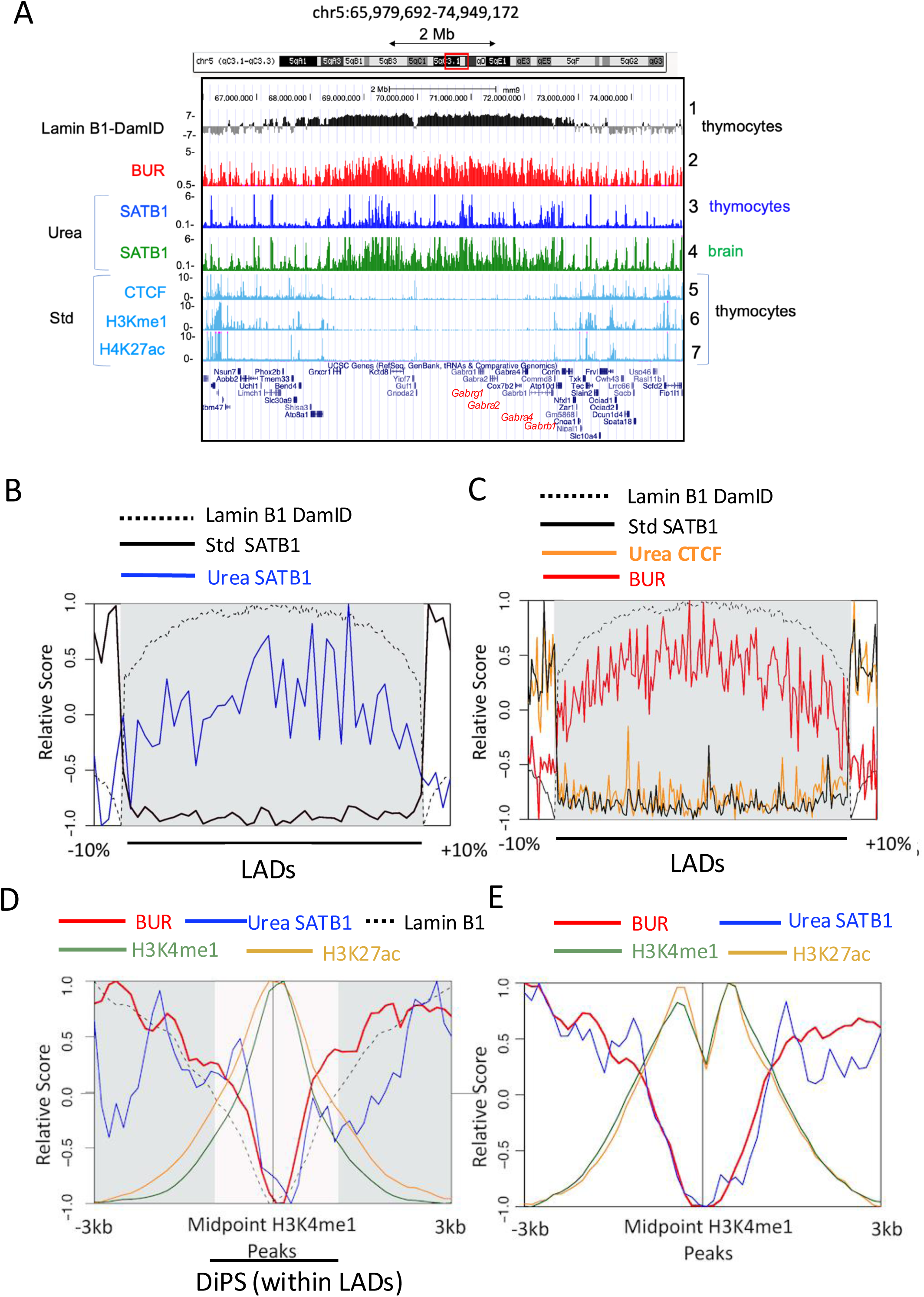
BURs largely co-map with LADs. A) The BUR distribution (BUR reference map, track 2) was compared against Lamin B1-DAM-ID sites(*86*) (track 1) and SATB1-binding sites derived from urea-ChIP-seq (Urea) (track 3 and 4) over a 8.97 Mb region in chromosome 5 containing neuronal genes, *Gabrg1*, *Gabra2, Gabra4,* and *Gabrb1*. In this region, co-mapping of BURs and Lamin B1-DAM-ID regions is confirmed. Among BURs, SATB1-bound BURs derived from urea ChIP-seq (Urea) show difference in the number of peaks depending on cell type (see *supplementary Table 1* and *2*). SATB1-binding sites derived from urea ChIP-seq avoid CTCF-binding sites, H3Kme1, H3K27ac sites (from Encode LICR). B) Average Lamin B1 DamID signal (black dotted), standard ChIP signal for SATB1 (black) and urea ChIP for SATB1 (blue) signal over LADs and flanking regions (interLAD sized at10% of adjacent LAD). C) Average Lamin B1 DamID signal (dotted black), BUR signal (red), and urea ChIP for CTCF (orange) signal over LADs (gray shaded) and flanking regions (interLAD sized at 10% of adjacent LAD). D) Average BUR signal (red), urea ChIP for SATB1 (blue) signal, Lamin B1 DamID (black, dotted), H3K4me1 (light green) and H3K27ac (light orange) signals over a +/− 3kb window centered at H3K4me1 peaks within LADs (LADs are gray shaded and DiPC that reside inside large LADs are unshaded, see text). E) Average BUR signal (red), urea ChIP for SATB1 (blue) signal, H3K4me1 (light green) and H3K27ac (light orange) signals over a ±3kbswindow centered at H3K4me1 peaks outside LADs. Relative distance, projection, and Jaccard tests all indicate strong correlation (p< 0.05) between urea ChIP-seq peaks for SATB1 and LADs, between BURs and LADs, but no overlap (p<0.05) between urea ChIP-seq SATB1 and H3K27ac and H3K4me3 peaks. The results in B to E represent normalized average intensities of signals.

We then examined the relationship between BUR and LAD distribution at the genome-wide scale. We compared the genomic proximity to lamin B1 (as measured by Dam), SATB1-bound genomic sites (from standard ChIP-seq), SATB1-bound BURs (from urea ChIP-seq) over LADs and flanking regions (+/− 10% outside of the LADs). We found that the genome-wide distribution of LADs coincides with the majority of SATB1-bound BURs, but it is mutually exclusive with SATB1-bound genomic sites that are mapped by standard ChIP-seq, which avoid LADs (*Figure 7B*). In agreement with the result shown in *Figure2A*, CTCF-bound sites mapped by urea-ChIP-seq coincide with those mapped by standard ChIP-seq and are found at LAD borders and inter-LAD regions (*Figure 7C*), as previously described (*9*). Collectively, these analyses confirmed that the majority of potential BURs from the BUR reference map as well as directly SATB1-bound BURs overlap with LADs (Figure 7B and C), whereas CTCF-binding sites avoid LAD-BUR regions and are confined to inter-LADs and LAD boundaries (*Figure 7C*). We conclude that, at the genome-wide scale, the majority of BURs overlap with LADs, suggesting that BURs represent as an important sequence component of LADs, some of which are specifically targeted by SATB1. This agrees with earlier findings that LADs are enriched in AT sequences as many BURs are AT-rich (>65% AT) and further identifies LADs as specifically enriched in BURs.

Within larger LAD domains identified, there are smaller regions (<10kb) that display localized looping away from the lamina (*89-91*). These DiPS (Depleted in Peripheral Signal) display all the hallmarks of poised or active promoters or enhancers and are flanked by CTCF and cohesin. Thus, these small regions embedded in larger LADs, are akin to small inter-LADs that are constrained at the lamina due to their short lengths. We next asked if SATB1 binds BURs in these active regions within LAD (DiPS). We examined the distribution of average signals of SATB1-bound BURs relative to enhancers in DiPS marked by H3K4me1 and H3K27ac modifications over a +/− 3kb window centered at H3K4me1 peaks. The peaks of enhancer histone marks (H3K27ac and H3K4me1) corresponded to minimum occupancies of SATB1 in DiPS and also excluded BURs, indicating that enhancers are largely excluded from BURs and SATB1-directly bound BURs in DiPS (*Figure 7D*).

We also examined SATB1-bound BURs outside LADs (inter-LADs) as well. Although less-densely distributed compared to LADs, SATB1-bound BURs are detected in inter-LAD regions enriched in actively transcribed genes. As an example, we show SATB1-bound BUR distribution in representative inter-LADs containing *Rag1/Rag2* (this region was used for 4C analyses above) as well as other two other SATB1-regulated loci*: Foxp3,* and *Runx3* relative to nearby LADs (*Figure 7—figure supplement 2*). Genome-wide analyses showed that in inter-LAD regions, SATB1-bound BURs are generally excluding the peaks of enhancers marked by H3K4me1 and H3K27ac (Figure 7E). Although the positions are not precisely coincident, ∼10% of SATB1-bound BURs in inter-LADs are located close (within +/− 5kb) to the peaks of H3K4me1 or H3K27ac. We viewed relative occupancies of BURs and SATB1-bound sites in LADs and inter-LADs by heatmap *(Figure 7—figure supplement 3)*. Consistent with the results described above, SATB1 occupancy is restricted to BUR sites, a fraction of BURs is targeted by SATB1 in both inter-LADs and LADs (the former being a much fewer number of BURs). Also, for both LADs and inter-LADs, most of the SATB1-bound regions coincide with BURs. We also examined the distribution of urea4C-detected BUR-1 or BUR-2 interactions relative to LADs. These SATB1-mediated interactions were found to be well constrained within inter-LADs enriched in histone marks for enhancers *(Figure 7—figure supplement 4).* This remains the case even for sites with much longer interactions, as these sites are located in accessible regions between LADs, confirmed by viewing individual sites at high resolution. These results demonstrate that the inter-LAD BURs (BUR-1 and BUR-2), bound to SATB1 are contacting accessible chromatin at numerous locations over long distances.

## Discussion

Here we used a systematic approach to compare and interrogate the chromatin binding profiles of a genome organizing protein SATB1 detected by two alternative ChIP-seq approaches. Critically, by establishing a urea purification-based ChIP-seq method that enriches for direct DNA-binding events, we found that SATB1 directly binds BURs genome-wide. This is in stark contrast to previously identified SATB1 binding to enhancers and promoters in open chromatin by standard ChIP-seq. Nevertheless, the two contrasting DNA-binding profiles for SATB1 are both valid, requiring SATB1 for the ChIP-seq peaks. These differing SATB1-binding profiles suggest that SATB1 is engaged in chromatin organization through both direct binding to BURs, anchoring them to the subnuclear SATB1 protein meshwork, as well as by indirect binding to the accessible (active) chromatin region, enriched in genes and regulatory regions. We show that SATB1 is required for megabase-level interactions, connecting select BURs to “open” chromatin, and that such interactions are linked with cell-type specific gene expression regulated by SATB1 in thymocytes.

Using urea ChIP-seq, which stringently purifies chromatin, we identified a new type of chromatin organization formed by direct binding of SATB1 with BURs, genomic elements distributed across the mouse genome. While this SATB1-mediated chromatin organization contrasts with previous data from standard ChIP-seq or CUT&Tag assays, which indicate SATB1 mainly interacts with enhancers and promoters located within the “open chromatin” regions, our data agree with even earlier *in vitro* findings that SATB1 binds to AT-rich BURs. Thus, standard ChIP-seq (or Cut&Tag) and urea ChIP-seq enrich different and almost mutually exclusive SATB1 profiles, suggesting that these approaches are interrogating the genome in different ways. It is important to note that while urea ChIP-seq and standard ChIP-seq for SATB1 showed very different profiles, the binding profiles of CTCF and other proteins did not differ between the two methods. This is consistent with SATB1 interacting with chromatin in a unique way by forming a subnuclear proteinaceous network anchoring a subset of BURs (*39*).

### Differences between standard and urea ChIP-seq

In order to understand why chromatin binding proteins might show different binding modalities in urea ChIP-seq versus more standard approaches, it is important to understand the differences in the protocols. Standard ChIP-seq protocols use crosslinked whole cells or nuclei without purification of crosslinked chromatin, and immunoprecipitation is typically performed with the ‘cleared’ suspension after extracts of sonicated crosslinked chromatin are centrifuged, thus removing most of the insoluble material. Even if this fraction is not cleared in this way, the insoluble material does not perform well in immunoprecipitation assays (*61*). Previous studies have identified SATB1 as a salt-extraction resistant (nuclear matrix) protein (*39, 40*). Thus, using canonical ChIP-seq approaches, SATB1-directly bound BURs in the inaccessible nuclear substructure would necessarily be lost in the insoluble fraction. Conversely any SATB1-associated genomic regions in open chromatin regions (direct or indirect), would become the predominantly captured fraction that is amplified during PCR-based library preparations. Thus, SATB1 directly bound to BURs in the high-salt extraction resistant nuclear substructure is likely missed by standard ChIP-seq (*53, 54*) and by CUT&Tag (*55*), which is also known to preferentially detect protein interactions in open chromatin regions. In contrast, in urea ChIP-seq, whole, intact crosslinked chromatin is purified through urea ultracentrifugation, first removing all indirectly bound proteins from chromatin. Subsequently, the entire urea-purified crosslinked chromatin is quantitatively solubilized. The two main advantages of urea ChIP-seq are that (1) only direct-binding profiles will be obtained for a protein of interest, and (2) chromatin tightly associated with nuclear substructures, which are typically discarded in the insoluble fractions in standard ChIP-seq, are retained as whole intact crosslinked genomic DNA in urea ChIP-seq. Therefore, all direct DNA binding profiles can be obtained, regardless of the original chromatin state (i.e. no bias toward open chromatin). Thus, urea ChIP-seq is uniquely suited to detect direct binding sites of SATB1 as well as other BUR-binding proteins genome wide. For non-BUR-binding proteins (e.g. CTCF and PRC2 core subunits), urea ChIP-seq generates identical DNA binding profiles as standard ChIP-seq. Although we detected some “phantom” or false-positive peaks using SATB1-KO cells by standard ChIP-seq (Table 3 and 4), under our experimental conditions, we confirmed that the majority of SATB1-binding peaks are SATB1-dependent. Importantly, these SATB1 peaks from standard ChIP-seq were essentially undetected by urea ChIP-seq, suggesting that SATB1 indirectly binds to these sites presumably by associating with nuclear proteins bound to their target sites in “open” chromatin. SATB1 is equipped with domains responsible for specific binding to BURs (*92-95*) as well as for associating with or recruiting chromatin binding proteins to specific genomic sites (e.g. CHRAC and ACF1 nucleosome mobilizing complex subunits, Brg1, p300, Hdac1, Gata3, Stat6, c-Maf, Pit1, and β-catenin) (*39, 42, 44, 50, 96-100*). Therefore, we anticipate that SATB1 binds indirectly and dynamically to regulatory regions in open chromatin by associating with many chromatin-modifying and transcription factors to form protein complexes to support tissue-specific gene expression. Despite the contrasting SATB1-binding profiles obtained from standard ChIP-seq and urea ChIP-seq, generation of the peaks from both methods require SATB1. Therefore, SATB1-mediated chromatin organization appears to have two components: one involves direct binding to BURs, while the other involves indirect binding to other chromatin-interacting proteins at genomic sites in accessible chromatin.

### Two-tiered chromatin organization mediated by SATB1

Considering that the two contrasting SATB1-binding profiles require SATB1, an important question emerges as to whether there is a functional or structural link between the two SATB1-bound chromatin regions. We conceive that SATB1 might have a role in bringing together chromatin structure formed by BURs tightly bound to nuclear substructure and a highly accessible “open” chromatin enriched in transcription related proteins. Our urea 4C-seq results suggest that these two regions are connected via SATB1-mediated interactions involving BURs. The ligation step in urea 4C-seq enabled us to capture chromatin sites that were in close spatial proximity to the bait site (e.g. a BUR) at the time of crosslinking. Apparently, 3D structures fixed by formaldehyde retained sufficient chromatin proximity after urea purification to enable capture of BUR-interacting fragments. The urea 4C-seq revealed that a SATB1-bound BUR site (BUR-1 as a bait), located at the border of a gene desert and a gene-rich region containing SATB1-regulated genes (*Rag 1* and *Rag2*), interacts extensively over a 5.8Mb gene-rich region with many gene loci, such as enhancers, promoters, BURs, and intergenic regions. Such ultra-long-distance interactions from BUR-1 predominantly cover the entire 5.8Mb gene-rich region, mostly avoiding the gene desert region. Such interactions were confirmed with another bait (BUR-2).

The SATB1-mediated chromatin interactions from BUR-1 and BUR-2 (validation control) as baits, covering the 5.8Mb *Rag1/Rag2*-containing gene-rich region, are tightly linked to SATB1-dependent expression of eight genes including *Rag1* and *Rag2* (P<0.05) out of 43 genes in this region in thymocytes. BUR-1 further interacts with far distal gene-rich regions *in cis* in chromosome 2 (>44Mb), skipping interstitial gene desert regions in a SATB1-dependent manner, albeit through weaker interactions (*Figure 5—figure supplement 3*). These chromatin interactions are at a much greater scale than those previously observed, such as T helper 2 cell activation induced dense chromatin looping in the 200kb cytokine gene cluster, including several BURs and regulatory regions, tightly linked to activation of the cytokine genes (*42*) and interactions at the subTAD levels (or <∼500Kb) between specific enhancers or enhancers to promoters in thymocytes (*53-55*). Combining results from SATB1-binding profiles derived from urea ChIP-seq and standard ChIP-seq as well as urea 4C-seq, a model is illustrated how the two chromatin regions might interact depending on SATB1 (*Figure 8*). In this model, the stable SATB1 binding to BURs in the SATB1-rich nuclear substructure forms a foundational chromatin scaffold and SATB1 tethers highly accessible “open” chromatin regions to this scaffold by binding to regulatory protein complexes—the SATB1 proteinaceous matrix would serve as a scaffold and interface. We anticipate that the latter interactions to be dynamic and form the highly interacting chromatin architecture over gene-rich accessible chromatin regions, bringing regulatory sequences in close vicinities. Both components of SATB1-mediated chromatin organization likely underlie regulation of gene expression. This two-tier model of SATB1 chromatin organization is highly reminiscent of previously published results on SATB1’s role in transcription factor Pit1-regulated gene expression in pituitary glands (*44*). SATB1 associates with Pit1 and β-catenin, and SATB1 is required for tethering Pit1-bound enhancers to the subnuclear architectural structure that is resistant to salt extraction (note: it is referred to as matrin-3 rich network, as matrin-3 is enriched in such substructure as well). A naturally occurring mutation of human PIT1 (R271W) that causes combined pituitary hormone deficiency fails to associate with SATB1 and β-catenin. Further studies with mutated R271WPit1 revealed that the tethering of Pit1-bound enhancers to the insoluble nuclear substructure is essential for effective activation of the Pit1-regulated transcriptional program.

**Figure 8.**
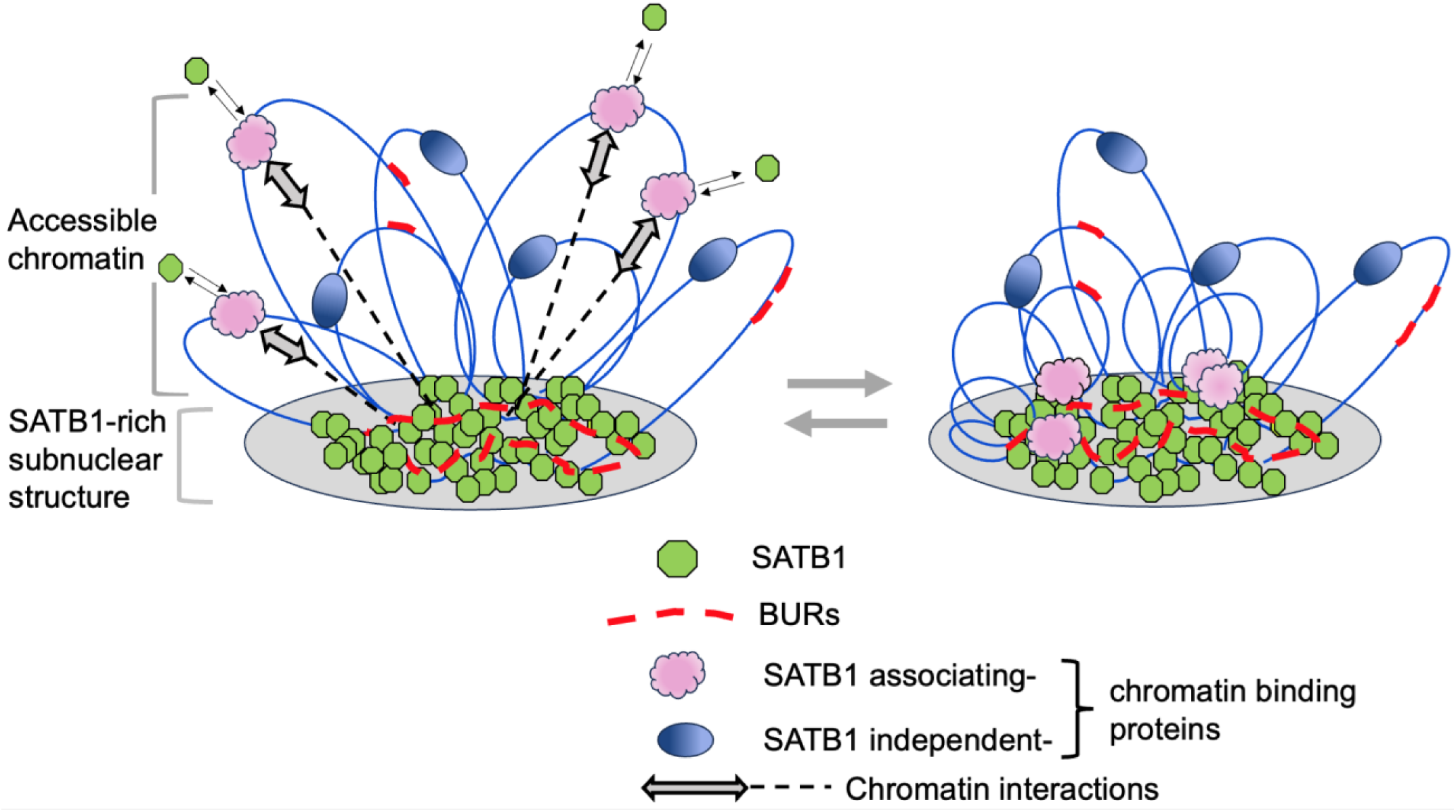
A model for SATB1-mediated chromatin organization with direct and indirect binding of SATB1 with chromatin. This model is based on the chromatin-binding profile of SATB1 obtained from urea ChIP-seq, which shows direct binding of SATB1 to BURs, and from standard ChIP-seq, which indicates presumably indirect binding of SATB1 to gene-rich, accessible chromatin regions. Additionally, the model incorporates chromatin interactions of selected SATB1-bound BURs detected by urea 4C-seq in this study. These BURs are mapped within accessible chromatin. Once BURs are bound by SATB1, they are tightly anchored to the SATB1 subnuclear structure, which resists high salt extraction, as demonstrated in previous studies (*39, 40*). We propose that the direct binding of SATB1 to BURs within the SATB1-rich subnuclear structure is stable and forms a chromatin scaffold. This chromatin scaffold interacts dynamically with active gene-rich regions through indirect binding, creating functional chromatin complexes (or hubs) that regulate gene expression depending on the cell type. Future studies are needed to determine the potential role of soluble SATB1 proteins and other regulatory factors in these interaction events.

Additional research will deepen our understanding of how SATB1 regulates cell-type specific gene expression. The Wnt signaling pathway plays pivotal roles in carcinogenesis and development (*101, 102*) and it will be important to investigate the role and nature of SATB1’s involvement in this pathway. Beta-catenin is the key component of the canonical Wnt signaling pathway; upon nuclear translocation, it associates with the TCF/LEF family of transcription factors to activate target genes. Because SATB1 also associates with beta-catenin, it is expected that SATB1 may influence target gene expression, similar to its effects on Pit1’s target genes. In addition, non-coding RNAs may have an important role in SATB1 chromatin organization and gene expression. Especially, specific long non-coding RNAs (lncRNAs) that remain associated with crosslinked chromatin after urea purification may deserve attention (unpublished results). Most non-coding RNA is associated with RNA-binding proteins (RBP) to form the ribonucleoprotein network which provides RNA structural scaffolds for the 3D chromatin organization and function (*103-106*). The SATB1 subnuclear architecture has a prominent cage-like distribution in thymocytes while neurons have a spider-web like distribution. It would be of interest to address whether RNA, including lncRNAs, plays a role in the SATB1-rich subnuclear proteinaceous scaffold formation depending on cell type and directing SATB1 to target specific sets of BURs in a cell function-specific manner.

### Lack of overlapping function for SATB1 and CTCF

Results obtained using urea-purified crosslinked chromatin show no relationship between CTCF and SATB1. Several pieces of evidence support this conclusion. First, genome-wide CTCF-binding profiles and Hi-C profiles remained largely unaltered by SATB1 deletion in thymocyte (*Figure 3*), a finding supported by other studies (*53, 54*). Second, by urea ChIP-seq, SATB1:BUR binding sites precisely exclude CTCF-binding sites, which contrasts with recent findings that SATB1 and CTCF-binding profiles overlap by standard ChIP-seq(*54, 55*) or by CUT&Tag (*55*).Third, chromatin interactions (>5.7Mb) from a SATB1-bound BUR are not constrained to individual TADs but span multiple TADs. Fourth, by urea ChIP-Western, SATB1 does not interact with CTCF or cohesin on chromatin. Fifth, CTCF does not bind to BURs. Therefore, chromatin organization mediated by SATB1 through direct binding to BURs is distinct from that mediated by CTCF and cohesion.

### Potential biological significance of BURs mostly enriched in LADs

We identify BURs as genomic elements enriched in LADs which are large chromatin domains located proximal to the nuclear lamina. LADs and LAD borders have critical roles in gene regulation (e.g. the *Tcrb* locus) (*2, 86*). Except for being generally AT-rich, genomic features of LADs have thus far been poorly characterized. BURs are not simply any AT-rich sequence, but rather, are characterized by a specific ATC sequence context. BURs are important genomic elements that are specifically targeted by SATB1 which has a role in cell-type specific gene expression (*41*). Thus, discovering BUR enrichment in LADs uncovers mechanistic insights into the biological function of LADs in gene regulation. Although LADs form heterochromatin proximal to the nuclear envelope, LAD organization is dynamic. While LADs are certainly constrained to the nuclear lamina, regions within LADs extend well into the nuclear interior and display dynamic interactions with the lamin proteins. In addition, LADs are dynamic during differentiation or other cell-state changes. For instance, during differentiation of mouse cells, hundreds of genes display dynamic intranuclear spatial repositioning relative to the nuclear lamina (*107*). Some LAD regions dynamically contact the nuclear lamina, and gene movement in and out of the nuclear periphery in a circadian rhythm has been reported (*108*). Recently, relocation of the *hunchback* gene in *Drosophila* neuroblasts to nuclear lamina has been shown to require polycomb proteins binding to a specific intronic element of the gene locus. This process is important for heritable gene silencing to regulate neuroblast competence (*109*). The organization of genomic DNA at the lamina has been linked to both inactive chromatin state and the nuclear lamins, along with other nuclear membrane proteins (*2, 110-115*). While chromatin state is likely important for both positioning at and de-localization from the lamina, little information is available concerning *in vivo* mechanisms specifying which regions within LADs could be targeted for de-localization and subsequent cell-type specific gene activation. The SATB subnuclear architecture is found mostly in the nuclear interior of thymocytes where many individually cloned BURs have been detected by DNA-FISH. Among such interiorly localized BURs, a BUR 5’ of *Myc* was confirmed to co-localize with SATB1 by immunoDNA-FISH in thymocytes (*39, 40*). A closer look at immunostaining profiles of SATB1 distribution in thymocyte nuclei, however, shows that a small reproducible fraction of SATB1 resides in the peripheral zone of nuclei and in some cells the tip of the SATB1 network extends to and touches this region of the nucleus (*39*).

Future studies are needed to investigate the spatial context of a subset of BURs and LADs in relation to SATB1 subnuclear architecture, depending on the cell type. For instance, many SATB1-bound BURs are identified in the *Gabrg1* and *Gabra2* loci in neurons, but not in thymocytes (Fig. 6B), even though thymocytes express SATB1 at high levels. Therefore, studies on nuclear positioning of select BURs in neurons, thymocytes, and in non-SATB1-expressing cells would provide valuable insights. In addition to LADs, transcriptionally active gene-rich regions outside LADs also contain BURs, albeit at a considerably lower density compared to LADs (e.g. Rag1/Rag2, Foxp3, Runx3 loci). Some of these BURs located near enhancers are bound by SATB1. It is of interest that the number of BURs targeted by SATB1 greatly varies depending on cell type, where neurons have by far the greatest number of BURs captured by SATB1. This may reflect the level of plasticity for gene transcription or number of genes that must be dynamically regulated in each cell type in which SATB1 is expressed. There is growing evidence supporting the critical roles of chromatin topology and nuclear architecture in neuronal response to external stimuli to establish precise expression program associated with cognition (*116*). In fact, SATB1 has roles in early postnatal cortical neuron development and neuronal connectivity (*37, 117*). The exceptionally large number of BURs directly interacting with SATB1 in cortical neurons may reflect specialized function of neurons that might require highly dynamic transcriptional regulation to enable rapid response to a wide variety of environmental stimuli. How SATB1 selects specific subsets of BURs and if and how SATB1’s selection of BURs contributes to cell-type specific gene expression are important topics for future research.

### A broader significance of BUR-binding proteins beyond SATB1

Importantly, SATB1 is not the only BUR-binding protein, indicating the broader biological significance of BURs as genomic elements. Although SATB1 confers by far the highest affinity to BURs, several other nuclear factors (e.g. nucleolin, HMGI/Y, PARP1, Ku70/86 heterodimer) have also been verified to directly and specifically bind BURs *in vitro* (*84, 118, 119*). Recently, we identified Tip5/Baz2a as an additional strong BUR-direct binding protein from mouse embryonic stem cells (mESCs) that is essential for pluripotency (submitted for publication). Similar to SATB1, this nuclear protein also binds specifically to BURs in mESCs, and identification of BURs as its direct targets in mESCs requires urea ChIP-seq as well. Despite the identical BUR binding specificity of SATB1 and this stem cell factor, these two proteins confer entirely different functions in distinct cell types. Therefore, BURs may serve as core genomic marks that can be selectively targeted by distinct BUR-binding proteins to form unique 3D chromatin regulatory network depending on the BUR-binding protein for specific cell functions. BURs in the mammalian genome may have similarities in function with the “tethering elements” in *Drosophila* (*120, 121*).These elements can be bound by regulators such as pioneering factors and polycomb proteins and were shown promote interactions of regulatory regions up to ∼250kb within TADs to regulate transcription of genes. Most recently, meta-loops constituting a meta-domain spanning numerous TADs in CNS of *Drosophila* were reported (*122, 123*). Although meta-loops are similar in size with SATB1-mediated chromatin interactions, the CNS meta-loops have different features. They are formed by transcription factors (e.g. CTCF and GAF), the half of them is anchored at the TAD borders, and the metadomain boundaries are CTCF-bound. It appears that clusters of very long chromatin looping events linked to gene expression may form by multiple mechanisms across species. Future research on BUR-mediated genome organization regulated by distinct BUR-binding proteins will likely provide more in-depth understanding about cell-type specific mechanisms driving 3D chromatin architecture that contributes to major changes in gene expression and cell phenotypes.

## Materials and Methods

### Experimental design

#### Primary cells

All animal studies were performed under Institutional Animal Care & Use Program at University of California San Francisco for collecting brain and thymus and the Boston University for collecting skin. Thymi were isolated from 2 weeks-old *Satb1*^+/+^ and *Satb1*^−/−^ mice, as well as from *Satb1*^+/+^:*Cd4*-cre and *Satb1* ^F/F^:*Cd4*-cre mice. Brain was isolated from 6-10 weeks-old *Satb1*^+/+^ mice. Thymocytes were prepared from thymi and cortex was excised from brain for this study. Skin was isolated from postnatal day 0.5 of C57Bl/6 mice.

#### Antibodies

Antibodies used and for their purposes are the followings. CTCF for ChIP-seq: Active MOTIF (61311. lot# 34614003), CTCF for Western: Abcam (ab129973) and Millipore (07-729; lot#-3429114), RAD21 for ChIP-seq and Western: ABclonal (A18850, lot# 3521809001), SATB1 for ChIP-seq and Western: rabbit polyclonal 1583 prepared by Kohwi-Shigematsu’s laboratory and Abcam (ab109122, lot#GR137720-13) and Histone H3 for Western: Abcam (ab1791, lotGR3236305-2).

#### Urea-purified chromatin immunoprecipitation and urea ChIP-seq

Thymocytes and single cell suspension prepared for cortex and skin were cross-linked with 1% formaldehyde for 10 min at room temperature, followed by addition of glycine at 0.125 M for 5 min to quench cross-linking reaction. These cells were washed with 1X PBS, suspended in 1% BSA-PBS and cells were precipitated in a tube. These cells were re-suspended in lysis buffer (4% SDS, 50 mM Tris-HCl [pH 8.0], 10 mM EDTA, 100 mM NaCl) with Protease inhibitor (Roche) and RNase inhibitor. The resulting cell lysate was incubated for 30 min at room temperature and loaded on top of 8 M urea followed by ultracentrifugation in a Beckman TL-100 Ultracentrifuge for 4-5 hrs at 45K rpm (181,000g, g=relative centrifugation force or RCF; TLS55 rotor). Recovered chromatin pellet was sonicated to generate fragment size ranging 300–1,000bp using Branson sonifier (20% power, 10 sec on and 30 sec off for 4-6 cycles) in 300μl of sonication buffer [0.5% Na-laurylsarcosine, 0.1% deoxycholate (DOC), 1mM EDTA, 0.5mM EGTA, 10mM TrisHCl, 100mM NaCl] with Protease inhibitor and RNase inhibitor. Note: urea-purified crosslinked chromatin is highly vulnerable to sonication, therefore, it needs to adjust the sonication condition depending on cell type and/or the size of the precipitate DNA. The sonicated chromatin with 1% Triton X-100 were dialyzed against dialysis buffer (10 mM Tris-HCl [pH 8.0], 100 mM NaCl, 1 mM EDTA, 5% glycerol), and dialyzed for 16 hrs at 4°C (this dialysis is optional). The chromatin sample was centrifuged at 15,000g for 5 min at 4°C to remove a trace amount of residual insoluble components. The test-reverse cross-linking of chromatin aliquot (5-10μl of the final volume) was done at 65°C for 2-3 hrs in buffer containing 1% SDS, 100 mM NaCl, 1 mM EDTA, 10mMTris. Samples were further treated by 100 µg/ml RNase A at 37°C for 1 hr, 200 µg/ml proteinase K at 60°C for more than 3 hrs, followed by DNA purification using QIAquick PCR purification kit (Qiagen), and shearing efficiency was examined by Bioanalyzer 2100 (Agilent Technologies Inc.). For chromatin-immunoprecipitation, 1 µg of an antibody of choice were added to 10 ∼ 20 µg of sonicated solubilized chromatin in 350 μl in dilution buffer (1% Triton X-100, 0.01% SDS, 1 mM EDTA, 15 mM Tris-HCl, 100 mM NaCl) and incubated overnight with rotation at 4°C. At least three chromatin samples were prepared for ChIP-seq from independent experiments to confirm the results. The chromatin fragments bound to antibodies (after incubation overnight) were captured using 20 µl of protein-A, protein-G or protein AG Dynabeads (Invitrogen; pre-washed with 0.1% BSA [Fraction V, Sigma), by 4 hrs incubation time at 4°C. Dynabeads bound to chromatin fragments were washed twice with Binding buffer (10 mM Tris, pH8.0, 1 mM EDTA, 1 mM EGTA, 0.5% Triton X-100, 0.1% DOC, 100 mM NaCl), twice with LiCl buffer (100 mM Tris [pH 8.0], 1 mM EDTA, 250 mM LiCl, 0.5% NP-40, and 0.5% DOC). The tubes were changed and washed twice with 0.1M NaCl-TE buffer. The immunoprecipitated samples were eluted from Dynabeads by elution buffer (10 mM Tris-HCl [pH 8.0], 1 mM EDTA, 100 mM NaCl and 1% SDS) at 65°C for 20 min. After elution, ChIP DNAs was reverse-crosslinked at 65°C for 6 hrs, followed by RNase-A treatment at 37°C for 1 hr and proteinase K treatments for 3 hrs at 60°C. The recovered DNA was further purified with a QIAquick PCR purification kit (Qiagen), with elution in 15µl of Qiagen Elution Buffer (alternatively, phenol-chloroform extraction and ethanol precipitation). Briefly for library construction, ChIP DNA fragments were end-repaired and ligated to indexed sequencing adapters (NEB or Rubicon kit) and amplified with 8-11 rounds of PCR, depending on the DNA amount. Using SPRI beads (Beckman coulter), small fragments (<200bp) were removed from the amplified library. The size-selected library was sequenced to 50bp single-end read lengths on an HiSeq4000 (Vincent J. Coates Genomic Sequencing Laboratory, UC Berkeley).

#### Standard ChIP-seq

For standard ChIP-seq, the protocol described by Kitagawa et al was employed (*47*).

#### Urea Chromatin immunoprecipitation (urea ChIP)-Western analysis

Urea-purified chromatin prepared as described above (∼10μg of chromatin-DNA) was Immunoprecipitated with indicated antibody to produce urea ChIP samples, and the chromatin fragments bound by the antibody was trapped by Protein-A beads. The antibody-bound chromatin fragment-trapped beads were washed with binding buffer/washing buffer/NaCl-TE (total 6 times), and the chromatin fragments were released and isolated from beads (65°C for 20 min) in 100 μl of 1% SDS, 10 mM Tris, pH7.6, 1mM EDTA, 100mM NaCl solution. The isolated chromatin fragments were subjected to reverse-crosslinking by incubation at 65°C for additional 6 hrs. The released proteins were dissolved in the Laemmli buffer, boiled and subjected to Western blot assay. To compare multiple samples, we confirmed that we used a similar amount of original urea-purified crosslinked chromatin before ChIP by subjecting each chromatin sample to Western blot using anti-H3 antibody. The similar amount of urea-purified crosslinked chromatin was then used for ChIP-Western studies.

#### Urea 4C-seq

Urea-purified crosslinked chromatin (described above for its preparation) was processed for urea-4C-seq basically as described(*66*) with some modifications. For the current experiment, the purified crosslinked chromatin (appears as a transparent pellet at the bottom of a centrifugation tube) after urea centrifugation was briefly sonicated in 100μl of the *Hind*III digestion buffer with Branson sonifier (20% power, 10 sec, once). Next, *Hind*III was added to digest chromatin (10-20 units/μg DNA) overnight at 37°C. *Hind*III was heat inactivated at 80°C for 20 min, ligation was performed in 1.0 ml with 1X ligation buffer with ligase (600 units) at 16°C for overnight, followed by reverse-crosslinking at 65°C for overnight. The DNA was then purified by phenol-chloroform extraction or by spin column (NucleoSpin, Macherey-Nagel). The purified DNAs (1-4μg) was further digested with *Hae*III (100-125 units) overnight. The *Hind*III-*Hae*III digested DNA fragments were treated with Klenow. The resulting blunt-end *Hind*III-*Hae*III DNA fragments were ligated with our custom double-stranded (ds) Linker DNA with one strand-Biotin modified,

Linker-top strand sequence with Biotin modified (marked by *):

5’-aattcggtacctctagaga**t\***a**t\***ccgatcgctcgag(XhoI)aagctt(*Hind*III)-3’;

Linker-bottom strand sequence: 5’-aagcttctcgagcgatcggatatctctagaggtaccgaatt-3’ as described (*66*). The Linker-ligated dsDNA fragments were purified by spin column to remove free Linker. The Linker-ligated dsDNA fragments were further digested with *Xho*I at the linker site (next site from *Hind*III). Single stranded DNAs were isolated from the *Xho*I digested DNA fragments by coupling to StreptAvidin Dynabeads (Invitrogen, Dynabeads M-280 streptavidin) followed by incubation in 40μl of 0.15M NaOH for 10 min at 25°C, followed by neutralization. Only single-stranded DNA, containing biotin coupled with streptavidin, were retained on the Dynabeads, releasing the non-biotin modified single-stranded DNA (single strand DNA capture) (*66*). The released non-biotin modified single strand DNAs were amplified by PCR (AmpliTaq Gold system, Applied Biosystems); 10 cycles at 94C for 30”, 55°C for 60” and 72°C for 60”, followed by 15-20 cycles at 94°C for 30”, 65°C for 60”, 72°C for 60”, using the custom Illumina sequence-bait oligomers and Illumina sequence-linker oligomer for library preparations as described below. The PCR amplified DNAs were purified by high and low size selection by beads (SPRIselect, Beckman Coulter). The purified DNAs were analyzed by Bioanalyzer to confirm their average size of ∼300-500 bp with mostly uniform distribution, followed by the Illumina sequencing using custom bait-specific sequence oligomers and Linker specific barcode oligomer. We used two bait sequences: 5’ gaagggggtaggaaagaaagctt 3’ (BUR-1 bait: adjacent to a SATB1-bound BUR, located1.4 kb from ASE) and 5’ cgactcattcctactgaagctt 3’ (BUR-2-bait: enhancer located in the second intron of B230118H07Rik close to a SATB1-bound BUR), both including a *Hind*III site.

The library was constructed by amplification using either Illumina sequence-BUR-1-bait or Illumina sequence-BUR-2-bait primer with Illumina sequence-linker oligomer. (Capital letters represent Illumina sequence, lower case letters represent bait sequences or the linker sequence)

Illumina sequence-BUR-1-bait oligomer:

5’-AATGATACGGCGACCACCGAGATCTACACTCTTTCCCTACACGAC-gaagggggtaggaaagaaagctt

Illumina sequence-BUR-2-bait oligomer:

5’-AATGATACGGCGACCACCGAGATCTACACTCTTTCCCTACACGAC-cgactcattcctactgaagctt

Illumina sequence-linker oligomer:

5’-CAAGCAGAAGACGGCATACGAGAT (barcode; e.g. CGTGAT) GTGACTGGAGTTC-gcgatcggatatctctagaggtaccgaattc

To validate successful experiments, we confirmed that no PCR amplified product was generated by using either the bait oligomer or linker oligomer alone.

For the library construction, Taq-Gold polymerase (NEB) was used for PCR by the following condition: 94°C for 5-10min (one cycle), 94C for 30 seconds, 55C for 30 seconds, 72C for 60 seconds (for 5-10 cycles), 94°C for 30 second, 65°C for 30 seconds, 72°C for 60 seconds (for 20 cycles), followed by 72°C for 10 min. PCR products were purified by phenol-chloroform extraction twice, CHCl3 extraction once, followed by EtOH precipitation (Optional; PCR purification by spin column, Qiagen).

The 4C libraries were sequenced by Illumina-HiSeq4000, using BUR-1-bait sequence and BUR-2-bait sequence as custom sequence oligomers as follows:

BUR-1-bait sequence oligomer: 5’-CTTTCCCTACACGAC-gaagggggtaggaaagaaagctt

BUR-2-bait sequence oligomer: 5’-ACTCTTTCCCTACACGAC-cgactcattcctactgaagctt

Each library sequence was identified with Linker specific barcode oligomer;

5’-gaattcggtacctctagagatatccgatcgc-GAACTCCAGTCAC. (Capital letters represent Illumina sequence and lower-case letters represent linker sequence). For sequencing, the above custom barcode sequence oligomer was required instead of the common Illumina barcode sequencing oligomer.

#### Hi-C

Hi-C was performed as described (*17*). Primary thymocyte samples from *Satb1*^+/+^:*Cd4*-cre and *Satb1* ^F/F^:*Cd4*-cre mice were formaldehyde crosslinked at 2% at room temperature for 10 min. Two biological replica per sample were prepared for Hi-C. Hi-C libraries was prepared using the Arima HiC^+^ kit (Arima Genomics) exactly following the associated protocol. Paired-end sequencing was performed by Illumina NovaSeq 6000.

### Statistical analysis

#### Significance of overlap of SATB1 with BURS

We used the *bootRanges* function that is part of the *nullranges* bioconductor package (*71, 124*) It was used to compute the significance of overlap of two sets of genomic regions (e.g., BUR versus the called peaks for each of the ChIP-seq data sets). This method is based on subsampling genomic regions of a given size from within genomic segments of the same stateHere the mouse genome mm9 was segmented into regions capturing 3 states based on density of known genes using the segmentDensity function using the CBS method 4 (*125*) and assuming a segment length of 2Mb. Regions represented blacklisted regions (queried using the terms “excluderanges” and “mm9”) in the AnnotationHub package were excluded from the analysis. For each comparison [reference regions (known BUR peak calls) versus query regions (SATB1 Urea or Standard Chip-seq peak calls)], the observed number of overlapping regions is first computed, 1000 sets of random query region peak calls were sampled given the estimated genome segmentation using blocks of length 300kb, the overlaps for each of these 1000 random query regions with the reference regions were then computed and finally, the estimated p-value is based on the fraction of 1000 random overlaps greater than or equal to the observed number of overlaps. These results were validated by evaluating the statistical enrichment of SATB1 binding at BURs using the regioneR package (v1.30.0) in R (*126*), generating a P value of 0.000999, Z-score of 543.37, and a one-sided test confirmed greater than expected overlap. SATB1-bound genomic regions (satb1_gr) and BURs (bur_gr) were represented as GRanges objects. To determine whether SATB1 binding events occur more frequently at BURs than expected by chance, we performed a permutation test using permTest with 1,000 iterations. SATB1 regions were randomly shuffled across the mm9 mouse genome using randomizeRegions, while excluding masked regions such as repeats and assembly gaps via the built-in soft masking available in the BSgenome.Mmusculus.UCSC.mm9 reference. Enrichment was evaluated using numOverlaps to count the number of overlapping SATB1–BUR events in each permutation. A one-sided test was used to assess whether observed overlaps were significantly greater than the randomized distribution.

For genome-wide heatmaps we quantified SATB1 chromatin occupancy relative to mapped base unpairing regions (BURs). BURs were provided in narrowPeak format, and the SATB1 ChIP-seq peaks were obtained from a processed peak file containing signal intensity. Using the normalizeToMatrixfunction from the EnrichedHeatmap package (v1.30.0)(*127*), we computed SATB1 signal intensity in 50 bp bins across a ±3 kb window centered on each BUR. BURs were either plotted as a total genome-wide dataset or stratified into lamina-associated domains (LADs) and inter-LADs (iLADs) based on overlap with DamID-defined LAD intervals using the LADetector script(*86, 88*). Normalized signal matrices were used to generate heatmaps of SATB1 enrichment using ComplexHeatmap, with LADs and inter-LADs plotted separately. Line plots above the heatmaps show the average SATB1 signal across all BURs in each category. Color scaling was applied using colorRamp2 from the circlize package (v0.4.15)(*128*). All visualizations were constructed using ComplexHeatmap (v2.18.0), with rasterization enabled for performance.

To assess the alignment of BURs to SATB1-bound loci, we performed the inverse analysis by quantifying BUR signal intensity around SATB1 binding peaks. SATB1 peaks were split into LAD- and iLAD-associated groups using genomic overlap with LAD intervals. BURs were used as a quantitative signal (score column from the narrowPeak file), and enrichment around each SATB1 site was computed in 50 bp bins across a ±3 kb window using normalizeToMatrix function from the EnrichedHeatmap R package (v1.30.0). The resulting matrices were visualized as heatmaps and line plots of average BUR signal genome-wide or in each domain context (LAD vs iLAD). All visualizations were constructed using ComplexHeatmap (v2.18.0).

#### Hi-C data processing

Hi-C datasets were analyzed using the HiC-bench platform (*129*). The application of this method is described in detail(*130*). Briefly, data were aligned against the mouse reference genome (mm10) by bowtie2 (version 2.3.1)(*131*) with mostly default parameters (specific settings: --very-sensitive-local --local). For Hi-C, aligned reads were filtered by the GenomicTools(*132*) tools-hic filter command integrated in HiC-bench. Interaction matrices for each chromosome separately were created by the HiC-bench platform at a 10kb resolution. The Hi-C filtered contact matrices were normalized using “iterative correction and eigenvector decomposition” (ICE) algorithm (*133*) build into HiC-bench. To account for variances of read counts of more distant loci, distance normalization for each matrix was performed as recently described(*134*). TADs were called using the algorithm developed within HiC-bench(*129*) at 10kb bin resolution with the insulating window of 100kb. For visualization of Hi-C data, we generated HiC files using the Juicer(*135*).

#### Quality Control

The total numbers of reads per biological replicate for each genotype ranged from 200 to 230 million reads. The percentage of reads aligned was >98% in all samples. The percentage of accepted reads (ds-accepted-intra” and “ds-accepted-inter”) was in the range of 50% to 55.5%. The absolute number of accepted intra-chromosomal read-pairs were ∼90-100 million reads for each sample. We analyzed data from individual samples as well as combined data from two replicates (∼200 million reads) for each genotype.

Scaling plot was performed by aggregating the counts per 10kb distance bins. Values were cpm-normalized by dividing by the total number of valid intra-chromosomal pairs.

#### Compartments

Compartment analysis was performed using the Homer pipeline (v4.6)(*136*). Homer performs a principal component analysis of the normalized interaction matrices and uses the PCA1 component (cis-Eigenvector 1) to predict regions of transcriptionally active (A compartments) and transcriptionally inactive chromatin (B compartments). Homer works under the assumption that gene-rich regions with active chromatin marks have similar PC1 values, while gene deserts show the opposite (http://homer.ucsd.edu/homer/interactions/HiCpca.html). HiC filtered matrices were given as input to Homer together with a list of house-keeping genes for compartment prediction. The house-keeping genes coordinates are used by Homer as prior information of active regions. We generated density plots to compare the cis-eigenvector 1 values of the knock-out and wild-type samples with 10kb genomic bins. Pearson correlation coefficients were calculated (‘Cor’ function in R) between the counts for every pair of 10kb bins on chromosome 2. Heatmaps representing the Pearson correlation of interactions of chromosome 2 for WT and KO thymocytes are displayed by juice box (500kb resolution).

#### urea ChIP-seq analysis

Sequenced reads were quality-checked using FastQC prior to downstream analysis. Unaligned reads were mapped to the mouse reference genome (NCBI37/mm9) using bowtie2 (version 2.3.1)(*131*) allowing for three mismatches in the seed region and only unique matches. Candidate binding sites were called using MACS2 (v2.2.7)(*137*) using a q-value threshold of 0.01. Coverage tracks are presented in RPM (read pileups <u>p</u>er <u>m</u>illion reads) and utilities from UCSC were used for creating browser tracks (*138*). To determine intersection of peaks across ChIP-seq samples, deepTools(*83*) was used.

#### urea 4C-seq analysis

Reads were mapped with Bowtie, then assigned to *HindIII* sites with the same utility tools used in (*77*). These same tools also normalized the counts to per million intrachromosomal reads and removed reads to restriction fragments comprising more than 2% of the total reads (namely the contiguous adjacent restriction fragment corresponding to undigested chromatin during the urea 4C, and a rare (<5) numbers of fragments with extreme PCR duplications). The generated bedGraph files and these were converted to bigWig format using the bedGraphToBigWig program for direct visualization of the peaks on the UCSC browser.

#### Analysis of SATB1 bound sites and other genomic features

LADs were identified as genomic intervals using the previously described LADetector algorithm (https://github.com/thereddylab/pyLAD) or sequencing data (https://github.com/thereddylab/LADetector) (*114, 139*).SATB1 urea-ChIP data from thymocytes were tested for statistically significant overlap of LADs using the GenometriCorr package(*140*). The GenometriCorr package applies multiple spatial tests of independence including a relative distance test (i.e., if two elements such as LADs and SatB1 sites are spaced more often than expected at a certain relative distance from each other), projection test (i.e., testing for significant overlap between these two positions assuming they represent single points points), and the Jaccard test (testing for correlation of these two positions assuming that they are not points but instead occupy some interval of the genome). To generate genome wide line plots of averages of intensities (relative score) of DamID and ChIP-seq peak data were generated over inter-LAD (10% of the size of the adjacent LAD)/border flanked LADs scaled to the same relative size using the Genomation R package(*141*). The average intensities were then normalized to fit a scale of −1 to 1 and plotted over the LADs (normalized relative score). To plot genomic features in the vicinity of enhancers, H3K4me1 peaks outside of LADs were filtered by subtracting away peaks that intersected LADs using the BEDTools (*142*). To plot genomic features in the vicinity DiPS in LADs, regions within LADs and DiPS were identified by LADetector. DiPS were then mapped as intervals and the method described above was applied to DiPS versus SATB1, CTCF, and chromatin modifications. Genome-wide averages of the respective ChIP-seq/DamID signal data were then generated within a +/− 3kb window centered at H3K4me1 peaks, using Genomation R package and plotted on a vertical scale ranging from −1 to 1.

## Acknowledgements

We thank Dr. Aristotelis Tsirigos, Ziyan Lin, and Javier Rodriguez Hernaez at New York University School of Medicine for Hi-C and intersection analyses, Dr. Anthony Schmitt from Arima Genomics for help with Hi-C experiments. We also thank Dr. Elphege Nora for reviewing Hi-C results and providing critical comments on the manuscript and Ms. Bao Ho for providing technical assistance. We thank Dr. Minoree Kohwi for critical reviewing of the manuscript and Dr. Peter Krijger for kindly testing our samples and Dr. Elzo de Wit for discussion on the 4C-seq analysis.

## Author contribution

Conceptualization: Terumi Kohwi-Shigematsu and Yoshinori Kohwi

Method development and validation: Terumi Kohwi-Shigematsu and Yoshinori Kohwi

Data production: Yoshinori Kohwi, assisted by Mari Grange

Formal analyses and bioinformatic analyses: Terumi Kohwi-Shigematsu, Karen L. Reddy, Xianrong Wong, Thomas Sexton, Hunter W. Richards and Ya-Chen Liang (Karen L. Reddy and Xianrong Wong performed all bioinformatic and statistical analyses on LADs and Thomas Sexton conducted data processing and bioinformatic analysis for the urea 4C-seq data)

Supervision: Terumi Kohwi-Shigematsu, Yoshinori Kohwi and Karen L. Reddy

Materials: Ichiro Taniuchi provided thymocytes from *Satb1*^F/F^: *Cd4*-*cre and Satb1*^+/+^: *Cd4*-*cre* mice for the Hi-C study. Yohko Kitagawa and Shimon Sakaguchi provided modified histone H3 ChIP-seq data for different stages of thymocyte development. Vladimir A. Botchkarev provided mouse skin for the urea ChIP-seq studies.

Funding acquisition: Terumi Kohwi-Shigematsu, Vladimir A. Botchkarev and Cheng-Ming Chuong

Writing—original draft: Terumi Kohwi-Shigematsu, Karen L. Reddy and Yoshinori Kohwi

Writing—review, editing and/or discussion: Terumi Kohwi-Shigematsu, Karen L. Reddy, Xianrong Wong, Yoshinori Kohwi, Ichiro Taniuchi, Thomas Sexton, Yohko Kitagawa and Shimon Sakaguchi

## Data availability

All sequencing data for ChIP-seq and Hi-C in this study was uploaded to NCBI Gene Expression Omnibus (GEO) and is available under the accession GSE191146.

## Figure Supplements and Supplementary Tables

**Figure 2—figure supplement 1.**
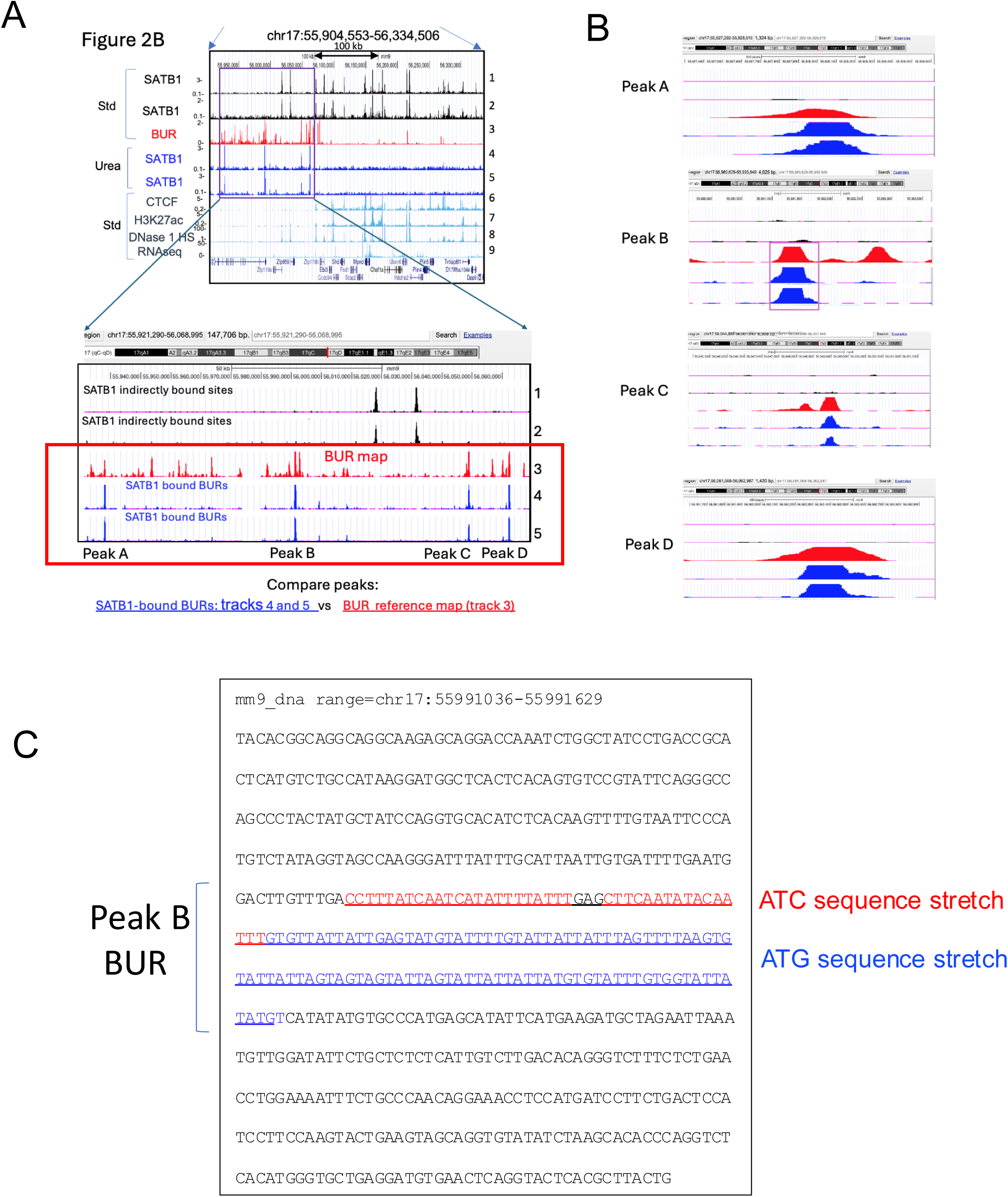
Zoomed-in view of Figure 2B showing precise coincidence of SATB1 urea ChIP-seq peaks with a subset of BURs. A) Peak A, Peak B, Peak C, and Peak D shown in tracks 4 and 5 overlap with BURs in track 3. The bottom panel is a zoom-in view of the top panel (from Figure 2B). B) High-resolution views of each peak region showing precise co-mapping of BUR (red peak) and SATB1 peak (blue peak). C) A representative BUR sequence (a cluster of ATC/ATC sequences) from Peak B.

**Figure 3—figure supplement 1.**
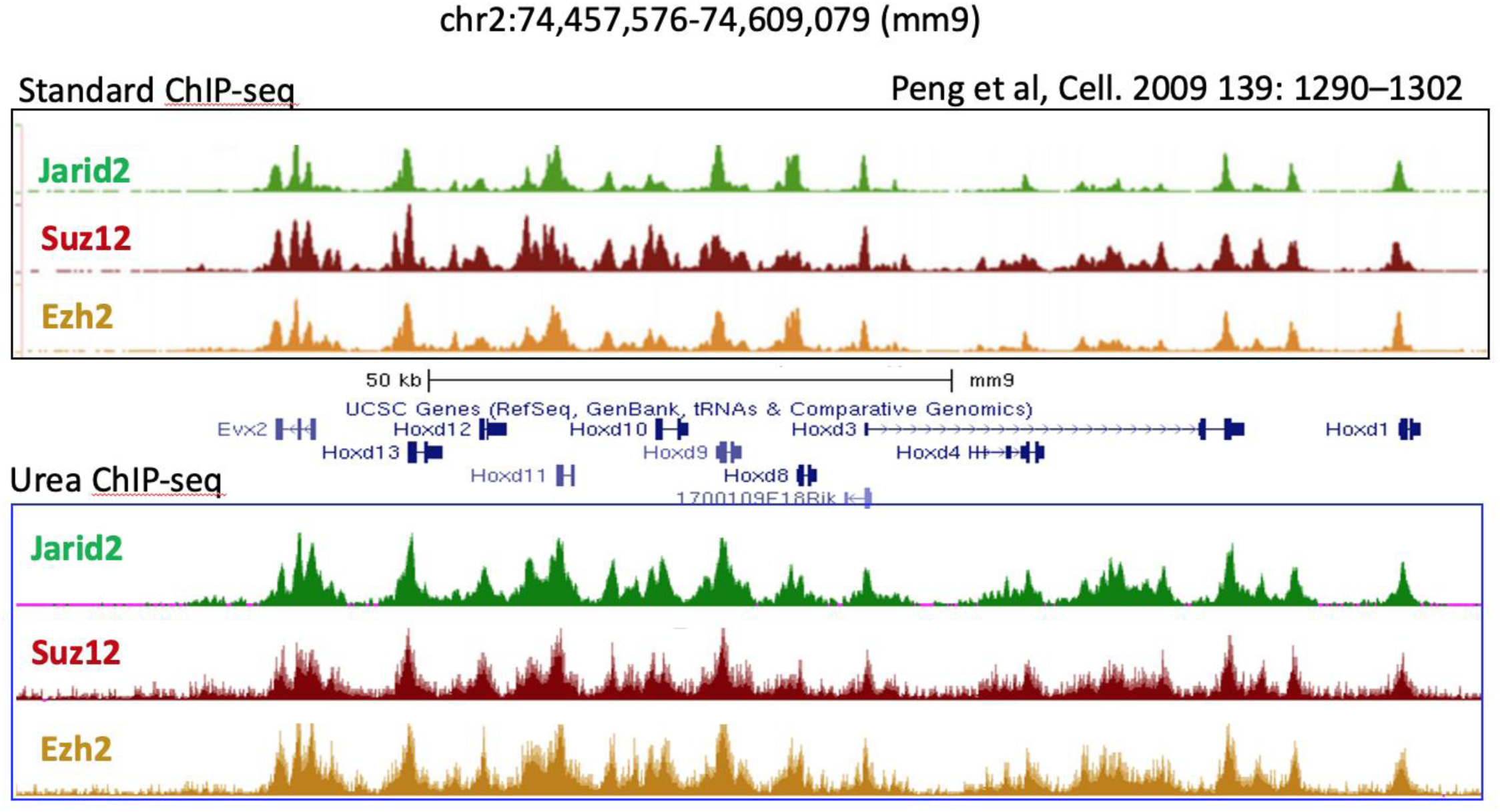
Similar DNA binding profiles for PRC2 core subunits obtained by standard ChIP-seq and Urea ChIP-seq. The DNA binding profiles of PRC2 core subunits, Jarid2, Suz12, and Ezh2 obtained by standard ChIP-seq (Ref 73) [Peng et al., Cell, 2009, GEO repository (GSE18776)] were compared against the DNA binding profiles of these proteins by urea ChIP-seq. Their binding profiles in the *Hoxd* cluster region on chromosome 2 (mm9) are shown. Their DNA binding profiles by urea ChIP-seq and by standard ChIP-seq are very similar.

**Figure 3—figure supplement 2.**
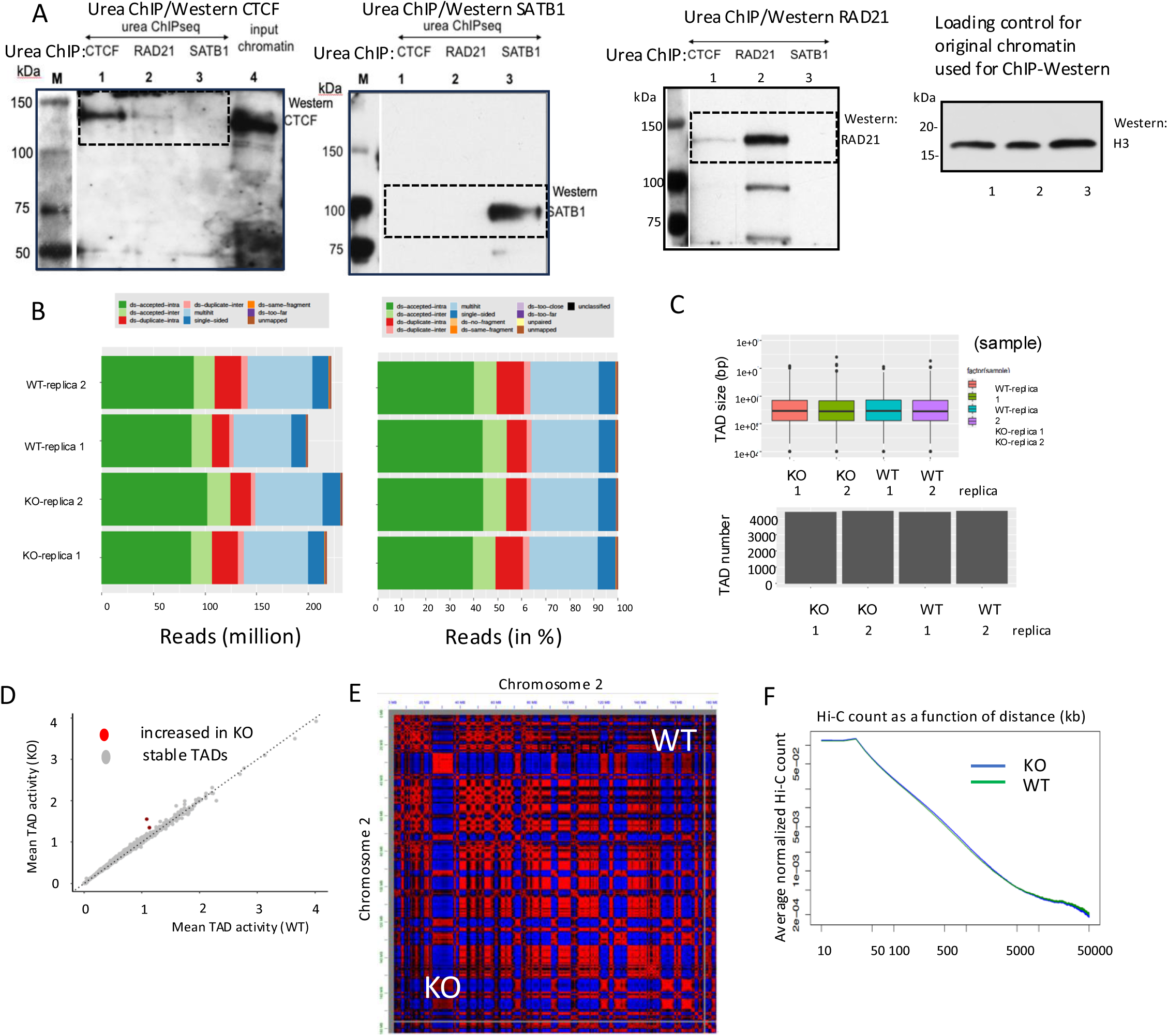
SATB1 does not directly co-bind chromatin with CTCF and RAD21 or has a role in TAD formation. **(A)** Urea ChIP-Western studies for CTCF, RAD21 and SATB1. Original data for Figure 2C is shown. The excised areas are indicated by dotted boxes (first three panels). Western blot with the anti-H3 antibody shows that the original urea-purified chromatin samples before ChIP were similar in amount (right panel). **(B)** Read alignment statistics for Hi-C datasets, as absolute reads (left) and relative reads (in %, right). “ds.accepted.intra” are all intra-chromosomal reads used for all downstream analyses. **(C)** Distribution of TAD sizes and number of TAD calls per sample using hic-ratio TAD caller at 10kb bin resolution and the insulation score of 100kb windows. **(D)** Scatter plot of pair-wise comparisons of TAD activities (the sum of normalized Hi-C chromatin contacts inside the TAD in WT) between WT and KO thymocyte samples, showing well-conserved intra-TAD activity in KO thymocytes. **(E)** The Pearson correlation estimates computed between the counts for every pair of 10kb bins on chromosome 2 shows that WT (control: upper right triangle) and KO (observed: lower left triangle) thymocytes have no difference in compartmentalization. The compartment A (red) and compartment B (blue). **(F)** The scaling plots displaying how the Hi-C contacts decrease as a function of genomic distance.

**Figure 4—figure supplement 1.**
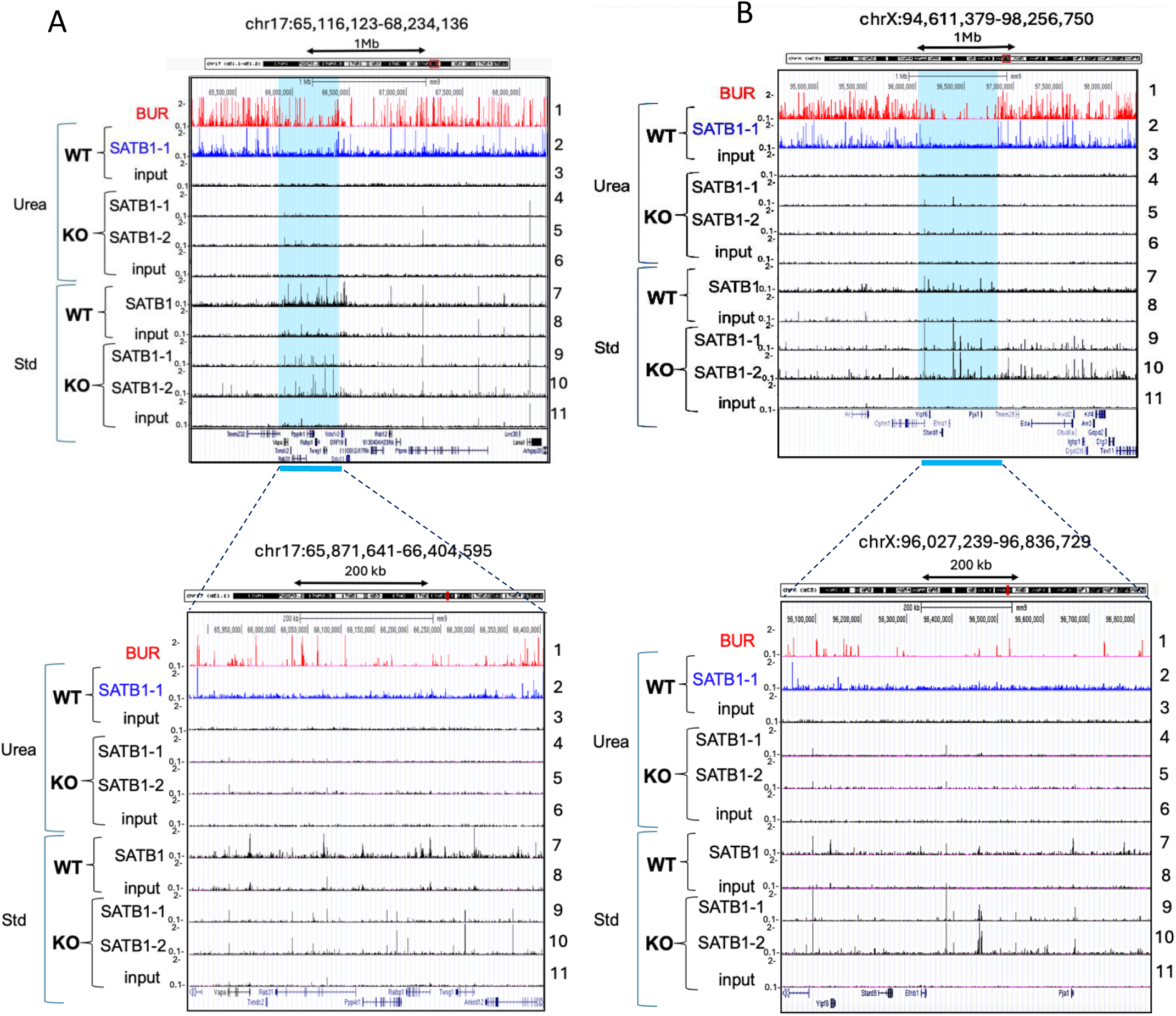
Standard ChIP-seq for SATB1 produces non-specific peaks in *Satb1*^−/−^ thymocytes. **(A)** Urea ChIP-seq (Urea) and std ChIP-seq (Std) profiles for SATB1 and input chromatin in a randomly chosen, 3.12 Mb region in chromosome 17 shows appearance of some peaks in input chromatin from std ChIP-seq that match with peaks for SATB1 in std ChIP-seq (top). For *Satb1*^−/−^ thymocytes (KO), using either anti-SATB1 antibody 1583 (rabbit polyclonal, SATB1-1) or rabbit monoclonal ab109122 (SATB1-2), urea and std ChIP-seq experiments for SATB1 were performed. With std ChIP-seq, multiple peaks were generated in KO thymocytes in the similar regions observed in WT thymocytes (panel A track 7 versus 9 and 10), while urea ChIP-seq produced only few peaks (panel A top, track 2 versus 4 and 5). A zoomed-in view of the blue-highlighted SATB1-clustered regions (bottom) shows that in standard ChIP-seq, multiple peaks in KO thymocytes co-mapped with peaks in WT thymocytes, while others did not (panel A bottom, track 7 versus 9 and 10). **(B)** Another randomly chosen 3.45 Mb region in X chromosome (panel B top) and its zoom-in view (panel B bottom) shows high incidence of std ChIP-seq peaks in KO thymocytes co-mapping with std ChIP-seq peaks in WT thymocytes (tracks 7 versus 9 and 10).

**Figure 5—figure supplement 1.**
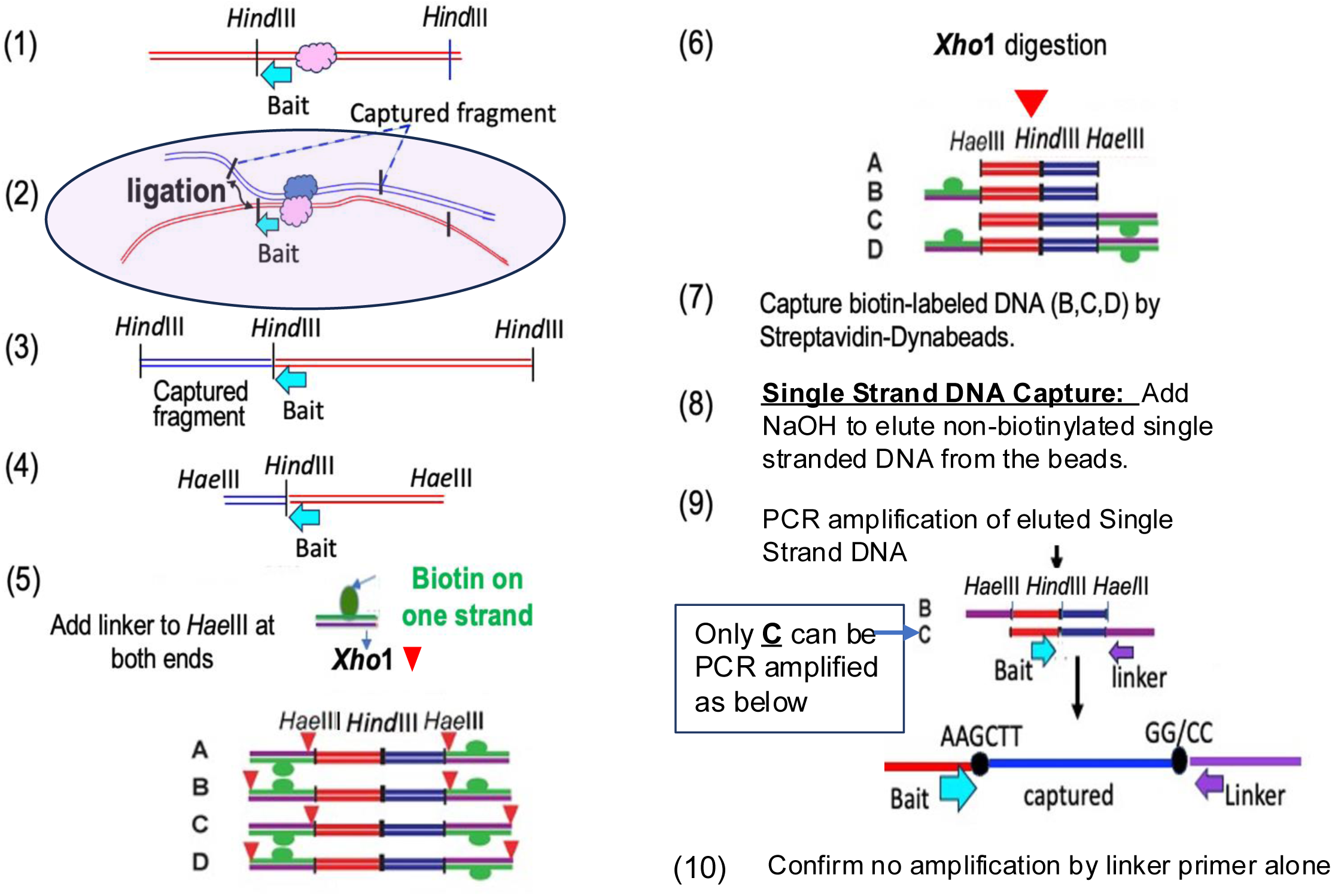
Urea 4C-seq protocol. Starting with urea-ultracentrifugation purified formaldehyde crosslinked chromatin: (**1**) A bait sequence is designed adjacent to a *Hind*III site of a *Hind*III–*Hind*III fragment containing SATB1 bound site (e.g. BUR-1). (**2**) Ligation is performed to ligate the original SATB1-bound fragment (bait) with other fragments whose close proximity was fixed by formaldehyde at the time of crosslinking and presumably retained after urea purification. The fixed chromatin structure with proximity is illustrated by pink shade. (**3**) Chromatin is decrosslinked and purified. (**4**) Ligated genomic DNA fragments are digested with *Hae*III (secondary restriction enzyme). (**5**) A Linker fragment containing an *Xho*1 site and Biotin labeled on one strand is blunt-end ligated to *Hae*III sites of the ligated genomic DNA fragments shown in step 4 to generate A, B, C, D double stranded fragments. The *Xho* 1 sites are indicated by red arrowhead. (**6**) The ligated products are digested with *Xho*1. (**7**) Biotin-labeled double stranded DNA (B, C, and D) are captured by Streptavidin-Dynabeads. (**8**) The beads are treated with NaOH to convert DNA to single strand DNA. This will result in dissociation and elution of non-biotinylated single strand DNA (derived from B and C) from the Dynabeads. (**9**) Using a specific primer including the *Hind*III of the bait sequence at the end and another primer from the Linker, captured DNA fragments are PCR amplified. (**10**) Control experiment is performed to confirm that Linker primer alone does not generate any amplified products. Once this is confirmed, the experiment is expected to contain minimum background signals from non-specific sequences.

**Figure 5—figure supplement 2.**
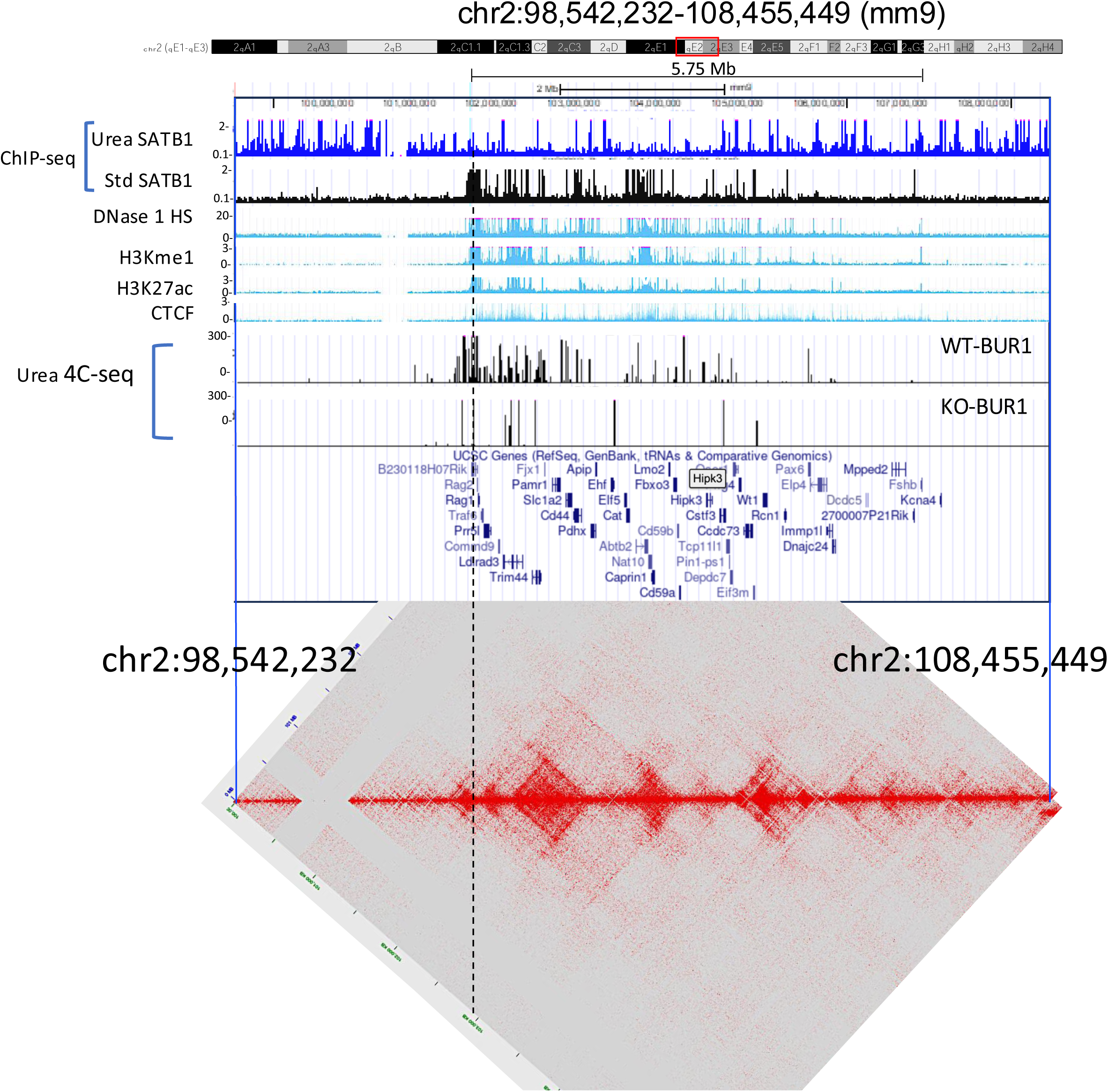
Long-range chromatin interactions from BUR-1 traverse TADs. BUR-1 interactions determined by urea4C-seq for wild type (WT) and SATB1-KO thymocytes covers the entire 5.75 Mb gene-rich region and these interactions bypass multiple small TADs revealed by the Hi-C heatmap (mm10) generated in the same region. Dotted line indicates the position of BUR-1

**Figure 5—figure supplement 3.**
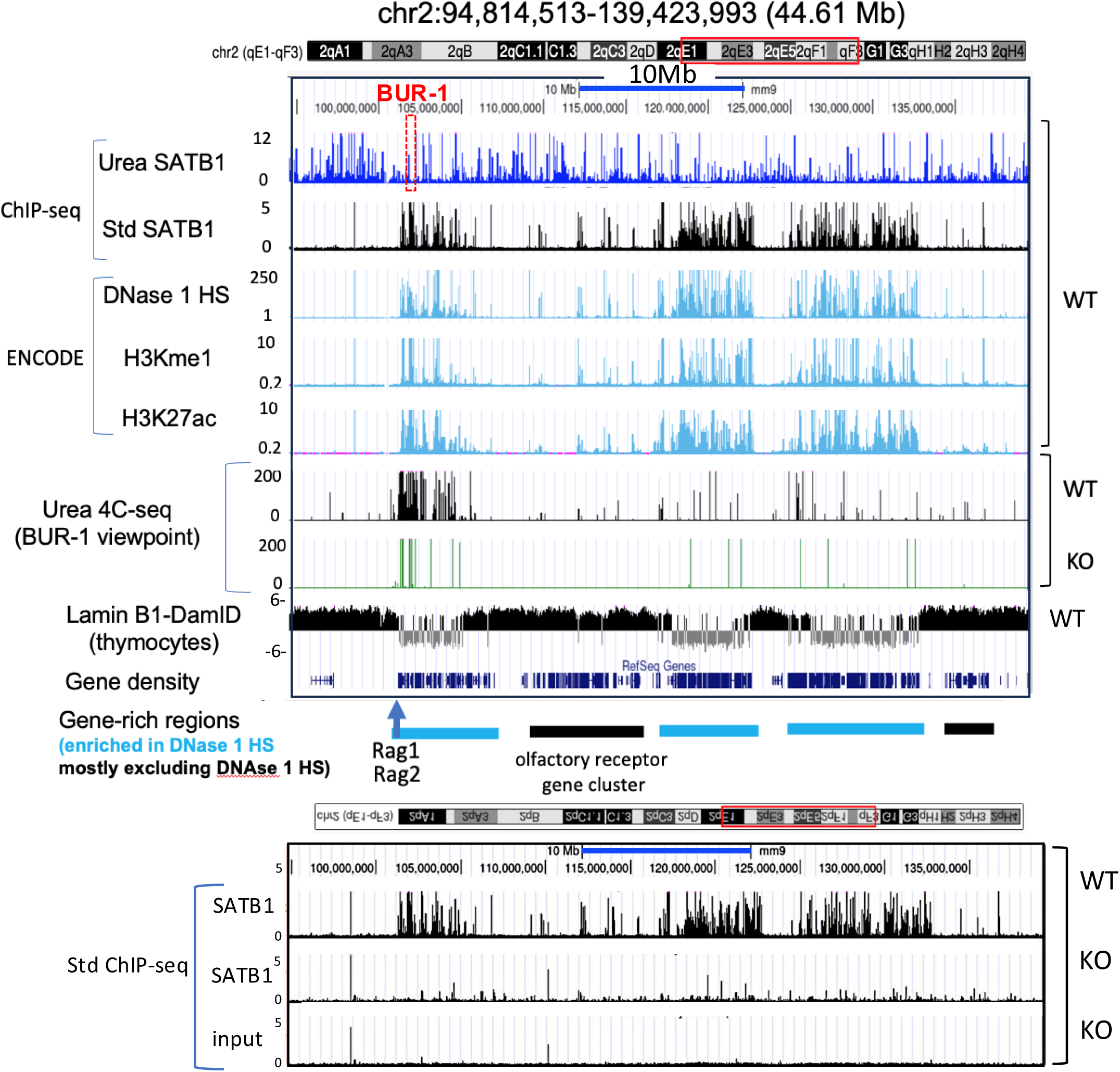
Long-distance interactions covering distal gene-rich regions by BUR-1. **(A)** A wide view (44.61 Mb) covering distal gene-rich regions shows BUR-1 (red bracket) frequently interacts with distal gene-rich DNase 1 HS regions including clusters of H3Kme1 and H3K27a (indicated by blue horizontal bars). DNase 1 hypersensitive (HS) regions, which indicate highly accessible chromatin (or “**open**” chromatin). These interactions were greatly reduced in SATB1 deficient thymocytes (KO). A gene-rich region (indicated by a black horizontal bar) that does not correspond to DNase 1 HS is enriched in olfactory receptor genes. Lamin B-DamID data shown is from Chen et al., *Cell Rep 2018* (Ref 86). H3Kme1 and H3K27ac data are from ENCODE (LICR) and DNase 1 HS data are from ENCODE (University of Washington). **(B)** SATB1 peaks identified by standard (Std) ChIP-seq in this region are mostly valid and only detected in wildtype (WT) and not in SATB1-null (KO) thymocytes, indicating that SATB1 is binding to these regions, presumably indirectly.

**Figure 5—figure supplement 4.**
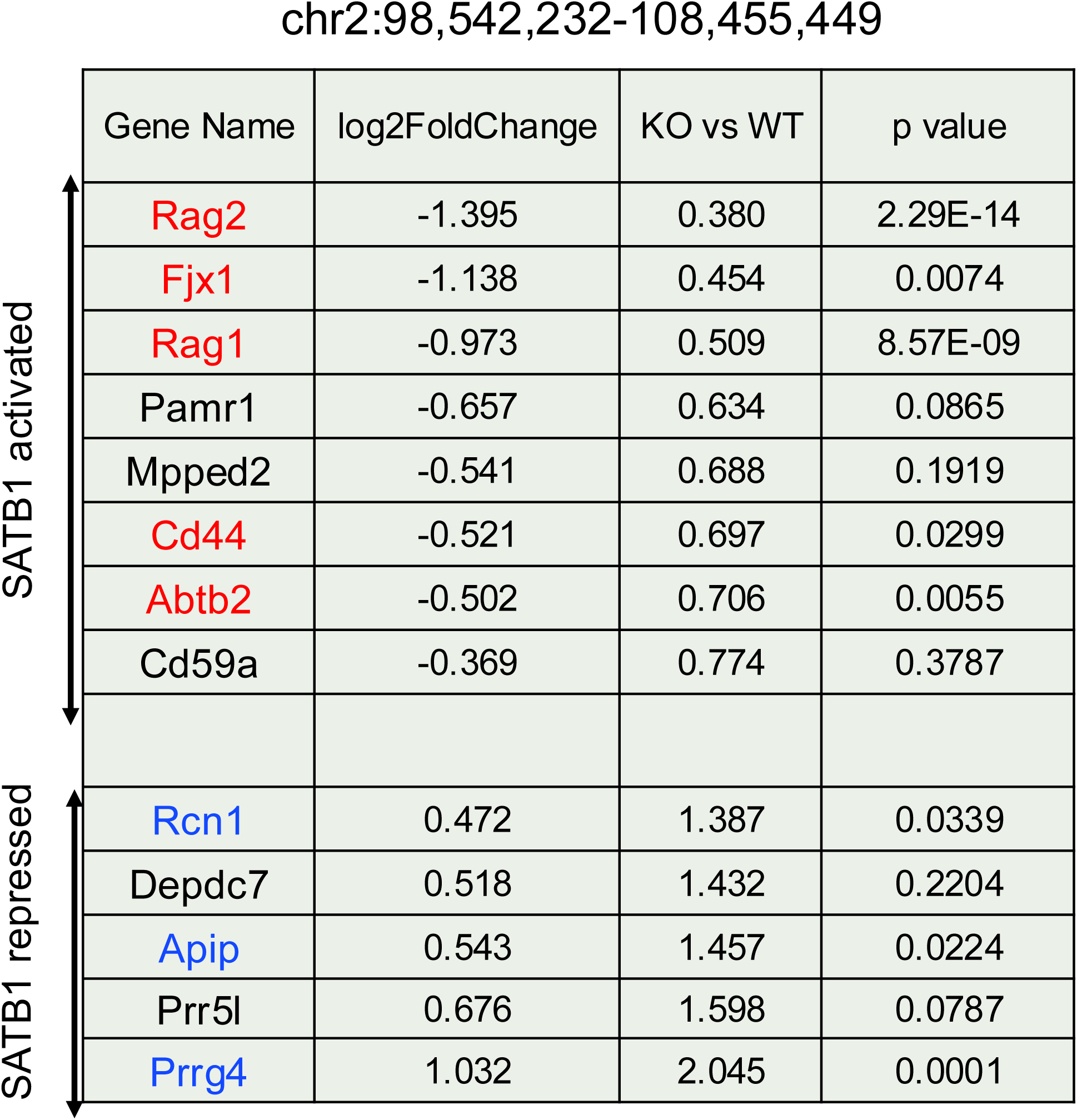
Genes within the gene-rich region (chr2:98,542,232-108,455,449) that exhibit SATB1-dependent expression. Genes showing differential expression for SATB1 with p<0.05 are shown in red (SATB1 activated) and in blue (SATB1 repressed). Data are from Zelenka et al., *Nat Commun*.,13, 6954, 2022 (Ref 54).

**Figure 7—figure supplement 1.**
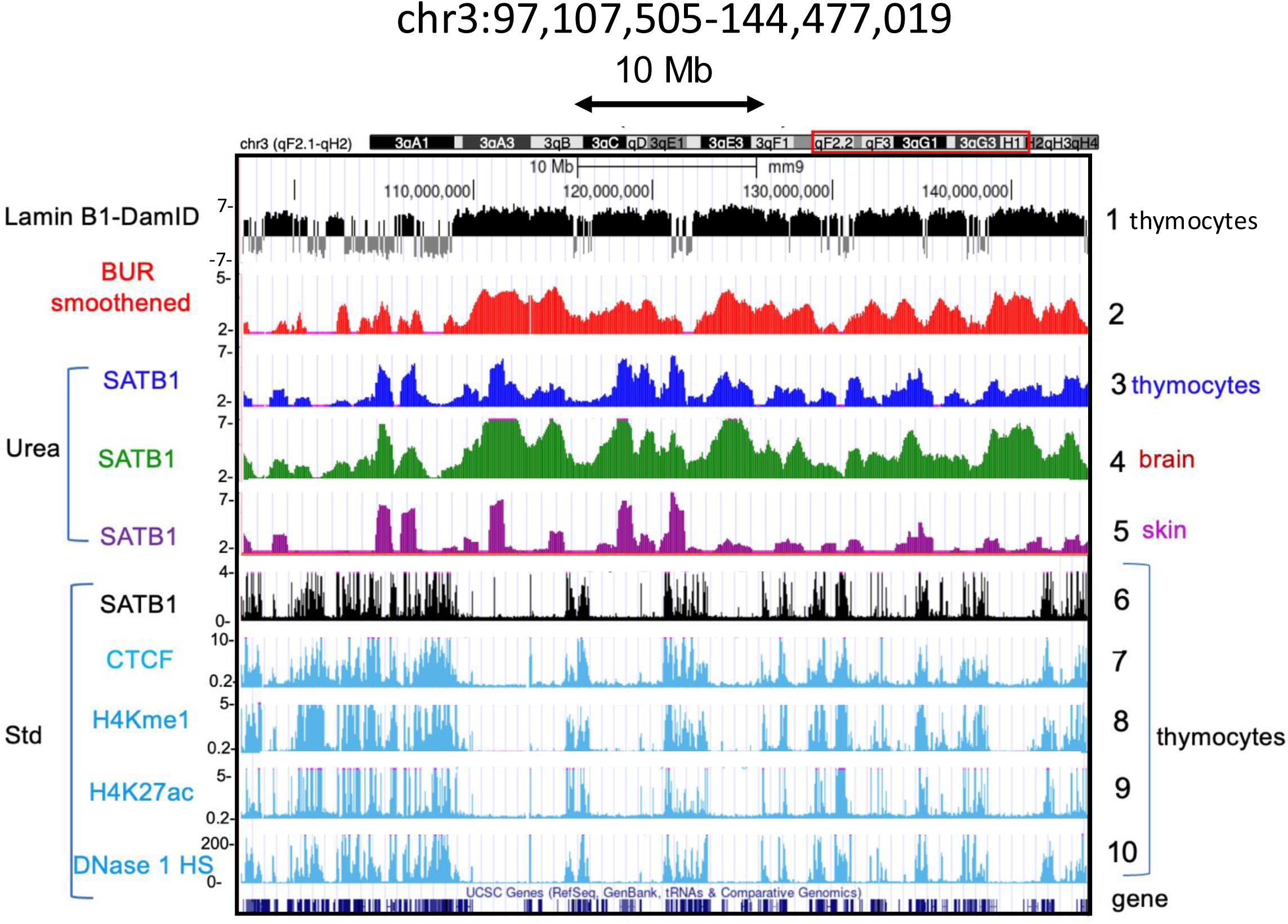
BURs and BUR-bound SATB1 largely co-map with LaminB1-DamID. The BUR distribution assessed by the BUR reference map (tracks 2) was compared against Lamin B1-DamID (track 1) over a 47.4Mbp region in chromosome 3. Lamin B1-DamID data are from Chen et al., *Cell Rep 2018* (Ref 86). The BUR distribution shows very similar overall distribution of LADs mapped by Lamin B1-DamID. BUR-binding frequency of SATB1 in thymocytes (track 3) is lower than that in brain (track 4). Distribution of BURs (track 2) and SATB1-binding regions determined by urea ChIP-seq (Urea) (track 3, 4, and 5) largely overlap with LADs (track 1). These regions mostly avoid DNase hypersensitive regions (HS) (track 10), which overlap with SATB1-binding sites determined by standard ChIP-seq (Std) (track 6), CTCF-binding sites (track 7), enhancer marks (H3Kme1 and H3K27ac (tracks 8 and 9, respectively). Data for CTCF, H3Kme1 and H3K27ac are from ENCODE (LICR) and for DNase 1 hypersensitive sites (HS) is from ENCODE (University of Washington).

**Figure 7—figure supplement 2.**
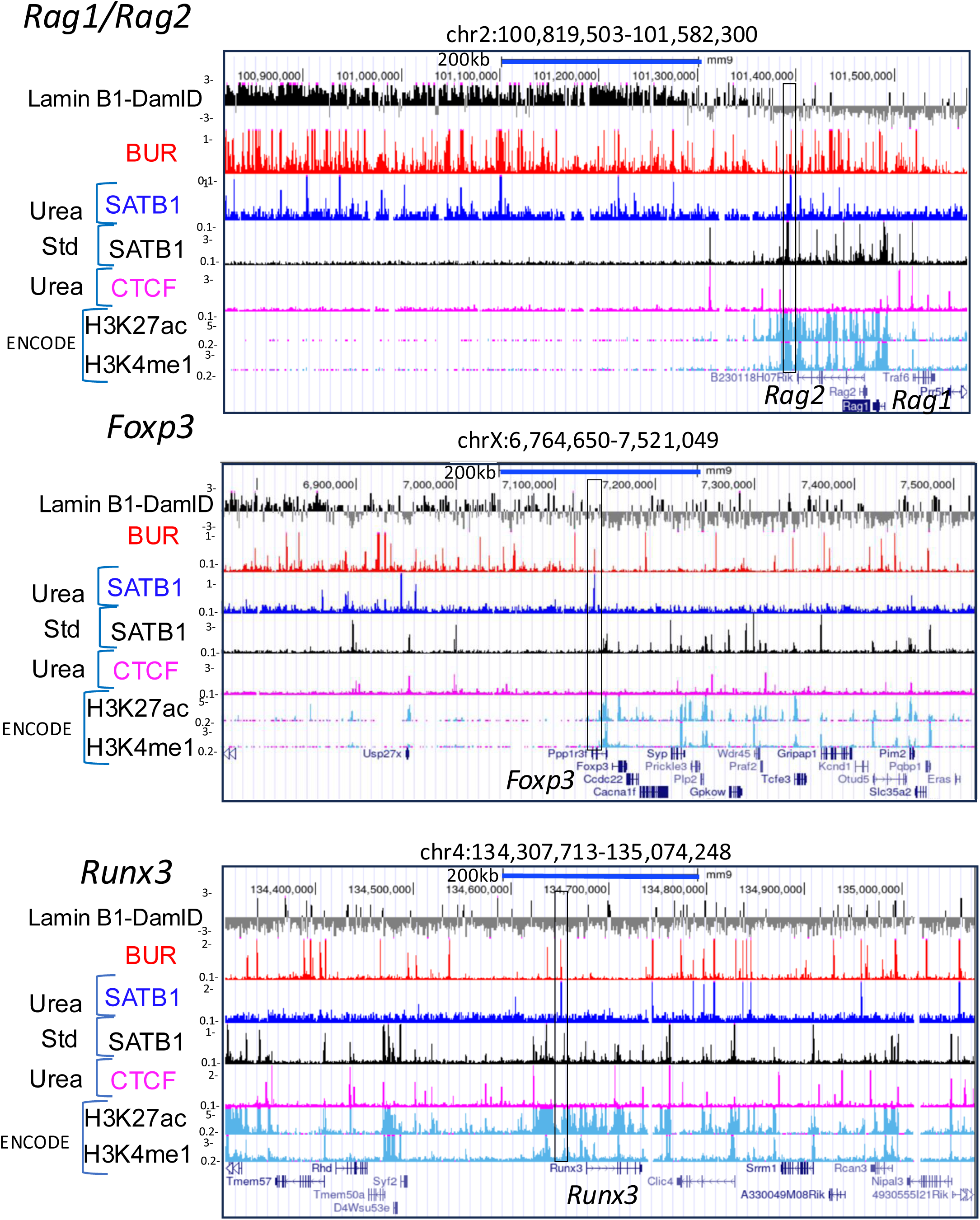
Some SATB1-bound BURs located in SATB1-dependent gene loci are outside of LADs. SATB1-bound BURs shown in Figure 4 (indicated by black box) in the *Rag1/Rag2, Foxp3, and Runx3* gene loci are found outside LADs mapped by LaminB1-DamID. These are close to H3Kme1 and H3K27ac sites. H3Kme1 and H3K27ac data are from ENCODE (LICR). Lamin B-DamID data shown is from Chen et al., *Cell Rep 2018* (Ref 86).

**Figure 7—figure supplement 3.**
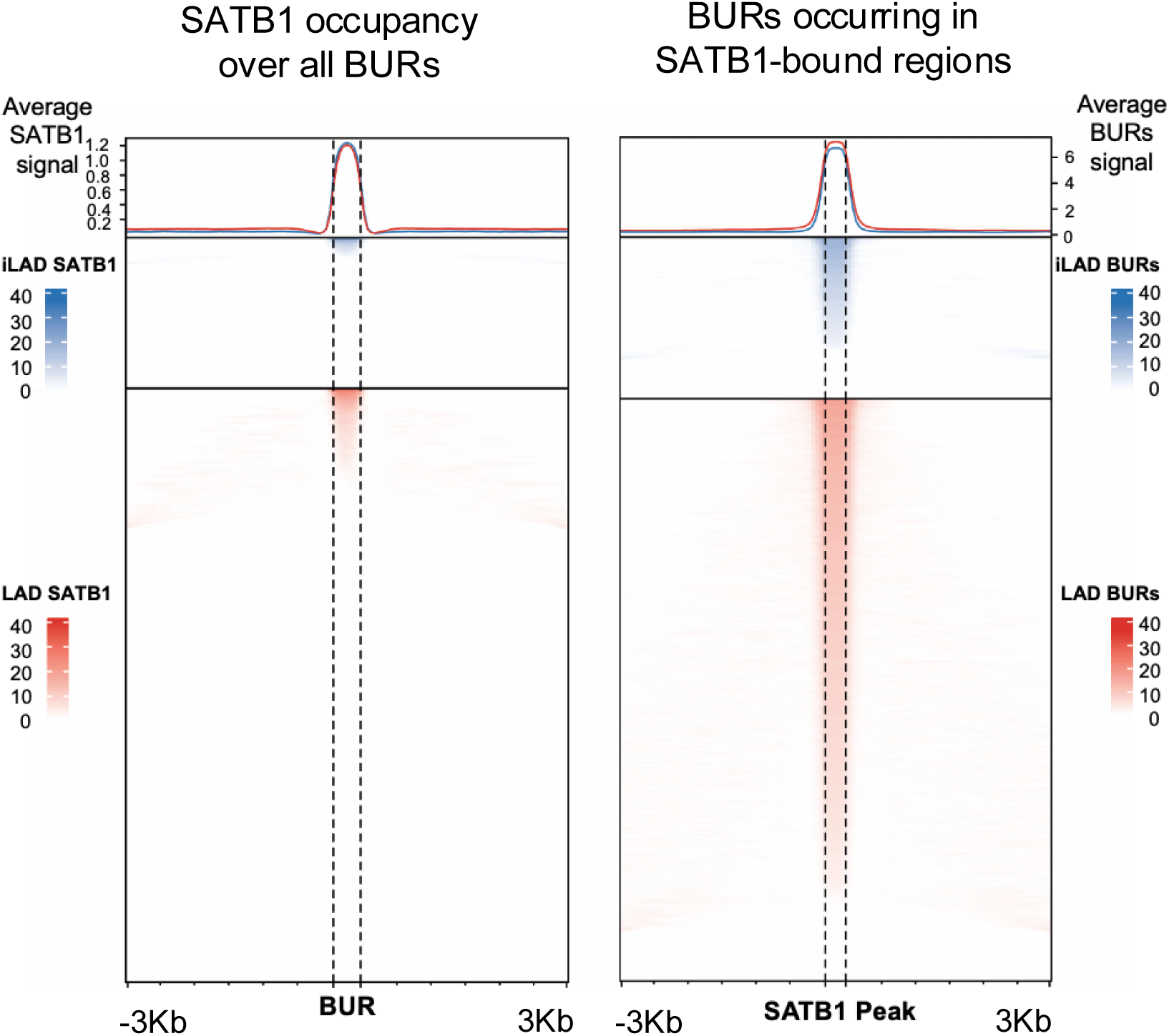
SATB1 occupancy relative to BURs, and BUR signal relative to SATB1-bound loci. (Left) Heatmaps showing SATB1 ChIP-seq signal intensity centered on all BUR elements, stratified by lamina association (LAD vs inter-LAD). Signal is plotted within ±3 kb of each BUR center using 50 bp bins using EnrichedHeatmap package (v1.30.0) The line plots above the heatmaps display the average SATB1 signal across all regions. Only a subset of BURs exhibit strong SATB1 binding, consistent with selective occupancy. (Right) Heatmaps showing BUR signal intensity centered on SATB1-bound regions (from ChIP-seq peaks), also separated into LAD and iLAD groups. The vast majority of SATB1 binding events are positioned within BURs, as evidenced by sharp, centered BUR signal at nearly all SATB1 peaks. Line plots reflect the average BUR signal profile across all SATB1 sites in each domain class. BUR intensities are derived from *in vitro* SATB1 BUR-binding assays.

**Figure 7—figure supplement 4.**
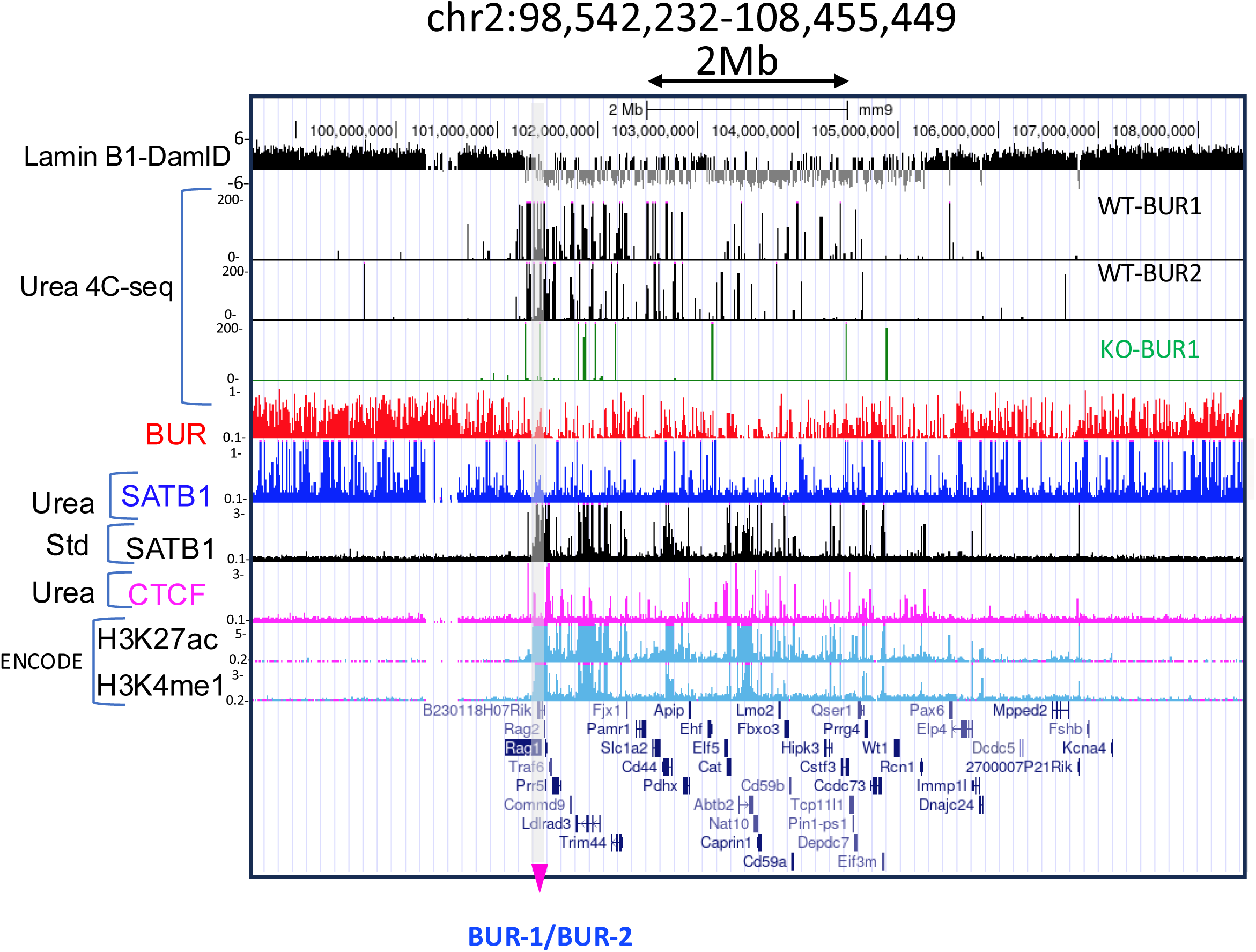
Interaction sites from BUR-1 or BUR-2 are mostly confined to the inter-LAD region. The region containing BUR-1 and BUR-2 as baits is light gray shaded with a pink arrowhead. This region is near the border of a LAD and an inter-LAD region, shown by LADs identified by Lamin B-DamID. Interaction profiles from either BUR-1 and BUR-2 obtained in wild-type thymocytes (WT) are found in a region surrounded by LADs. These interactions are greatly reduced in Satb1-KO thymocytes (KO). Many interaction sites are close to H3Kme1 and H3K27ac sites. H3Kme1 and H3K27ac data are from ENCODE (LICR). Lamin B-DamID data shown is from Chen et al., *Cell Rep 2018* (Ref 86).

**Supplementary Table 1.** Intersection of peaks were determined by deepTools between two samples of choice. Percent of peaks in each sample on the left column intersected with peaks in each of the samples on the top row are indicated.

|  | BUR-L | UREA SATB1 thymus | UREA SATB1 thymus | Standard SATB1 thymus | Standard SATB1 thymus | UREA SATB1 brain | UREA SATB1 brain | UREA SATB1 skin | UREA SATB1 skin | UREA CTCF thymus | UREA CTCF skin | Standard CTCF thymus |
| --- | --- | --- | --- | --- | --- | --- | --- | --- | --- | --- | --- | --- |
| % | A | B | C | D | E | F | G | H | I | J | K | L |
| A-120A |  | 12.68 | 14.34 | 0.77 | 0.60 | 36.41 | 33.54 | 3.69 | 3.62 | 0.25 | 0.58 | 0.60 |
| B- 355B | 77.71 |  | 85.02 | 1.12 | 0.86 | 92.07 | 90.28 | 24.13 | 23.56 | 0.77 | 1.00 | 0.86 |
| C-355C | 75.87 | 73.38 |  | 0.96 | 0.75 | 92.39 | 91.74 | 20.82 | 20.17 | 0.68 | 0.93 | 0.77 |
| D-328B | 3.83 | 0.91 | 0.90 |  | 47.92 | 1.28 | 1.11 | 1.93 | 1.91 | 22.35 | 23.31 | 36.39 |
| E-408B | 5.23 | 1.23 | 1.25 | 85.04 |  | 1.72 | 1.57 | 3.96 | 3.59 | 18.08 | 21.99 | 37.71 |
| F-345A | 73.58 | 30.35 | 35.28 | 0.52 | 0.40 |  | 78.12 | 7.80 | 7.59 | 0.31 | 0.58 | 0.56 |
| G-345B | 73.25 | 32.16 | 37.86 | 0.49 | 0.39 | 84.42 |  | 8.33 | 8.09 | 0.30 | 0.56 | 0.56 |
| H-404B | 78.54 | 83.69 | 83.65 | 8.30 | 9.59 | 82.06 | 81.13 |  | 81.06 | 2.35 | 12.85 | 1.78 |
| I-404E | 79.49 | 84.45 | 83.79 | 8.47 | 8.98 | 82.60 | 81.45 | 83.78 |  | 2.38 | 12.17 | 1.70 |
| J-346D | 2.96 | 1.47 | 1.52 | 52.97 | 24.15 | 1.78 | 1.61 | 1.29 | 1.27 |  | 83.53 | 60.22 |
| K-403C | 4.36 | 1.23 | 1.34 | 35.64 | 18.94 | 2.18 | 1.95 | 4.57 | 4.19 | 53.87 |  | 42.86 |
| L-Encode LICR | 2.68 | 0.63 | 0.65 | 33.00 | 19.27 | 1.24 | 1.14 | 0.37 | 0.35 | 23.04 | 25.42 |  |

**Supplementary Table 2.** Identification of each sample and their total number of peaks are shown.

|  |  | Sample ID | Total #Peaks |
| --- | --- | --- | --- |
| A. BUR-L |  | A-120A | 238380 |
| B. UREA SATB1 thymus | replica 1 | B-355B | 38882 |
| C. UREA SATB1 thymus | replica 2 | C-355C | 45049 |
| D. Std SATB1 thymus | replica 1 | D-328B | 48175 |
| E. Std SATB1 thymus | replica 2 | E-408B | 27147 |
| F. UREA SATB1 brain | replica 1 | F-345A | 117963 |
| G. UREA SATB1 brain | replica 2 | G-345B | 109154 |
| H. UREA SATB1 skin | replica 1 | H-404B | 11210 |
| I. UREA SATB1 skin | replica 2 | I-404E | 10846 |
| J. UREA CTCF thymus | replica 1 | J-346D | 20323 |
| K. UREA CTCF skin | replica 1 | K-403C | 31509 |
| L. Std CTCF thymus | Encode | L-Encode LICR | 53124 |

**Supplementary Table 3:** Intersection of peaks were determined by deepTools between two samples of choice. Percent of peaks in each sample on the left column that intersected with peaks in each sample on the top row are indicated. Important sets of data for comparison are highlighted in blue for urea ChIP-seq and pink for standard ChIP-seq.

|  | urea ChIP-seq |  |  |  |  | standard ChIP-seq |  |  |  |  |
| --- | --- | --- | --- | --- | --- | --- | --- | --- | --- | --- |
|  | TKS355B | TKS400B | TKS400D | TKS340A | TKS356A | TKS408B | TKS407D | TKS407B | TKS390A | TKS407C |
|  | SATB1 ChIP | Phantom (KO SATB1 ChIP) | Phantom (KO SATB1 ChIP) | Urea Input | Urea Input | SATB1 ChIP | Phantom (KO SATB1 ChIP) | Phantom (KO SATB1 ChIP) | Standard Input | Standard Input |
|  | UREA | UREA antibody 1 | UREA antibody 2 | WT | KO | STANDARD | Standard Antibody 1 | Standard Antibody 2 | WT | KO |
| TKS355B |  | 0.0 | 0.0 | 0.4 | 0.4 | 0.8 | 0.1 | 0.1 | 0.5 | 0.3 |
| TKS400B | 1.0 |  | 78.5 | 33.6 | 6.1 | 85.7 | 89.7 | 93.8 | 53.0 | 15.3 |
| TKS400D | 0.8 | 66.1 |  | 25.4 | 4.2 | 88.9 | 93.2 | 93.2 | 41.5 | 11.8 |
| TKS340A | 10.7 | 1.4 | 1.3 |  | 51.2 | 55.6 | 7.8 | 10.6 | 79.2 | 39.3 |
| TKS356A | 17.7 | 0.4 | 0.3 | 90.8 |  | 76.1 | 8.3 | 12.3 | 93.4 | 59.4 |
| TKS408B | 1.2 | 0.3 | 0.3 | 4.1 | 3.1 |  | 3.6 | 4.5 | 12.7 | 3.0 |
| TKS407D | 2.3 | 3.1 | 3.8 | 6.7 | 4.3 | 36.4 |  | 90.9 | 9.6 | 4.1 |
| TKS407B | 1.3 | 1.3 | 1.5 | 3.5 | 2.3 | 18.2 | 37.2 |  | 4.9 | 2.2 |
| TKS390A | 3.6 | 0.7 | 0.6 | 25.2 | 16.9 | 52.0 | 3.6 | 4.6 |  | 14.3 |
| TKS407C | 16.7 | 1.2 | 1.1 | 75.7 | 63.9 | 79.8 | 8.3 | 12.0 | 86.5 |  |

**Supplementary Table 4:** Identification of each sample and their total number of peaks are shown.

| Sample ID | Description | Total # Peaks |
| --- | --- | --- |
| TKS355B | SATB1 ureaChIP-seq | 38882 |
| TKS400B | KO SATB1 ureaChIP-seq 1 | 98 |
| TKS400D | KO SATB1 ureaChIP-seq 2 | 118 |
| TKS340A | WT Input ureaChIP-seq 1 | 2381 |
| TKS356A | KO Input ureaChIP-seq 1 | 1372 |
| TKS408B | SATB1 standard ChIP-seq | 27147 |
| TKS407D | KO SATB1 standard ChIP-seq 1 | 2884 |
| TKS407B | KO SATB1 standard ChIP-seq 2 | 7003 |
| TKS390A | WT Input Standard ChIP-seq 1 | 7377 |
| TKS407C | KO Input Standard ChIP-seq 1 | 1224 |

## References

1. J. C. Politz, D. Scalzo, M. Groudine, Something silent this way forms: the functional organization of the repressive nuclear compartment. Annu Rev Cell Dev Biol 29, 241–270 (2013). DOI: 10.1146/annurev-cellbio-101512-122317.

2. B. van Steensel, A. S. Belmont, Lamina-Associated Domains: Links with Chromosome Architecture, Heterochromatin, and Gene Repression. Cell 169, 780–791 (2017). DOI: 10.1016/j.cell.2017.04.022.

3. S. Schoenfelder, P. Fraser, Long-range enhancer-promoter contacts in gene expression control. Nat Rev Genet 20, 437–455 (2019). DOI: 10.1038/s41576-019-0128-0.

4. E. Bashkirova, S. Lomvardas, Olfactory receptor genes make the case for inter-chromosomal interactions. Curr Opin Genet Dev 55, 106–113 (2019). DOI: 10.1016/j.gde.2019.07.004.

5. A. Gondor, R. Ohlsson, Enhancer functions in three dimensions: beyond the flat world perspective. F1000Res 7, (2018). DOI: 10.12688/f1000research.13842.1.

6. V. Loubiere, G. L. Papadopoulos, Q. Szabo, A. M. Martinez, G. Cavalli, Widespread activation of developmental gene expression characterized by PRC1-dependent chromatin looping. Sci Adv 6, eaax4001 (2020). DOI: 10.1126/sciadv.aax4001.

7. T. Misteli, The Self-Organizing Genome: Principles of Genome Architecture and Function. Cell 183, 28–45 (2020). DOI: 10.1016/j.cell.2020.09.014.

8. E. Lieberman-Aiden, N. L. van Berkum, L. Williams, M. Imakaev, T. Ragoczy, A. Telling, I. Amit, B. R. Lajoie, P. J. Sabo, M. O. Dorschner, R. Sandstrom, B. Bernstein, M. A. Bender, M. Groudine, A. Gnirke, J. Stamatoyannopoulos, L. A. Mirny, E. S. Lander, J. Dekker, Comprehensive mapping of long-range interactions reveals folding principles of the human genome. Science 326, 289–293 (2009). DOI: 10.1126/science.1181369.

9. L. Guelen, L. Pagie, E. Brasset, W. Meuleman, M. B. Faza, W. Talhout, B. H. Eussen, A. de Klein, L. Wessels, W. de Laat, B. van Steensel, Domain organization of human chromosomes revealed by mapping of nuclear lamina interactions. Nature 453, 948–951 (2008). DOI: 10.1038/nature06947.

10. J. Kind, L. Pagie, S. S. de Vries, L. Nahidiazar, S. S. Dey, M. Bienko, Y. Zhan, B. Lajoie, C. A. de Graaf, M. Amendola, G. Fudenberg, M. Imakaev, L. A. Mirny, K. Jalink, J. Dekker, A. van Oudenaarden, B. van Steensel, Genome-wide maps of nuclear lamina interactions in single human cells. Cell 163, 134–147 (2015). DOI: 10.1016/j.cell.2015.08.040.

11. B. van Steensel, J. Delrow, S. Henikoff, Chromatin profiling using targeted DNA adenine methyltransferase. Nat Genet 27, 304–308 (2001). DOI: 10.1038/85871.

12. J. R. Dixon, S. Selvaraj, F. Yue, A. Kim, Y. Li, Y. Shen, M. Hu, J. S. Liu, B. Ren, Topological domains in mammalian genomes identified by analysis of chromatin interactions. Nature 485, 376–380 (2012). DOI: 10.1038/nature11082.

13. E. P. Nora, B. R. Lajoie, E. G. Schulz, L. Giorgetti, I. Okamoto, N. Servant, T. Piolot, N. L. van Berkum, J. Meisig, J. Sedat, J. Gribnau, E. Barillot, N. Bluthgen, J. Dekker, E. Heard, Spatial partitioning of the regulatory landscape of the X-inactivation centre. Nature 485, 381–385 (2012). DOI: 10.1038/nature11049.

14. T. Sexton, E. Yaffe, E. Kenigsberg, F. Bantignies, B. Leblanc, M. Hoichman, H. Parrinello, A. Tanay, G. Cavalli, Three-dimensional folding and functional organization principles of the Drosophila genome. Cell 148, 458–472 (2012). DOI: 10.1016/j.cell.2012.01.010.

15. A. L. Sanborn, S. S. Rao, S. C. Huang, N. C. Durand, M. H. Huntley, A. I. Jewett, I. D. Bochkov, D. Chinnappan, A. Cutkosky, J. Li, K. P. Geeting, A. Gnirke, A. Melnikov, D. McKenna, E. K. Stamenova, E. S. Lander, E. L. Aiden, Chromatin extrusion explains key features of loop and domain formation in wild-type and engineered genomes. Proc Natl Acad Sci U S A 112, E6456–6465 (2015). DOI: 10.1073/pnas.1518552112.

16. G. Fudenberg, M. Imakaev, C. Lu, A. Goloborodko, N. Abdennur, L. A. Mirny, Formation of Chromosomal Domains by Loop Extrusion. Cell Rep 15, 2038–2049 (2016). DOI: 10.1016/j.celrep.2016.04.085.

17. S. S. Rao, M. H. Huntley, N. C. Durand, E. K. Stamenova, I. D. Bochkov, J. T. Robinson, A. L. Sanborn, I. Machol, A. D. Omer, E. S. Lander, E. L. Aiden, A 3D Map of the Human Genome at Kilobase Resolution Reveals Principles of Chromatin Looping. Cell 159, 1665–1680 (2014). DOI: 10.1016/j.cell.2014.11.021.

18. E. P. Nora, A. Goloborodko, A. L. Valton, J. H. Gibcus, A. Uebersohn, N. Abdennur, J. Dekker, L. A. Mirny, B. G. Bruneau, Targeted Degradation of CTCF Decouples Local Insulation of Chromosome Domains from Genomic Compartmentalization. Cell 169, 930–944 e922 (2017). DOI: 10.1016/j.cell.2017.05.004.

19. S. S. P. Rao, S. C. Huang, B. Glenn St Hilaire, J. M. Engreitz, E. M. Perez, K. R. Kieffer-Kwon, A. L. Sanborn, S. E. Johnstone, G. D. Bascom, I. D. Bochkov, X. Huang, M. S. Shamim, J. Shin, D. Turner, Z. Ye, A. D. Omer, J. T. Robinson, T. Schlick, B. E. Bernstein, R. Casellas, E. S. Lander, E. L. Aiden, Cohesin Loss Eliminates All Loop Domains. Cell 171, 305–320 e324 (2017). DOI: 10.1016/j.cell.2017.09.026.

20. G. Wutz, C. Varnai, K. Nagasaka, D. A. Cisneros, R. R. Stocsits, W. Tang, S. Schoenfelder, G. Jessberger, M. Muhar, M. J. Hossain, N. Walther, B. Koch, M. Kueblbeck, J. Ellenberg, J. Zuber, P. Fraser, J. M. Peters, Topologically associating domains and chromatin loops depend on cohesin and are regulated by CTCF, WAPL, and PDS5 proteins. EMBO J 36, 3573–3599 (2017). DOI: 10.15252/embj.201798004.

21. W. Schwarzer, N. Abdennur, A. Goloborodko, A. Pekowska, G. Fudenberg, Y. Loe-Mie, N. A. Fonseca, W. Huber, C. H. Haering, L. Mirny, F. Spitz, Two independent modes of chromatin organization revealed by cohesin removal. Nature 551, 51–56 (2017). DOI: 10.1038/nature24281.

22. D. I. Cattoni, A. M. Cardozo Gizzi, M. Georgieva, M. Di Stefano, A. Valeri, D. Chamousset, C. Houbron, S. Dejardin, J. B. Fiche, I. Gonzalez, J. M. Chang, T. Sexton, M. A. Marti-Renom, F. Bantignies, G. Cavalli, M. Nollmann, Single-cell absolute contact probability detection reveals chromosomes are organized by multiple low-frequency yet specific interactions. Nat Commun 8, 1753 (2017). DOI: 10.1038/s41467-017-01962-x.

23. B. Bintu, L. J. Mateo, J. H. Su, N. A. Sinnott-Armstrong, M. Parker, S. Kinrot, K. Yamaya, A. N. Boettiger, X. Zhuang, Super-resolution chromatin tracing reveals domains and cooperative interactions in single cells. Science 362, (2018). DOI: 10.1126/science.aau1783.

24. E. H. Finn, G. Pegoraro, H. B. Brandao, A. L. Valton, M. E. Oomen, J. Dekker, L. Mirny, T. Misteli, Extensive Heterogeneity and Intrinsic Variation in Spatial Genome Organization. Cell 176, 1502–1515 e1510 (2019). DOI: 10.1016/j.cell.2019.01.020.

25. M. Gabriele, H. B. Brandao, S. Grosse-Holz, A. Jha, G. M. Dailey, C. Cattoglio, T. S. Hsieh, L. Mirny, C. Zechner, A. S. Hansen, Dynamics of CTCF- and cohesin-mediated chromatin looping revealed by live-cell imaging. Science 376, 496–501 (2022). DOI: 10.1126/science.abn6583.

26. P. Mach, P. I. Kos, Y. Zhan, J. Cramard, S. Gaudin, J. Tunnermann, E. Marchi, J. Eglinger, J. Zuin, M. Kryzhanovska, S. Smallwood, L. Gelman, G. Roth, E. P. Nora, G. Tiana, L. Giorgetti, Cohesin and CTCF control the dynamics of chromosome folding. Nat Genet 54, 1907–1918 (2022). DOI: 10.1038/s41588-022-01232-7.

27. Y. Chi, J. Shi, D. Xing, L. Tan, Every gene everywhere all at once: High-precision measurement of 3D chromosome architecture with single-cell Hi-C. Front Mol Biosci 9, 959688 (2022). DOI: 10.3389/fmolb.2022.959688.

28. L. F. Chen, J. Lee, A. Boettiger, Recent progress and challenges in single-cell imaging of enhancer-promoter interaction. Curr Opin Genet Dev 79, 102023 (2023). DOI: 10.1016/j.gde.2023.102023.

29. A. Hafner, M. Park, S. E. Berger, S. E. Murphy, E. P. Nora, A. N. Boettiger, Loop stacking organizes genome folding from TADs to chromosomes. Mol Cell 83, 1377–1392 e1376 (2023). DOI: 10.1016/j.molcel.2023.04.008.

30. T. Kohwi-Shigematsu, Y. Kohwi, Torsional stress stabilizes extended base unpairing in suppressor sites flanking immunoglobulin heavy chain enhancer. Biochemistry 29, 9551–9560. (1990). DOI:

31. J. Bode, Y. Kohwi, L. Dickinson, T. Joh, D. Klehr, C. Mielke, T. Kohwi-Shigematsu, Biological significance of unwinding capability of nuclear matrix-associating DNAs. Science 255, 195–197 (1992). DOI: 10.1126/science.1553545.

32. L. A. Dickinson, T. Joh, Y. Kohwi, T. Kohwi-Shigematsu, A tissue-specific MAR/SAR DNA-binding protein with unusual binding site recognition. Cell 70, 631–645. (1992). DOI:

33. T. Kohwi-Shigematsu, Y. Kohwi, Detection of non-B-DNA structures at specific sites in supercoiled plasmid DNA and chromatin with haloacetaldehyde and diethyl pyrocarbonate. Methods Enzymol 212, 155–180 (1992). DOI:

34. J. D. Alvarez, D. H. Yasui, H. Niida, T. Joh, D. Y. Loh, T. Kohwi-Shigematsu, The MAR-binding protein SATB1 orchestrates temporal and spatial expression of multiple genes during T-cell development. Genes Dev 14, 521–535 (2000). DOI:

35. Y. Zhang, L. Zheng, M. Le, Y. Nakano, B. Chan, Y. Huang, P. M. Torbaty, Y. Kohwi, R. Marcucio, S. Habelitz, P. K. Den Besten, T. Kohwi-Shigematsu, SATB1 establishes ameloblast cell polarity and regulates directional amelogenin secretion for enamel formation. BMC Biol 17, 104 (2019). DOI: 10.1186/s12915-019-0722-9.

36. M. Y. Fessing, A. N. Mardaryev, M. R. Gdula, A. A. Sharov, T. Y. Sharova, V. Rapisarda, K. B. Gordon, A. D. Smorodchenko, K. Poterlowicz, G. Ferone, Y. Kohwi, C. Missero, T. Kohwi-Shigematsu, V. A. Botchkarev, p63 regulates Satb1 to control tissue-specific chromatin remodeling during development of the epidermis. J Cell Biol 194, 825–839 (2011). DOI: jcb.201101148 [pii] 10.1083/jcb.201101148.

37. M. A. Balamotis, N. Tamberg, Y. J. Woo, J. Li, B. Davy, T. Kohwi-Shigematsu, Y. Kohwi, Satb1 ablation alters temporal expression of immediate early genes and reduces dendritic spine density during postnatal brain development. Mol Cell Biol 32, 333–347 (2012). DOI: 10.1128/MCB.05917-11.

38. Y. Huang, L. Zhang, N. N. Song, Z. L. Hu, J. Y. Chen, Y. Q. Ding, Distribution of Satb1 in the central nervous system of adult mice. Neurosci Res 71, 12–21 (2011). DOI: 10.1016/j.neures.2011.05.015.

39. S. Cai, H. J. Han, T. Kohwi-Shigematsu, Tissue-specific nuclear architecture and gene expression regulated by SATB1. Nat Genet 34, 42–51 (2003). DOI: 10.1038/ng1146.

40. I. de Belle, S. Cai, T. Kohwi-Shigematsu, The genomic sequences bound to special AT-rich sequence-binding protein 1 (SATB1) in vivo in Jurkat T cells are tightly associated with the nuclear matrix at the bases of the chromatin loops. J Cell Biol 141, 335–348 (1998). DOI: 10.1083/jcb.141.2.335.

41. T. Kohwi-Shigematsu, K. Poterlowicz, E. Ordinario, H. J. Han, V. A. Botchkarev, Y. Kohwi, Genome organizing function of SATB1 in tumor progression. Semin Cancer Biol, (2013). DOI: S1044-579X(12)00101-0 [pii] 10.1016/j.semcancer.2012.06.009.

42. S. Cai, C. C. Lee, T. Kohwi-Shigematsu, SATB1 packages densely looped, transcriptionally active chromatin for coordinated expression of cytokine genes. Nat Genet 38, 1278–1288 (2006). DOI:

43. Y. Satoh, T. Yokota, T. Sudo, M. Kondo, A. Lai, P. W. Kincade, T. Kouro, R. Iida, K. Kokame, T. Miyata, Y. Habuchi, K. Matsui, H. Tanaka, I. Matsumura, K. Oritani, T. Kohwi-Shigematsu, Y. Kanakura, The Satb1 protein directs hematopoietic stem cell differentiation toward lymphoid lineages. Immunity 38, 1105–1115 (2013). DOI: 10.1016/j.immuni.2013.05.014.

44. D. Skowronska-Krawczyk, Q. Ma, M. Schwartz, K. Scully, W. Li, Z. Liu, H. Taylor, J. Tollkuhn, K. A. Ohgi, D. Notani, Y. Kohwi, T. Kohwi-Shigematsu, M. G. Rosenfeld, Required enhancer-matrin-3 network interactions for a homeodomain transcription program. Nature 514, 257–261 (2014). DOI: 10.1038/nature13573.

45. B. Hao, A. K. Naik, A. Watanabe, H. Tanaka, L. Chen, H. W. Richards, M. Kondo, I. Taniuchi, Y. Kohwi, T. Kohwi-Shigematsu, M. S. Krangel, An anti-silencer- and SATB1-dependent chromatin hub regulates Rag1 and Rag2 gene expression during thymocyte development. J Exp Med 212, 809–824 (2015). DOI: 10.1084/jem.20142207.

46. K. Kakugawa, S. Kojo, H. Tanaka, W. Seo, T. A. Endo, Y. Kitagawa, S. Muroi, M. Tenno, N. Yasmin, Y. Kohwi, S. Sakaguchi, T. Kowhi-Shigematsu, I. Taniuchi, Essential Roles of SATB1 in Specifying T Lymphocyte Subsets. Cell Rep 19, 1176–1188 (2017). DOI: 10.1016/j.celrep.2017.04.038.

47. Y. Kitagawa, N. Ohkura, Y. Kidani, A. Vandenbon, K. Hirota, R. Kawakami, K. Yasuda, D. Motooka, S. Nakamura, M. Kondo, I. Taniuchi, T. Kohwi-Shigematsu, S. Sakaguchi, Guidance of regulatory T cell development by Satb1-dependent super-enhancer establishment. Nat Immunol 18, 173–183 (2017). DOI: 10.1038/ni.3646.

48. K. Yasuda, Y. Kitagawa, R. Kawakami, Y. Isaka, H. Watanabe, G. Kondoh, T. Kohwi-Shigematsu, S. Sakaguchi, K. Hirota, Satb1 regulates the effector program of encephalitogenic tissue Th17 cells in chronic inflammation. Nat Commun 10, 549 (2019). DOI: 10.1038/s41467-019-08404-w.

49. M. Goolam, M. Zernicka-Goetz, The chromatin modifier Satb1 regulates cell fate through Fgf signalling in the early mouse embryo. Development 144, 1450–1461 (2017). DOI: 10.1242/dev.144139.

50. H. J. Han, J. Russo, Y. Kohwi, T. Kohwi-Shigematsu, SATB1 reprogrammes gene expression to promote breast tumour growth and metastasis. Nature 452, 187–193 (2008). DOI:

51. A. Fromberg, K. Engeland, A. Aigner, The Special AT-rich Sequence Binding Protein 1 (SATB1) and its role in solid tumors. Cancer Lett 417, 96–111 (2018). DOI: 10.1016/j.canlet.2017.12.031.

52. E. Ordinario, H. J. Han, S. Furuta, L. M. Heiser, L. R. Jakkula, F. Rodier, P. T. Spellman, J. Campisi, J. W. Gray, M. J. Bissell, Y. Kohwi, T. Kohwi-Shigematsu, ATM suppresses SATB1-induced malignant progression in breast epithelial cells. PLoS One 7, e51786 (2012). DOI: 10.1371/journal.pone.0051786.

53. D. Feng, Y. Chen, R. Dai, S. Bian, W. Xue, Y. Zhu, Z. Li, Y. Yang, Y. Zhang, J. Zhang, J. Bai, L. Qin, Y. Kohwi, W. Shi, T. Kohwi-Shigematsu, J. Ma, S. Liao, B. Hao, Chromatin organizer SATB1 controls the cell identity of CD4(+) CD8(+) double-positive thymocytes by regulating the activity of super-enhancers. Nat Commun 13, 5554 (2022). DOI: 10.1038/s41467-022-33333-6.

54. T. Zelenka, A. Klonizakis, D. Tsoukatou, D. A. Papamatheakis, S. Franzenburg, P. Tzerpos, I. R. Tzonevrakis, G. Papadogkonas, M. Kapsetaki, C. Nikolaou, D. Plewczynski, C. Spilianakis, The 3D enhancer network of the developing T cell genome is shaped by SATB1. Nat Commun 13, 6954 (2022). DOI: 10.1038/s41467-022-34345-y.

55. B. Wang, L. Ji, Q. Bian, SATB1 regulates 3D genome architecture in T cells by constraining chromatin interactions surrounding CTCF-binding sites. Cell Rep 42, 112323 (2023). DOI: 10.1016/j.celrep.2023.112323.

56. D. Jain, S. Baldi, A. Zabel, T. Straub, P. B. Becker, Active promoters give rise to false positive ‘Phantom Peaks’ in ChIP-seq experiments. Nucleic Acids Res 43, 6959–6968 (2015). DOI: 10.1093/nar/gkv637.

57. D. Park, Y. Lee, G. Bhupindersingh, V. R. Iyer, Widespread misinterpretable ChIP-seq bias in yeast. PLoS One 8, e83506 (2013). DOI: 10.1371/journal.pone.0083506.

58. L. Teytelman, D. M. Thurtle, J. Rine, A. van Oudenaarden, Highly expressed loci are vulnerable to misleading ChIP localization of multiple unrelated proteins. Proc Natl Acad Sci U S A 110, 18602–18607 (2013). DOI: 10.1073/pnas.1316064110.

59. C. A. Meyer, X. S. Liu, Identifying and mitigating bias in next-generation sequencing methods for chromatin biology. Nat Rev Genet 15, 709–721 (2014). DOI: 10.1038/nrg3788.

60. S. Baumgarten, J. Bryant, Chromatin structure can introduce systematic biases in genome-wide analyses of Plasmodium falciparum. Open Res Eur 2, 75 (2022). DOI: 10.12688/openreseurope.14836.2.

61. K. Gesson, P. Rescheneder, M. P. Skoruppa, A. von Haeseler, T. Dechat, R. Foisner, A-type lamins bind both hetero- and euchromatin, the latter being regulated by lamina-associated polypeptide 2 alpha. Genome Res 26, 462–473 (2016). DOI: 10.1101/gr.196220.115.

62. T. Kohwi-Shigematsu, I. deBelle, L. A. Dickinson, S. Galande, Y. Kohwi, Identification of base-unpairing region-binding proteins and characterization of their in vivo binding sequences. Methods Cell Biol 53, 323–354 (1998). DOI:

63. E. A. Oprzeska-Zingrebe, J. Smiatek, Preferential Binding of Urea to Single-Stranded DNA Structures: A Molecular Dynamics Study. Biophys J 114, 1551–1562 (2018). DOI: 10.1016/j.bpj.2018.02.013.

64. A. Das, C. Mukhopadhyay, Urea-mediated protein denaturation: a consensus view. J Phys Chem B 113, 12816–12824 (2009). DOI: 10.1021/jp906350s.

65. S. Raghunathan, T. Jaganade, U. D. Priyakumar, Urea-aromatic interactions in biology. Biophys Rev 12, 65–84 (2020). DOI: 10.1007/s12551-020-00620-9.

66. T. Kohwi-Shigematsu, Y. Kohwi, K. Takahashi, H. W. Richards, S. D. Ayers, H. J. Han, S. Cai, SATB1-mediated functional packaging of chromatin into loops. Methods 58, 243–254 (2012). DOI: 10.1016/j.ymeth.2012.06.019.

67. P. Kourilsky, S. Manteuil, M. H. Zamansky, F. Gros, DNA-RNA hybridization at low temperature in the presence of urea. Biochem Biophys Res Commun 41, 1080–1087 (1970). DOI: 10.1016/0006-291x(70)90196-8.

68. P. Kourilsky, J. Leidner, G. Y. Tremblay, DNA-DNA hybridization on filters at low temperature in the presence of formamide or urea. Biochimie 53, 1111–1114 (1971). DOI: 10.1016/s0300-9084(71)80201-8.

69. E. Hegedus, E. Kokai, A. Kotlyar, V. Dombradi, G. Szabo, Separation of 1-23-kb complementary DNA strands by urea-agarose gel electrophoresis. Nucleic Acids Res 37, e112 (2009). DOI: 10.1093/nar/gkp539.

70. H. Summer, R. Gramer, P. Droge, Denaturing urea polyacrylamide gel electrophoresis (Urea PAGE). J Vis Exp, (2009). DOI: 10.3791/1485.

71. W. Mu, E. S. Davis, S. Lee, M. G. Dozmorov, D. H. Phanstiel, M. I. Love, bootRanges: flexible generation of null sets of genomic ranges for hypothesis testing. Bioinformatics 39, (2023). DOI: 10.1093/bioinformatics/btad190.

72. C. A. Davis, B. C. Hitz, C. A. Sloan, E. T. Chan, J. M. Davidson, I. Gabdank, J. A. Hilton, K. Jain, U. K. Baymuradov, A. K. Narayanan, K. C. Onate, K. Graham, S. R. Miyasato, T. R. Dreszer, J. S. Strattan, O. Jolanki, F. Y. Tanaka, J. M. Cherry, The Encyclopedia of DNA elements (ENCODE): data portal update. Nucleic Acids Res 46, D794–D801 (2018). DOI: 10.1093/nar/gkx1081.

73. J. C. Peng, A. Valouev, T. Swigut, J. Zhang, Y. Zhao, A. Sidow, J. Wysocka, Jarid2/Jumonji coordinates control of PRC2 enzymatic activity and target gene occupancy in pluripotent cells. Cell 139, 1290–1302 (2009). DOI: 10.1016/j.cell.2009.12.002.

74. Y. Tanaka, T. Sotome, A. Inoue, T. Mukozu, T. Kuwabara, T. Mikami, T. Kowhi-Shigematsu, M. Kondo, SATB1 Conditional Knockout Results in Sjogren’s Syndrome in Mice. J Immunol 199, 4016–4022 (2017). DOI: 10.4049/jimmunol.1700550.

75. M. Kondo, Y. Tanaka, T. Kuwabara, T. Naito, T. Kohwi-Shigematsu, A. Watanabe, SATB1 Plays a Critical Role in Establishment of Immune Tolerance. J Immunol 196, 563–572 (2016). DOI: 10.4049/jimmunol.1501429.

76. N. Yannoutsos, V. Barreto, Z. Misulovin, A. Gazumyan, W. Yu, N. Rajewsky, B. R. Peixoto, T. Eisenreich, M. C. Nussenzweig, A cis element in the recombination activating gene locus regulates gene expression by counteracting a distant silencer. Nat Immunol 5, 443–450 (2004). DOI: 10.1038/ni1053.

77. Y. Ben Zouari, A. Platania, A. M. Molitor, T. Sexton, 4See: A Flexible Browser to Explore 4C Data. Front Genet 10, 1372 (2019). DOI: 10.3389/fgene.2019.01372.

78. E. de Wit, B. A. Bouwman, Y. Zhu, P. Klous, E. Splinter, M. J. Verstegen, P. H. Krijger, N. Festuccia, E. P. Nora, M. Welling, E. Heard, N. Geijsen, R. A. Poot, I. Chambers, W. de Laat, The pluripotent genome in three dimensions is shaped around pluripotency factors. Nature 501, 227–231 (2013). DOI: 10.1038/nature12420.

79. G. Geeven, H. Teunissen, W. de Laat, E. de Wit, peakC: a flexible, non-parametric peak calling package for 4C and Capture-C data. Nucleic Acids Res 46, e91 (2018). DOI: 10.1093/nar/gky443.

80. P. H. L. Krijger, G. Geeven, V. Bianchi, C. R. E. Hilvering, W. de Laat, 4C-seq from beginning to end: A detailed protocol for sample preparation and data analysis. Methods 170, 17–32 (2020). DOI: 10.1016/j.ymeth.2019.07.014.

81. D. Noordermeer, M. Leleu, E. Splinter, J. Rougemont, W. De Laat, D. Duboule, The dynamic architecture of Hox gene clusters. Science 334, 222–225 (2011). DOI: 10.1126/science.1207194.

82. P. J. Wijchers, P. H. L. Krijger, G. Geeven, Y. Zhu, A. Denker, M. Verstegen, C. Valdes-Quezada, C. Vermeulen, M. Janssen, H. Teunissen, L. C. M. Anink-Groenen, P. J. Verschure, W. de Laat, Cause and Consequence of Tethering a SubTAD to Different Nuclear Compartments. Mol Cell 61, 461–473 (2016). DOI: 10.1016/j.molcel.2016.01.001.

83. F. Ramirez, D. P. Ryan, B. Gruning, V. Bhardwaj, F. Kilpert, A. S. Richter, S. Heyne, F. Dundar, T. Manke, deepTools2: a next generation web server for deep-sequencing data analysis. Nucleic Acids Res 44, W160–165 (2016). DOI: 10.1093/nar/gkw257.

84. L. A. Dickinson, T. Kohwi-Shigematsu, Nucleolin is a matrix attachment region DNA-binding protein that specifically recognizes a region with high base-unpairing potential. Mol Cell Biol 15, 456–465 (1995). DOI: 10.1128/MCB.15.1.456.

85. W. Meuleman, D. Peric-Hupkes, J. Kind, J. B. Beaudry, L. Pagie, M. Kellis, M. Reinders, L. Wessels, B. van Steensel, Constitutive nuclear lamina-genome interactions are highly conserved and associated with A/T-rich sequence. Genome Res 23, 270–280 (2013). DOI: 10.1101/gr.141028.112.

86. S. Chen, T. R. Luperchio, X. Wong, E. B. Doan, A. T. Byrd, K. Roy Choudhury, K. L. Reddy, M. S. Krangel, A Lamina-Associated Domain Border Governs Nuclear Lamina Interactions, Transcription, and Recombination of the Tcrb Locus. Cell Rep 25, 1729–1740 e1726 (2018). DOI: 10.1016/j.celrep.2018.10.052.

87. M. J. Vogel, D. Peric-Hupkes, B. van Steensel, Detection of in vivo protein-DNA interactions using DamID in mammalian cells. Nat Protoc 2, 1467–1478 (2007). DOI: 10.1038/nprot.2007.148.

88. K. L. Reddy, X. Wong, An Optimized Adaptation of DamID for NGS Applications. Methods Mol Biol 2866, 245–262 (2025). DOI: 10.1007/978-1-0716-4192-7_14.

89. B. Wen, H. Wu, Y. H. Loh, E. Briem, G. Q. Daley, A. P. Feinberg, Euchromatin islands in large heterochromatin domains are enriched for CTCF binding and differentially DNA-methylated regions. BMC Genomics 13, 566 (2012). DOI: 10.1186/1471-2164-13-566.

90. C. Leemans, M. C. H. van der Zwalm, L. Brueckner, F. Comoglio, T. van Schaik, L. Pagie, J. van Arensbergen, B. van Steensel, Promoter-Intrinsic and Local Chromatin Features Determine Gene Repression in LADs. Cell 177, 852–864 e814 (2019). DOI: 10.1016/j.cell.2019.03.009.

91. T. R. Luperchio, Sauria, M. EG., Wong, X., Gaillard, M-C., Tsang, P., Pekrun, K., Ach, R.A., Yamada, N.A., Taylor, J., Reddy, K.L., Chromosome Conformation Paints Reveal the Role of Lamina Association in Genome Organization and Regulation. bioRxiv. DOI: 10.1101/122226.

92. S. Galande, L. A. Dickinson, I. S. Mian, M. Sikorska, T. Kohwi-Shigematsu, SATB1 cleavage by caspase 6 disrupts PDZ domain-mediated dimerization, causing detachment from chromatin early in T-cell apoptosis. Mol Cell Biol 21, 5591–5604. (2001). DOI:

93. L. A. Dickinson, C. D. Dickinson, T. Kohwi-Shigematsu, An atypical homeodomain in SATB1 promotes specific recognition of the key structural element in a matrix attachment region. J Biol Chem 272, 11463–11470. (1997). DOI:

94. K. Nakagomi, Y. Kohwi, L. A. Dickinson, T. Kohwi-Shigematsu, A novel DNA-binding motif in the nuclear matrix attachment DNA-binding protein SATB1. Mol Cell Biol 14, 1852–1860. (1994). DOI:

95. Z. Wang, X. Yang, S. Guo, Y. Yang, X. C. Su, Y. Shen, J. Long, Crystal structure of the ubiquitin-like domain-CUT repeat-like tandem of special AT-rich sequence binding protein 1 (SATB1) reveals a coordinating DNA-binding mechanism. J Biol Chem 289, 27376–27385 (2014). DOI: 10.1074/jbc.M114.562314.

96. P. Pavan Kumar, P. K. Purbey, C. K. Sinha, D. Notani, A. Limaye, R. S. Jayani, S. Galande, Phosphorylation of SATB1, a global gene regulator, acts as a molecular switch regulating its transcriptional activity in vivo. Mol Cell 22, 231–243 (2006). DOI:

97. D. Notani, K. P. Gottimukkala, R. S. Jayani, A. S. Limaye, M. V. Damle, S. Mehta, P. K. Purbey, J. Joseph, S. Galande, Global regulator SATB1 recruits beta-catenin and regulates T(H)2 differentiation in Wnt-dependent manner. PLoS Biol 8, e1000296 (2010). DOI: 10.1371/journal.pbio.1000296.

98. D. Notani, P. L. Ramanujam, P. P. Kumar, K. P. Gottimukkala, C. Kumar-Sinha, S. Galande, N-terminal PDZ-like domain of chromatin organizer SATB1 contributes towards its function as transcription regulator. J Biosci 36, 461–469 (2011). DOI: 10.1007/s12038-011-9091-4.

99. T. L. Stephen, K. K. Payne, R. A. Chaurio, M. J. Allegrezza, H. Zhu, J. Perez-Sanz, A. Perales-Puchalt, J. M. Nguyen, A. E. Vara-Ailor, E. B. Eruslanov, M. E. Borowsky, R. Zhang, T. M. Laufer, J. R. Conejo-Garcia, SATB1 Expression Governs Epigenetic Repression of PD-1 in Tumor-Reactive T Cells. Immunity 46, 51–64 (2017). DOI: 10.1016/j.immuni.2016.12.015.

100. D. Yasui, M. Miyano, S. Cai, P. Varga-Weisz, T. Kohwi-Shigematsu, SATB1 targets chromatin remodelling to regulate genes over long distances. Nature 419, 641–645. (2002). DOI:

101. T. Zhan, N. Rindtorff, M. Boutros, Wnt signaling in cancer. Oncogene 36, 1461–1473 (2017). DOI: 10.1038/onc.2016.304.

102. P. Song, Z. Gao, Y. Bao, L. Chen, Y. Huang, Y. Liu, Q. Dong, X. Wei, Wnt/beta-catenin signaling pathway in carcinogenesis and cancer therapy. Journal of hematology & oncology 17, 46 (2024). DOI: 10.1186/s13045-024-01563-4.

103. J. A. Nickerson, G. Krochmalnic, K. M. Wan, S. Penman, Chromatin architecture and nuclear RNA. Proc Natl Acad Sci U S A 86, 177–181 (1989). DOI: 10.1073/pnas.86.1.177.

104. J. A. Nickerson, The ribonucleoprotein network of the nucleus: a historical perspective. Curr Opin Genet Dev 75, 101940 (2022). DOI: 10.1016/j.gde.2022.101940.

105. J. Thakur, S. Henikoff, Architectural RNA in chromatin organization. Biochem Soc Trans 48, 1967–1978 (2020). DOI: 10.1042/BST20191226.

106. D. Michieletto, N. Gilbert, Role of nuclear RNA in regulating chromatin structure and transcription. Curr Opin Cell Biol 58, 120–125 (2019). DOI: 10.1016/j.ceb.2019.03.007.

107. D. Peric-Hupkes, W. Meuleman, L. Pagie, S. W. Bruggeman, I. Solovei, W. Brugman, S. Graf, P. Flicek, R. M. Kerkhoven, M. van Lohuizen, M. Reinders, L. Wessels, B. van Steensel, Molecular maps of the reorganization of genome-nuclear lamina interactions during differentiation. Mol Cell 38, 603–613 (2010). DOI: 10.1016/j.molcel.2010.03.016.

108. H. Zhao, E. G. Sifakis, N. Sumida, L. Millan-Arino, B. A. Scholz, J. P. Svensson, X. Chen, A. L. Ronnegren, C. D. Mallet de Lima, F. S. Varnoosfaderani, C. Shi, O. Loseva, S. Yammine, M. Israelsson, L. S. Rathje, B. Nemeti, E. Fredlund, T. Helleday, M. P. Imreh, A. Gondor, PARP1- and CTCF-Mediated Interactions between Active and Repressed Chromatin at the Lamina Promote Oscillating Transcription. Mol Cell 59, 984–997 (2015). DOI: 10.1016/j.molcel.2015.07.019.

109. T. Lucas, T. L. Hafer, H. G. Zhang, N. Molotkova, M. Kohwi, Discrete cis-acting element regulates developmentally timed gene-lamina relocation and neural progenitor competence in vivo. Dev Cell 56, 2649–2663 e2646 (2021). DOI: 10.1016/j.devcel.2021.08.020.

110. I. Solovei, M. Kreysing, C. Lanctot, S. Kosem, L. Peichl, T. Cremer, J. Guck, B. Joffe, Nuclear architecture of rod photoreceptor cells adapts to vision in mammalian evolution. Cell 137, 356–368 (2009). DOI: 10.1016/j.cell.2009.01.052.

111. J. Kind, L. Pagie, H. Ortabozkoyun, S. Boyle, S. S. de Vries, H. Janssen, M. Amendola, L. D. Nolen, W. A. Bickmore, B. van Steensel, Single-cell dynamics of genome-nuclear lamina interactions. Cell 153, 178–192 (2013). DOI: 10.1016/j.cell.2013.02.028.

112. Q. Bian, N. Khanna, J. Alvikas, A. S. Belmont, beta-Globin cis-elements determine differential nuclear targeting through epigenetic modifications. J Cell Biol 203, 767–783 (2013). DOI: 10.1083/jcb.201305027.

113. X. Wong, T. R. Luperchio, K. L. Reddy, NET gains and losses: the role of changing nuclear envelope proteomes in genome regulation. Curr Opin Cell Biol 28, 105–120 (2014). DOI: 10.1016/j.ceb.2014.04.005.

114. J. C. Harr, T. R. Luperchio, X. Wong, E. Cohen, S. J. Wheelan, K. L. Reddy, Directed targeting of chromatin to the nuclear lamina is mediated by chromatin state and A-type lamins. J Cell Biol 208, 33–52 (2015). DOI: 10.1083/jcb.201405110.

115. X. Wong, V. E. Hoskins, A. J. Melendez-Perez, J. C. Harr, M. Gordon, K. L. Reddy, Lamin C is required to establish genome organization after mitosis. Genome Biol 22, 305 (2021). DOI: 10.1186/s13059-021-02516-7.

116. L. A. Watson, L. H. Tsai, In the loop: how chromatin topology links genome structure to function in mechanisms underlying learning and memory. Curr Opin Neurobiol 43, 48–55 (2017). DOI: 10.1016/j.conb.2016.12.002.

117. J. Close, H. Xu, N. De Marco Garcia, R. Batista-Brito, E. Rossignol, B. Rudy, G. Fishell, Satb1 is an activity-modulated transcription factor required for the terminal differentiation and connectivity of medial ganglionic eminence-derived cortical interneurons. J Neurosci 32, 17690–17705 (2012). DOI: 10.1523/JNEUROSCI.3583-12.2012.

118. S. Galande, T. Kohwi-Shigematsu, Poly(ADP-ribose) polymerase and Ku autoantigen form a complex and synergistically bind to matrix attachment sequences. J Biol Chem 274, 20521–20528. (1999). DOI:

119. W. M. Liu, F. K. Guerra-Vladusic, S. Kurakata, R. Lupu, T. Kohwi-Shigematsu, HMG-I(Y) recognizes base-unpairing regions of matrix attachment sequences and its increased expression is directly linked to metastatic breast cancer phenotype. Cancer Res 59, 5695–5703. (1999). DOI:

120. M. Levo, J. Raimundo, X. Y. Bing, Z. Sisco, P. J. Batut, S. Ryabichko, T. Gregor, M. S. Levine, Transcriptional coupling of distant regulatory genes in living embryos. Nature 605, 754–760 (2022). DOI: 10.1038/s41586-022-04680-7.

121. P. J. Batut, X. Y. Bing, Z. Sisco, J. Raimundo, M. Levo, M. S. Levine, Genome organization controls transcriptional dynamics during development. Science 375, 566–570 (2022). DOI: 10.1126/science.abi7178.

122. G. Mohana, J. Dorier, X. Li, M. Mouginot, R. C. Smith, H. Malek, M. Leleu, D. Rodriguez, J. Khadka, P. Rosa, P. Cousin, C. Iseli, S. Restrepo, N. Guex, B. D. McCabe, A. Jankowski, M. S. Levine, M. C. Gambetta, Chromosome-level organization of the regulatory genome in the Drosophila nervous system. Cell 186, 3826–3844 e3826 (2023). DOI: 10.1016/j.cell.2023.07.008.

123. L. Lizana, Y. B. Schwartz, The scales, mechanisms, and dynamics of the genome architecture. Sci Adv 10, eadm8167 (2024). DOI: 10.1126/sciadv.adm8167.

124. W. Huber, V. J. Carey, R. Gentleman, S. Anders, M. Carlson, B. S. Carvalho, H. C. Bravo, S. Davis, L. Gatto, T. Girke, R. Gottardo, F. Hahne, K. D. Hansen, R. A. Irizarry, M. Lawrence, M. I. Love, J. MacDonald, V. Obenchain, A. K. Oles, H. Pages, A. Reyes, P. Shannon, G. K. Smyth, D. Tenenbaum, L. Waldron, M. Morgan, Orchestrating high-throughput genomic analysis with Bioconductor. Nat Methods 12, 115–121 (2015). DOI: 10.1038/nmeth.3252.

125. A. B. Olshen, E. S. Venkatraman, R. Lucito, M. Wigler, Circular binary segmentation for the analysis of array-based DNA copy number data. Biostatistics 5, 557–572 (2004). DOI: 10.1093/biostatistics/kxh008.

126. B. Gel, A. Diez-Villanueva, E. Serra, M. Buschbeck, M. A. Peinado, R. Malinverni, regioneR: an R/Bioconductor package for the association analysis of genomic regions based on permutation tests. Bioinformatics 32, 289–291 (2016). DOI: 10.1093/bioinformatics/btv562.

127. Z. Gu, R. Eils, M. Schlesner, N. Ishaque, EnrichedHeatmap: an R/Bioconductor package for comprehensive visualization of genomic signal associations. BMC Genomics 19, 234 (2018). DOI: 10.1186/s12864-018-4625-x.

128. Z. Gu, L. Gu, R. Eils, M. Schlesner, B. Brors, circlize Implements and enhances circular visualization in R. Bioinformatics 30, 2811–2812 (2014). DOI: 10.1093/bioinformatics/btu393.

129. C. Lazaris, S. Kelly, P. Ntziachristos, I. Aifantis, A. Tsirigos, HiC-bench: comprehensive and reproducible Hi-C data analysis designed for parameter exploration and benchmarking. BMC Genomics 18, 22 (2017). DOI: 10.1186/s12864-016-3387-6.

130. A. Kloetgen, P. Thandapani, P. Ntziachristos, Y. Ghebrechristos, S. Nomikou, C. Lazaris, X. Chen, H. Hu, S. Bakogianni, J. Wang, Y. Fu, F. Boccalatte, H. Zhong, E. Paietta, T. Trimarchi, Y. Zhu, P. Van Vlierberghe, G. G. Inghirami, T. Lionnet, I. Aifantis, A. Tsirigos, Three-dimensional chromatin landscapes in T cell acute lymphoblastic leukemia. Nat Genet 52, 388–400 (2020). DOI: 10.1038/s41588-020-0602-9.

131. B. Langmead, S. L. Salzberg, Fast gapped-read alignment with Bowtie 2. Nat Methods 9, 357–359 (2012). DOI: 10.1038/nmeth.1923.

132. A. Tsirigos, N. Haiminen, E. Bilal, F. Utro, GenomicTools: a computational platform for developing high-throughput analytics in genomics. Bioinformatics 28, 282–283 (2012). DOI: 10.1093/bioinformatics/btr646.

133. M. Imakaev, G. Fudenberg, R. P. McCord, N. Naumova, A. Goloborodko, B. R. Lajoie, J. Dekker, L. A. Mirny, Iterative correction of Hi-C data reveals hallmarks of chromosome organization. Nat Methods 9, 999–1003 (2012). DOI: 10.1038/nmeth.2148.

134. Y. Gong, C. Lazaris, T. Sakellaropoulos, A. Lozano, P. Kambadur, P. Ntziachristos, I. Aifantis, A. Tsirigos, Stratification of TAD boundaries reveals preferential insulation of super-enhancers by strong boundaries. Nat Commun 9, 542 (2018). DOI: 10.1038/s41467-018-03017-1.

135. N. C. Durand, M. S. Shamim, I. Machol, S. S. Rao, M. H. Huntley, E. S. Lander, E. L. Aiden, Juicer Provides a One-Click System for Analyzing Loop-Resolution Hi-C Experiments. Cell Syst 3, 95–98 (2016). DOI: 10.1016/j.cels.2016.07.002.

136. S. Heinz, C. Benner, N. Spann, E. Bertolino, Y. C. Lin, P. Laslo, J. X. Cheng, C. Murre, H. Singh, C. K. Glass, Simple combinations of lineage-determining transcription factors prime cis-regulatory elements required for macrophage and B cell identities. Mol Cell 38, 576–589 (2010). DOI: 10.1016/j.molcel.2010.05.004.

137. Y. Zhang, T. Liu, C. A. Meyer, J. Eeckhoute, D. S. Johnson, B. E. Bernstein, C. Nusbaum, R. M. Myers, M. Brown, W. Li, X. S. Liu, Model-based analysis of ChIP-Seq (MACS). Genome Biol 9, R137 (2008). DOI: gb-2008-9-9-r137 [pii] 10.1186/gb-2008-9-9-r137.

138. W. J. Kent, A. S. Zweig, G. Barber, A. S. Hinrichs, D. Karolchik, BigWig and BigBed: enabling browsing of large distributed datasets. Bioinformatics 26, 2204–2207 (2010). DOI: 10.1093/bioinformatics/btq351.

139. J. M. Zullo, I. A. Demarco, R. Pique-Regi, D. J. Gaffney, C. B. Epstein, C. J. Spooner, T. R. Luperchio, B. E. Bernstein, J. K. Pritchard, K. L. Reddy, H. Singh, DNA sequence-dependent compartmentalization and silencing of chromatin at the nuclear lamina. Cell 149, 1474–1487 (2012). DOI: 10.1016/j.cell.2012.04.035.

140. A. Favorov, L. Mularoni, L. M. Cope, Y. Medvedeva, A. A. Mironov, V. J. Makeev, S. J. Wheelan, Exploring massive, genome scale datasets with the GenometriCorr package. PLoS Comput Biol 8, e1002529 (2012). DOI: 10.1371/journal.pcbi.1002529.

141. A. Akalin, V. Franke, K. Vlahovicek, C. E. Mason, D. Schubeler, Genomation: a toolkit to summarize, annotate and visualize genomic intervals. Bioinformatics 31, 1127–1129 (2015). DOI: 10.1093/bioinformatics/btu775.

142. A. R. Quinlan, I. M. Hall, BEDTools: a flexible suite of utilities for comparing genomic features. Bioinformatics 26, 841–842 (2010). DOI: 10.1093/bioinformatics/btq033.

